# Circumpolar peoples and their languages: lexical and genomic data suggest ancient Chukotko-Kamchatkan–Nivkh and Yukaghir-Samoyedic connections

**DOI:** 10.1101/2021.02.27.433193

**Authors:** George Starostin, N. Ezgi Altınışık, Mikhail Zhivlov, Piya Changmai, Olga Flegontova, Sergey A. Spirin, Andrei Zavgorodnii, Pavel Flegontov, Alexei S. Kassian

## Abstract

Relationships between universally recognized language families represent a hotly debated topic in historical linguistics, and the same is true for correlation between signals of genetic and linguistic relatedness. We developed a weighted permutation test and applied it on basic vocabularies for 31 pairs of languages and reconstructed proto-languages to show that three groups of circumpolar language families in the Northern Hemisphere show evidence of relationship though borrowing in the basic vocabulary or common descent: [Chukotko-Kamchatkan and Nivkh]; [Yukaghir and Samoyedic]; [Yeniseian, Na-Dene, and Burushaski]. The former two pairs showed the most significant signals of language relationship, and the same pairs demonstrated parallel signals of genetic relationship implying common descent or substantial gene flows. For finding the genetic signals we used genome-wide genetic data for present-day groups and a bootstrapping model comparison approach for admixture graphs or, alternatively, haplotype sharing statistics. Our findings further support some hypotheses on long-distance language relationship put forward based on the linguistic methods but lacking universal acceptance.

**Significance statement:** Indigenous people inhabiting polar and sub-polar regions in the Northern Hemisphere speak diverse languages belonging to at least seven language families which are traditionally thought of as unrelated entities. We developed a weighted permutation test and applied it to basic vocabularies of a number of languages and reconstructed proto-languages to show that at least three groups of circumpolar language families show evidence of relationship though either borrowing in the basic vocabulary or common descent: Chukotko-Kamchatkan and Nivkh; Yukaghir and Samoyedic; Yeniseian, Na-Dene, and Burushaski. The former two pairs showed the most significant signals of language relationship, and the same pairs demonstrated parallel signals of genetic relationship implying common descent or substantial gene flows.

## Introduction

There is a large number of populations speaking an impressive variety of diverse languages around the Arctic Circle in Northern Europe, Siberia, Alaska, the Canadian Arctic, and Greenland. Some of these populations are recent newcomers, e.g., Russians, others are thought to have arrived at least several thousand years before present, e.g., Chukchi and Itelmens. As for their languages, those that are more recent in the area usually belong to well-studied language families, such as Turkic and Indo-European, while many others are classified as isolates (Nivkh, Haida). Some of these small linguistic groups and dispersed isolates have been traditionally labeled as Paleo-Asiatic (or Paleo-Siberian) languages; however, there is no evidence for considering Paleo-Asiatic a valid historical clade, and the term is usually understood to represent a wide contact area affected by multiple migrations in historical and prehistorical epochs. The goal of the present study is to examine whether one can find evidence for prehistoric language relationship and/or language contacts in the Circumpolar area and whether the results of this linguistic examination are in agreement with reconstructions of population history.

A special related linguistic issue that is also addressed in the present paper is the external connections of the Yeniseian language family. As of now, there are two main views on the problem of Yeniseian linguistic ancestry. According to the first one, the closest linguistic relative of the Yeniseian family is the Na-Dene family in North America (1). This was first explicitly proposed by Ruhlen (2) and is currently being extensively promoted by Vajda (3–5). This hypothesis has found some support in linguistic circles (6), but many remain skeptical (7), and evidence for a particularly close relationship between the two families seems to be insufficient at the current stage of research (8, 9). The second, earlier, hypothesis, proposed by S. Starostin (10, 11), defines Yeniseian as a member of the hypothetical Sino-Caucasian macrofamily (a genealogical linguistic unity of substantial time depth which at its current maximum extent comprises the North Caucasian, Basque, Yeniseian, Burushaski, Sino-Tibetan and Na-Dene families, see (12) for a short overview). According to S. Starostin’s view, later elaborated by G. Starostin (8, 9), the corresponding fragment of the Sino-Caucasian macrofamily has the following structure: [[Yeniseian, Burushaski], Na-Dene], meaning that the Yeniseian and Na-Dene families are indeed distantly related, but Burushaski is considered the closest relative of the Yeniseian family.

In this study, we explored population relationship of Circumpolar peoples (Fig. 1) in an interdisciplinary way utilizing linguistic and genetic methods. At the moment there is little consensus regarding major events in the population history of Siberia and the American Arctic. The relationships of populations forming prehistoric migration waves in Siberia to present-day Siberians are not settled (1, 13), as well as the sources and number of admixture events between Siberians and First Americans (1, 13–15). In this study we used sensitive methods of genetic analysis for checking if there are parallels between genetic and linguistic history in the Circumpolar region: a) a model-comparison approach for admixture graphs of arbitrary complexity and an admixture graph for Siberians introduced by Ning et al. (14); b) a haplotype-sharing statistic introduced by Flegontov et al. (1).

**Fig. 1.**
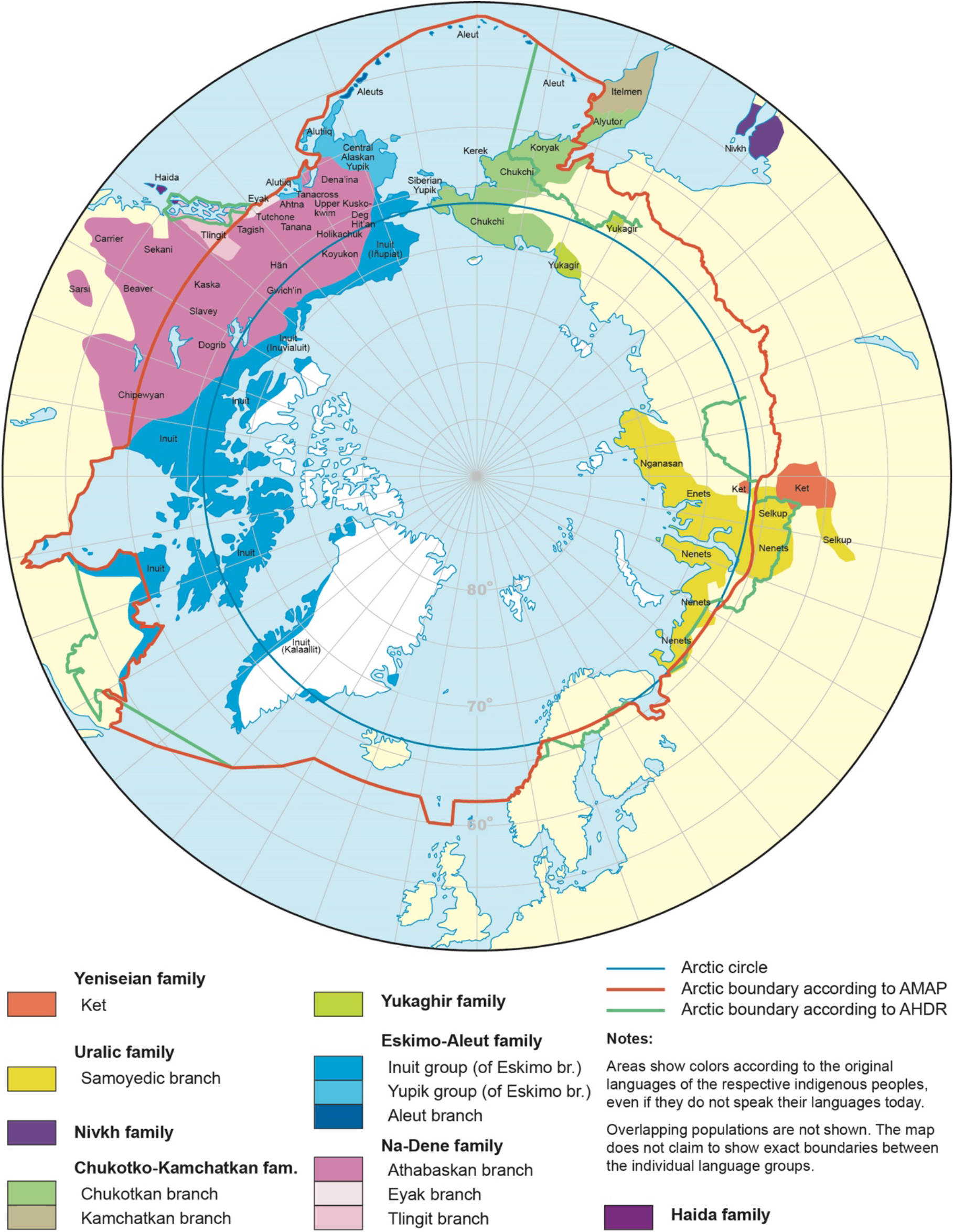
Present-day Circumpolar peoples subdivided according to language families analyzed in the present paper. The figure is based on the map by Winfried K. Dallmann (Norwegian Polar Institute) and Peter Schweitzer (University of Alaska Fairbanks), https://arctic-council.org/index.php/en/learn-more/maps.

## Results

### Linguistic Findings

Probabilistic results of the pairwise comparison of 8 Circumpolar wordlists are presented in Table 1. To cope with the multiple testing problem, we have applied the Holm-Bonferroni correction (16) with α=0.05 and found two pairs with statistically significant p-values: Chukotko-Kamchatkan– Nivkh (p=4.2×10^−5^) and Samoyedic-Yukaghir (p=5.12×10^−4^). The Chukotko-Kamchatkan–Nivkh p-values remains significant even at the α=0.003 (three sigma) significance threshold. See Tables S5-S6 for lexical matches in the aforementioned pairs of wordlists.

**Table 1.** Probabilities of phonetic matches between Circumpolar languages obtained by the weighted permutation test. Statistically significant values under the Holm-Bonferroni correction are shadowed in green (α=0.003) or yellow (α=0.05).

|  | Proto-Athabaskan-Eyak + Tlingit | Haida | Proto-Eskimo + Proto-Aleut | Proto-Chukotian + Proto-Itelmen | Proto-Nivkh | Proto-Samoyedic | Proto-Yukaghir |
| --- | --- | --- | --- | --- | --- | --- | --- |
| Proto-Yeniseian | 0.014 | 0.221 | 0.151 | 0.34 | 0.116 | 0.445 | 0.469 |
| Proto-Athabaskan-Eyak + Tlingit | — | 0.997 | 0.01 | 0.474 | 0.124 | 0.557 | 0.466 |
| Haida |  | — | 0.094 | 0.262 | 0.233 | 0.577 | 0.615 |
| Proto-Eskimo + Proto-Aleut |  |  | — | 0.042 | 0.303 | 0.05 | 0.137 |
| Proto-Chukotian + Proto-Itelmen |  |  |  | — | **$4.2 \times 10^{-5}$** | 0.059 | 0.455 |
| Proto-Nivkh |  |  |  |  | — | 0.035 | 0.128 |
| Proto-Samoyedic |  |  |  |  |  | — | **$5.12 \times 10^{-4}$** |

Probabilistic results of the Yeniseian–Na-Dene–Burushaski comparison are presented in Table 2 (more detailed information is offered in Suppl. Text Table S4). Since S. Starostin’s Yeniseian-Burushaski connection and Ruhlen & Vajda’s Yeniseian–Na-Dene connection are two independent hypotheses proposed by preceding authors on the basis of their own methodologies, there is no need to introduce correction for multiple testing. The p-value for the Yeniseian-Burushaski pair is significant at α=0.01; the p-value for the Yeniseian–Na-Dene pair is significant at α=0.05.

**Table. 2.**
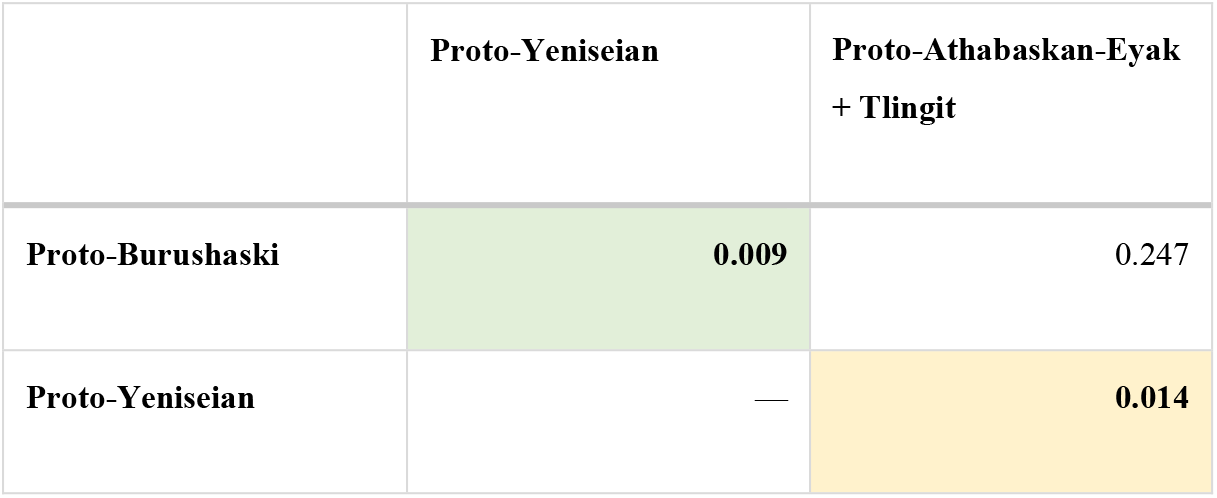
Probabilities of phonetic matches between Yeniseian, Na-Dene and Burushaski obtained by the weighted permutation test. Statistically significant values are shadowed in green (α=0.01) or yellow (α=0.05).

As a positive control, we reran the permutation test for pairs of languages which are genealogically related by consensus (Table 3) and found all the obtained p-values statistically significant (no correction for multiple testing was applied): 0.01 ≤ p < 0.05 for Athabaskan-Tlingit and p < 0.003 for the other pairs.

**Table 3.** Probabilities of phonetic matches between languages generally accepted to be related as obtained by the weighted permutation test.

|  | Eyak | Tlingit | Proto-Aleut | Proto-Itelmen |
| --- | --- | --- | --- | --- |
| **Proto-Athabaskan** | $< 10^{-7}$ | 0.01 | | |
| **Eyak** | | $1.17 \times 10^{-4}$ | | |
| **Proto-Eskimo** | | | $2.02 \times 10^{-3}$ | |
| **Proto-Chukotian** | | | | $< 10^{-7}$ |

### Genetic parallels for linguistic signals

Here we focus on two strongest signals of linguistic relationship found in this study (Samoyedic– Yukaghir and Nivkh–Chukotko-Kamchatkan) and explore signals of close genetic relationship for these pairs of groups. The genetic proximity of the former pair is apparent based on standard methods of autosomal genetic analysis: ADMIXTURE and principal component analysis (1, 17). “Forest Yukaghirs” fall into the West Siberian genetic cluster composed of Samoyedic speakers, Kets, Ugric speakers and Turkic speakers in the Altai region (1), although geographically Forest Yukaghirs are distant from West Siberia.

Using normalized haplotype sharing statistics (HSS) developed by Flegontov et al. (1), we investigated the relationship between Yukaghirs and Samoyedic speakers (represented by Enets, Nganasans, and Selkups, see Suppl. Table 1 for all population annotations). Average normalized HSS for populations are shown in Fig. 2a, and the same statistics for individuals are shown in Fig. 2b. Both Forest and Tundra Yukaghirs are significantly closer to Samoyedic-speaking Selkups and Nganasans than any other Siberians investigated here (after permutation tests all p-values < 0.05, Suppl. Table 2), although Tundra Yukaghirs are genetically heterogeneous, and a sub-group of them is closer to Samoyedic speakers than the rest (Fig. 2b). We conclude that signals of genetic and linguistic relationship are concordant in the case of present-day Samoyedic speakers and Yukaghirs.

**Fig. 2.**
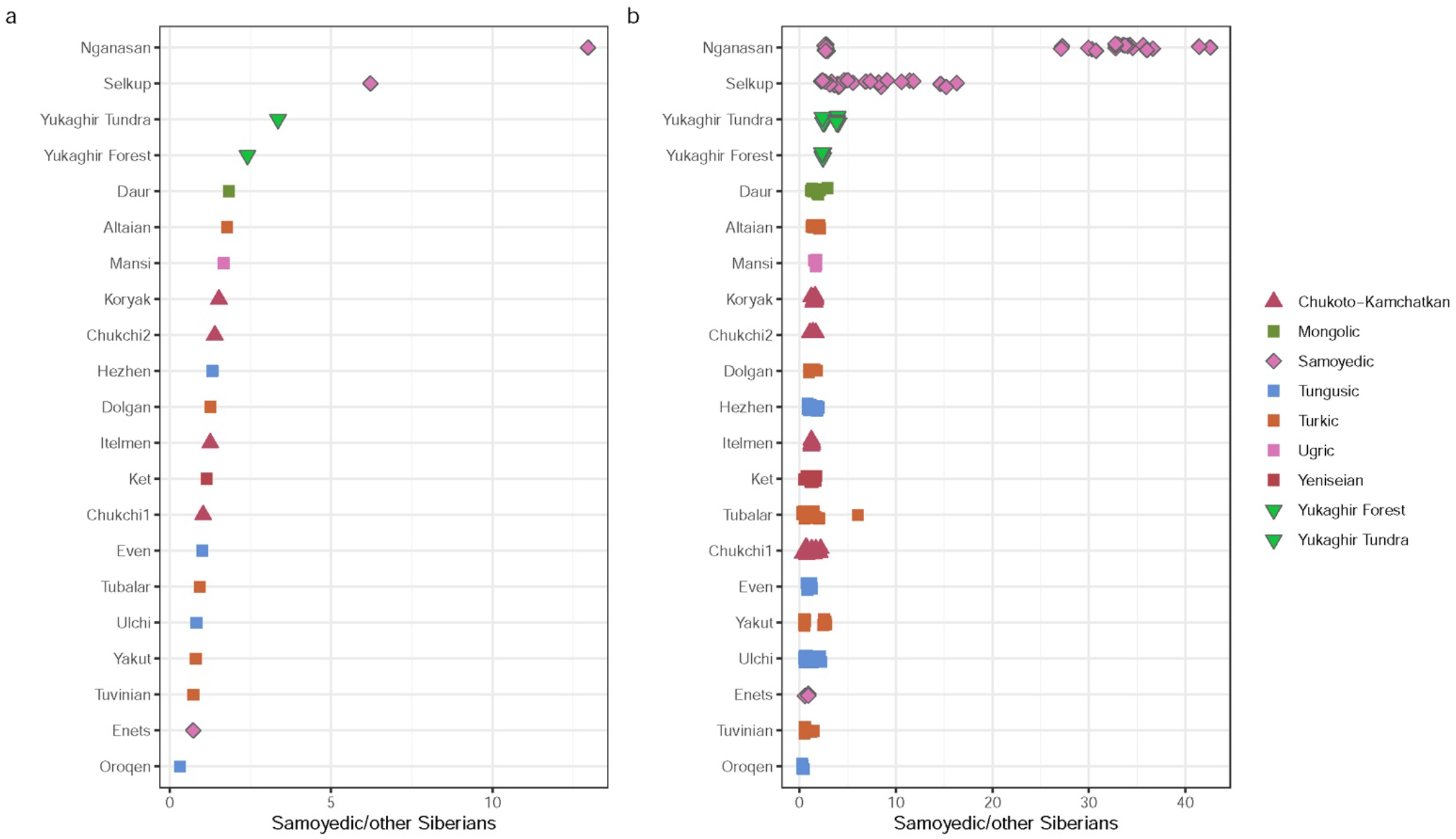
Samoyedic haplotype-sharing statistics normalized by haplotype sharing with other Siberians. Statistics averaged across individuals in groups are shown in panel **a**, and statistics for individuals are shown in panel **b**. When necessary, the statistics for individuals were calculated using a leave-one-out procedure.

The Nivkh–Chukotko-Kamchatkan pair is less straightforward for genetic analysis since these groups are clearly distant genetically as illustrated by principal component analysis (PCA, Fig. 3a) or multidimensional scaling (MDS, based on a matrix of statistics 1 - *f*_*3*_(Yoruba, X, Y)) (Fig. 3b) calculated on a dataset derived from the 1240K SNP panel (18). Those few Nivkh individuals whose genomes were sequenced (19) are genetically similar to Ulchi and other Tungusic speakers, and Chukotko-Kamchatkan speakers (Koryaks, Chukchi, and Itelmen) fall into a distant Paleo-Eskimo-related genetic cluster that was also influenced by a low-level gene flow from America mediated by Eskimo-Aleut speakers (see a further discussion of Arctic population history in (1)). Thus, approaches looking at the genome as a single entity (haplotype sharing and allele sharing statistics) are not expected to reveal genetic proximity of Nivkhs and Chukotko-Kamchatkan speakers, and we looked at methods that allow breaking a genome down into ancestry components.

**Fig. 3.**
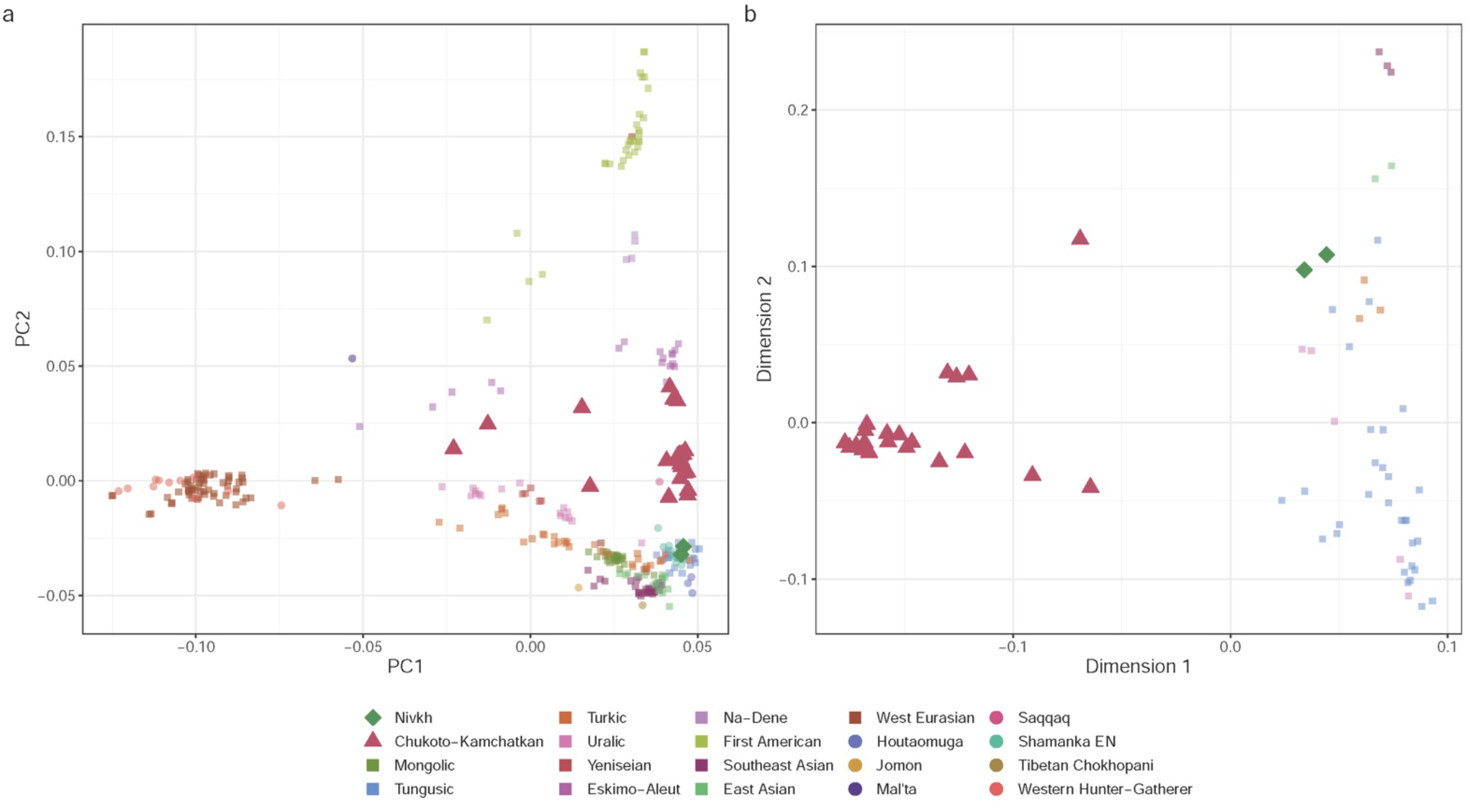
Principal component (**a**) and multidimensional scaling (**b**) analyses illustrating genetic distances between Nivkhs and Chukotko-Kamchatkan speakers. Linguistic and genetic groupings are labeled according to the legend. The plot in panel **b** is based on a matrix of statistics 1 – *f*_*3*_(Yoruba, X, Y), and the plot in panel **a** was constructed using the standard PCA algorithm for SNP genotyping data (72, 73).

We used methods developed by Ning et al. (14). In that study, a graph was constructed modelling divergence and admixture among five Siberian groups (a group from the Houtaomuga site in Northeast China spanning a period between the Paleolithic and Iron Age as a representative of ancient Amur River basin (ARB) populations, “Baikal hunter-gatherers” *sensu* (20), the Saqqaq Paleo-Inuit, Selkups and Nganasans) and other Asian groups (see Suppl. Table 3 for population annotations in this dataset). Here we illustrate that graph showing uncertainty in edge length and admixture weight estimates (Suppl. Fig. 1). A protocol was also developed for admixture inference via “mapping” a target group on a “skeleton” admixture graph as unadmixed, as a two-way and three-way admixture of all possible sources (14). Using the mapping protocol, it was found that Chukotko-Kamchatkan (C-K) speakers can be modelled as having Asian part of their ancestry derived from two lineages: a lineage represented by the Saqqaq Paleo-Inuit individual (21) and a First American lineage (14). That result is in line with an earlier study (1). It was also found that Nivkhs and Ulchi are best modelled as having Asian part of their ancestry derived from three lineages: Amur River basin groups (a bulk of their ancestry), a lineage represented by Nganasans, and a lineage represented by the Ikawazu Jomon (22) individual (14). In that study a bootstrapping method for comparing fits of two arbitrary admixture graph models was also proposed (14). We used the 14-population skeleton graph developed in that study (Suppl. Fig. 1) and a variation of the bootstrapping model comparison algorithm for tracing genetic connections between Nivkhs and Chukotko-Kamchatkan speakers (see Methods for details).

We grafted (Suppl. Fig. 2) all possible combinations of the C-K and Tungusic or Nivkh (TN) groups (and Turkic-speaking Tuvinians as a negative control) on the skeleton graph and allowed no missing SNP data at the group level. For each combination of the groups, the following models were compared: baseline with no gene flows between the C-K and TN groups vs. two models with a unidirectional C-K/TN gene flow, baseline vs. a model with the bidirectional C-K/TN gene flow, the bidirectional model vs. two models with a unidirectional C-K/TN gene flow (five comparisons per population combination).

Across 200 model comparisons performed (with 500 bootstrap iterations each), 10 comparisons demonstrated one-tailed empirical *p*-values < 0.05 (Suppl. Table 4). These significant outcomes of model comparison were non-randomly distributed across population pairs, with five pairs including Evens from Kamchatka (Magadan) and Koryaks or Itelmen from the same region (Suppl. Table 3). Two pairs included Evenks and Koryaks, one pair included Evens from Yakutia and Koryaks, and two pairs included Nivkhs and Koryaks. Thus, models including a C-K/TN gene flow were preferred to the baseline model in two cases: either when the C-K and TN groups reside in relative proximity, or, in the case of Nivkhs and Koryaks, separated by a large distance by land or by sea. The bidirectional model for the Nivkh/Koryak pair and the “C-K => Nivkh” model showed similar *p*-values (0.036 and 0.028, respectively) and similar average log-likelihood (LL) differences vs. the baseline model (88.1 and 85.9 ln-units, respectively) (Suppl. Table 4). Distributions of LL differences across bootstrap replicates for all 10 significant model comparisons are shown in Fig. 4. Since two significant model comparisons for Nivkhs and Koryaks demonstrated nearly identical average LL differences (Fig. 4), the simpler “C-K => Nivkh” model should be preferred, and the comparison of that model to the bidirectional model demonstrated a non-significant *p*-value (0.48, average LL difference = 2.2 ln-units). There was also no statistically significant difference in the fits of the “C-K => Nivkh” and “Nivkh => C-K” models (p-value = 0.33, average LL difference = 20 ln-units).

**Fig. 4.**
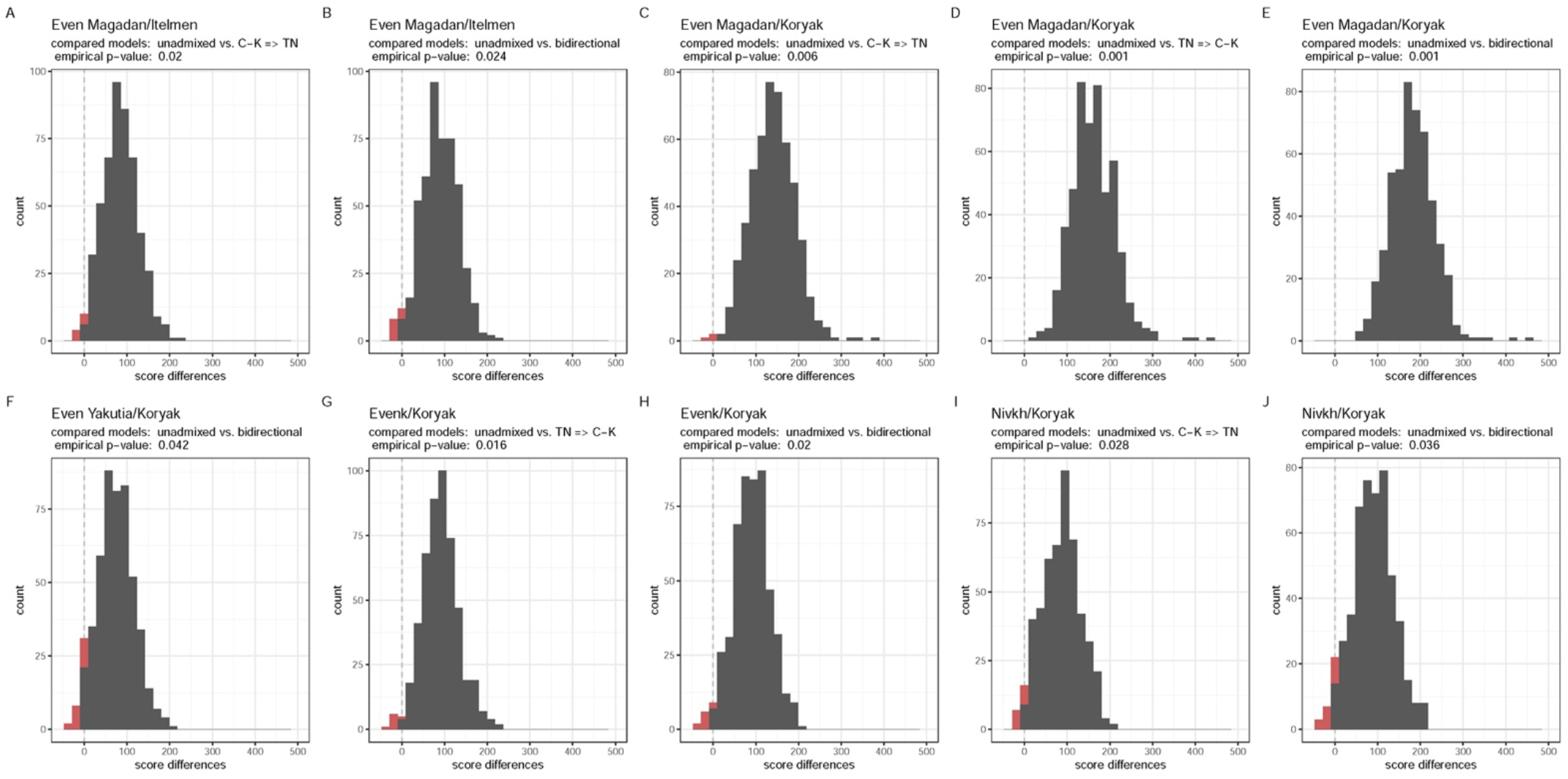
Distributions of log-likelihood differences across bootstrap replicates for admixture graph model comparisons that demonstrated significant one-tailed empirical *p*-values. Group and model labels and *p*-values are shown above each plot. Negative log-likelihood differences are highlighted in red.

In summary, we revealed a statistically supported gene flow from Chukotko-Kamchatkan speakers to Nivkhs and found that from 14% to 21% of Nivkh genomes are derived from Chukotko-Kamchatkan speakers (those are minimal and maximal values encountered across 100 bootstrap replicates of the dataset). Indeed, when the MDS plot based on *f*_*3*_-statistic is inspected (Fig. 3b), we see that Nivkhs are slightly shifted from Ulchi and other Siberians in the direction of Koryaks, Itelmen, and Chukchi. However, we cannot rule out a model where the gene flow happened in the other direction (Nivkh => C-K, from 10% to 15% of Koryak genomes derived from the Nivkh lineage under that model). These genetic signals are again concordant with the linguistic statistical analysis in this study.

## Discussion

The Circumpolar comparison (Table 1) detected two pairs of protolanguages with statistically significant similarity.

The first language pair is Chukotko-Kamchatkan–Nivkh (*p* < 0.003, corrected for multiple testing). To the best of our knowledge, a specific genealogical relationship between the Chukotko-Kamchatkan and Nivkh languages was first proposed by Fortescue (23), although it must be noted that earlier Mudrak and Nikolaev (24), relying on the controversial multilateral or “mass” comparison approach introduced by Greenberg, had suggested that Chukotko-Kamchatkan and Nivkh along with a number of languages of North America may constitute a hypothetical Almosan-Keresiouan macrofamily; later Nikolaev (25) claimed that lexical parallels between Chukotko-Kamchatkan and Nivkh are of contact origin, but did not provide reliable arguments for this contact scenario. It is notable that in his research, Fortescue relied purely on the comparative method traditionally accepted by historical linguists. In other words, both the traditional expert approach and the formal technique employed in the present paper have yielded the same result.

The second language pair is Samoyedic-Yukaghir (0.01 ≤ p < 0.05, corrected for multiple testing). Lexical parallels between Uralic (particularly Samoyedic) and Yukaghir vocabularies are striking and have attracted linguists’ attention for a long time. These can be viewed either as traces of a distant genealogical relationship between the two families (26), or as borrowings, mainly from pre-Proto-Samoyedic into Yukaghir (27). We regard the question of the origin of these parallels as yet unsolved, although some thoughts can be stated.

A scenario that implies a genealogical relationship between Uralic and Yukaghir has two serious shortcomings. First, the majority of Uralic roots with Yukaghir comparanda are either Common Uralic (i.e., attested in both branches of the family: Finno-Ugric and Samoyedic) or restricted to Samoyedic (undocumented for the Finno-Ugric branch). Out of 43 reliable Uralic/Samoyedic-Yukaghir comparisons in (27) only 8 lack a Samoyedic reflex. Moreover, two of these eight words show specific Proto-Samoyedic phonetic development which implies that these stems were present in Proto-Samoyedic, having been later lost in modern Samoyedic lects (27). According to the genealogical relationship scenario, one should additionally expect a substantial number of Finno-Ugric–Yukaghir etymologies without Samoyedic cognates, which is not the case. Second, it is likely that the closest relative of Proto-Uralic is Proto-Indo-European (28), but the abnormal situation with Yukaghir repeats itself in this instance as well: there are exclusive Uralic–Yukaghir matches without any parallels in Indo-European, but there are no (or almost no) exclusive Indo-European–Yukaghir matches without any parallels in Uralic.

The obvious alternate scenario which implies trivial adstrate loans from Samoyedic to Yukaghir is somewhat impeded by the circumstance that most of the detected parallels belong to the basic rather than cultural lexicon (27), when the opposite should be expected for a situation of intense areal contact: it is known that cultural vocabulary is always borrowed first (29).

This brings to mind the possibility of a third scenario, according to which the majority of Uralic words may have penetrated into Proto-Yukaghir from a Proto-Samoyedic-like lect as a result of a specific substrate-adstrate configuration, cf. such cases as the Malol, Niuafo’ou (30) and Reo Rapa (31) languages, when a population shifts to the language of the neighboring dominating population while retaining some basic words of its original (now abandoned) language. These prehistoric contacts may have taken place in South Central Siberia. The Uralic-Yukaghir interactions require scrupulous linguistic analysis, which is a task for further studies. At the current stage of research, we conclude that our formal statistical test has yielded results which conform to the expert views regardless of whether the Uralic-Yukaghir parallels are inherited or of contact origin.

Thus, in the Chukotko-Kamchatkan–Nivkh language pair the most likely scenario is genealogical relationship, implying that the Chukotko-Kamchatkan and Nivkh languages originate from a common proto-language. On the contrary, for the Samoyedic-Yukaghir language pairs, a contact scenario (possibly with an ancient language shift) is more probable.

Statistic testing of the two theories concerning external connections of the Yeniseian family (Table 2) detects a signal of ancient relationship for both pairs: Yeniseian-Burushaski (*p* < 0.01) and Yeniseian–Na-Dene (0.01 ≤ *p* < 0.05). It is not clear whether such *p-*values can be directly compared to linguistic distances, but it is worth noting that the obtained probabilities fit S. Starostin’s phylogeny of the Sino-Caucasian macrofamily [[Yeniseian, Burushaski], Na-Dene] very well, although the result does not directly contradict Ruhlen & Vajda’s idea about Na-Dene as the closest relative of Yeniseian. It is also worth noting that both Kets and Samoyedic speakers are among the closest present-day Siberian relatives (1, 14) of Paleo-Inuit (who in turn contributed 30-40% of ancestry to a common ancestor of Na-Dene speakers, but not to speakers of other Native American language families, (1)). However, it is impossible to say if the distinctive ancestry of Yeniseian speakers survives today, since present-day Kets are genetically indistinguishable from Samoyedic-speaking Selkups (17). Thus, genetic parallels for the Yeniseian–Na-Dene and Yeniseian–Burushaski language comparisons are impossible to investigate.

An important factor in the Yeniseian problem is that historical data show a south-to-north movement of the Yeniseian-speaking population over the last several millennia. Modern-day Kets occupy a more northern area along the Middle Yenisei as compared to the area of the Yeniseian-speaking peoples in the 17^th^ c. AD (32, 33). These historical Yeniseians occupied the Middle Yenisei taiga region during the Iron Age (5^th^ c. BC to 5^th^ c. AD) (34). As was traditionally supposed by Soviet archaeologists (35–37), cf. (34), the originators of the Iron Age Yeniseian migration were the bearers of the Late Bronze Age Karasuk culture, which is dated back to the 14^th^–10^th^ centuries BC (38). This culture flourished in the Minusinsk Hollow (39) to the south of the historical Yeniseian area, partly overlapping with the latter. There are several lines of evidence for an association between the Proto-Yeniseian-speakers and the Karasuk culture: i) material ties between Karasuk and the historical Kets (35–37); ii) present-day Yeniseian-language toponymics (especially hydronyms), characteristic of the territory of the Karasuk culture (36, 40, 41); iii) recent genomic data (17) show that the Ket population is the closest modern relative of the ancient Karasuk people (42). Chronologically the Karasuk culture is likely to have spread in a south-to-north direction along the Minusinsk Hollow (43, 44), being a result of convergence of the local Andronovo culture and newcomers who penetrated into the Minusinsk Hollow from the south and/or the west, i.e., from the territories of modern day Tuva, Mongolia and/or Kazakhstan (43–46); contrary (39). Such a south-to-north movement is in agreement with the Yeniseian-Burushaski hypothesis and does not contradict a more ancient relationship with Na-Dene but makes the binary Yeniseian–Na-Dene scenario much less attractive.

## Materials and Methods

### Linguistic Methods

We compared 110-item Swadesh wordlists for 8 reconstructed protolanguages and 4 modern languages of the Circumpolar area (plus Proto-Burushaski — a language of the Hindukush) using the consonant class encoding (that is, any given root is represented as a bi-consonantal skeleton with the shape *CC*, see Suppl. Text 2 for detailed description) and the weighted permutation test in order to establish what is the probability of getting the observed sound similarities between the lists, i.e., whether these are due to chance or should be treated as evidence for prehistoric language relationship/contacts.

The following language families/groups were analyzed: Yeniseian, Athabaskan & Eyak & Tlingit, Haida, Eskimo & Aleut, Chukotian & Itelmen, Nivkh, Samoyedic, Yukaghir, and Burushaski. (See Fig. 1 for the map, and Suppl. Text 5 for description of the language groups.)

There is a consensus or near-consensus among experts that some of the aforementioned (proto)languages are genealogically related to each other. The most prominent case is the generally accepted Na-Dene family, whose structure is usually seen as follows: [[Athabaskan, Eyak], Tlingit] (47, 48). However, Proto-Na-Dene historical phonology is not sufficiently advanced, and the lexical heritage shared by Athabaskan-Eyak and Tlingit is too scant — both facts make the reconstruction of a Swadesh wordlist for Proto-Na-Dene nearly impossible. The situation with Na-Dene is comparable with, say, the Indo-European family if the only attested Indo-European remnants were the bulk of modern Germanic languages, Modern Greek and a hypothetical descendant of Hittite. Because of this we first reconstructed the Proto-Athabaskan-Eyak wordlist and compiled a synchronic wordlist for Tlingit, whereupon a cumulative Proto-Athabaskan-Eyak + Tlingit wordlist was generated by adding up the Proto-Athabaskan-Eyak and Tlingit lists. The resulting “Na-Dene” wordlist thus has two or more synonyms for each Swadesh concept (one from Proto-Athabaskan-Eyak and one from Tlingit); such synonyms were then processed by the algorithm in the same way as it processes regular language synonymy.

A similar situation is observed for the Chukotko-Kamchatkan family (49), consisting of the distantly related Chukotian and Itelmen; for this pair we prefer to compile a cumulative Proto-Chukotian + Proto-Itelmen wordlist. Finally, a third cumulative list is used for the Eskimo-Aleut pair (50). Although phonological correspondences between Eskimo and Aleut are not very complex (51), the Eskimo-Aleut relationship is distant and the number of Eskimo-Aleut etymologies within the 110-item wordlist is quite modest, so in any case we would have to use a separate Proto-Eskimo and a separate Proto-Aleut form as technical synonyms for the majority of Swadesh concepts.

Summing up, the following eight 110-item wordlists were submitted for analysis:

1. Proto-Yeniseian
2. cumulative Proto-Athabaskan-Eyak + Tlingit (*scil*. Na-Dene)
3. Haida
4. cumulative Proto-Eskimo + Proto-Aleut
5. cumulative Proto-Chukotian + Proto-Itelmen (*scil*. Chukotko-Kamchatkan)
6. Proto-Nivkh
7. Proto-Samoyedic
8. Proto-Yukaghir

The ninth wordlist that was used for an additional experiment is Proto-Burushaski. All the reconstructed lists were based on high quality wordlists for modern languages compiled according to the strict semantic specifications accepted in the Moscow School (52). Semantic reconstruction within protolanguage wordlists is based on principles generally accepted by historical linguistics and currently formulated as a set of formal criteria: tree topology, external etymology, internal derivability, typology of semantic shifts, areal effect exclusion (28). See Suppl. Table 6 for lexical lists used. Detailed linguistic notes on the reconstructed proto-forms and their meanings are offered in Suppl. Text 6.

To obtain the probability *p* of getting the observed level of phonological matches between the lists by chance, a weighted permutation test was developed. The design of the permutation test is the same as in an already published study on the Indo-European-Uralic lexical comparison (28, 53) with one important modification: weights of Swadesh concepts depend on the typological stability of individual concepts (54, 55); see Suppl. Text 2 for further details), so that a phonological match for a more stable concept is more expensive than a match for a less stable concept, since the latter is more likely to represent a chance coincidence due to its general instability. In the weighted permutation test, the first of the two wordlists compared undergoes randomized reshuffling. If the sum of costs for all concepts showing phonological matches between two original lists is *X* (total cost), the number of trials with the total cost ≥ *X* divided by the total number of trials produces the probability *p*_1_. The procedure is then repeated with the second list being reshuffled, which produces the probability *p*_2_. The probability *P* of getting the level of phonological matches observed between the original lists by chance is max(*p*_1_, *p*_2_). See Suppl. Text 1-2 for a detailed description of the consonant classes and the weighted permutation test. The list of 110 Swadesh concepts ranked by stability is offered in Suppl. Table 5.

### Haplotype Sharing Statistics

Shared haplotypes were inferred on a phased HumanOrigins genotype panel (56) using ChromoPainter v.1 (57) for pairs of individuals (the same dataset was used by (1), but the questions explored here were not touched in that publication). For a tested individual the length of shared haplotypes in cM was averaged across all individuals in the reference population (Samoyedic speakers). That statistic, “Samoyedic HSS”, was normalized by another similar HSS, between the tested individual and other non-Uralic-speaking Siberians (see a list of individuals and group labels in Suppl. Table 1). The normalized HSS, “Samoyedic/other Siberian HSS”, was calculated for Samoyedic-speaking, Yukaghir, Ugric-speaking, and other Siberian individuals using a leave-one-out procedure. Optionally, normalized HSS were averaged across test populations. To test the significance of differences in “Samoyedic/other Siberian HSS” between Yukaghir and other Siberian groups, we performed a permutation test for all populations by randomly sampling the individual HSS values with replacement using the “coin” package in R (58).

### Principal Component Analysis (PCA) and Multi-Dimensional Scaling (MDS)

For exploring signals of gene flow between Chukotko-Kamchatkan and Nivkh speakers, we compiled a dataset based on the 1240K SNP panel (18) using public whole-genome shotgun data for present-day populations (19, 59–61) and ancient genomes (20–22, 62–71).

We computed PCA using Plink v1.9 with default options, taking only relevant populations into the analysis. To calculate MDS, we first computed pairwise outgroup *f*_*3*_-statistics of the form *f*_*3*_(Yoruba; Individual_1_, Individual_2_) where individuals belonged to Siberian, Chukotko-Kamchatkan, and Southeast Asian populations used for constructing admixture graphs. We generated a dissimilarity matrix based on 1-*f*_*3*_-statistics and then calculated MDS using a base R function `cmdscale()`.

### Admixture graphs

A set of genotypes used in this study (1,062,979 SNPs derived from the 1240K panel and including both transversions and transitions) is identical to that used by Ning et al. (14). For a list of individuals and group labels see Suppl. Table 3. We used four alternative Chukotko-Kamchatkan-speaking groups: Chukchi from two sources (59, 60), Itelmen, and Koryaks. We also used eight Tungusic-speaking groups (Evens, Evens from Magadan, Evens from Yakutia, Evenks, Hezhen, Oroqen, Ulchi, and Xibo), Nivkhs, and one Turkic-speaking group (Tuvinians) as a negative control (Suppl. Table 3). The former groups are termed C-K for brevity, and the latter groups are termed TN (Tungusic and Nivkh). C-K groups were grafted onto the skeleton graph as derived from the Saqqaq and First American branches, and also receiving gene flows from all distal West Eurasian sources (West European hunter-gatherers, Mal’ta, and basal Eurasians). As shown by Ning et al. (14), these modelling rules allow to find realistic admixture models for groups known to harbour a complex mixture of both Asian and European ancestries (e.g., present-day Aleuts, (19, 61)). TN groups and Tuvinians were grafted on the skeleton graph as derived from three Asian sources: the ARB branch, the Nganasan and Jomon branches, and also receiving gene flows from all three distal West Eurasian sources.

For each combination of the groups, the following models were tested: a baseline model with no gene flows between the C-K and TN lineages (Suppl. Fig. 2); a model with a C-K to TN gene flow; a model with a TN to C-K flow; and a model with gene flows in both directions. Since these models have different number of parameters (edges and gene flows), comparing their likelihoods is not straightforward (14). We used a variation of the bootstrapping model comparison method proposed by Ning. et al. For each bootstrap iteration *i*, we drew bootstrap samples of 5 cM SNP blocks (this block size is a standard in the AdmixTools package) and fitted two models being compared on the resampled dataset. The difference in log-likelihoods (LL) of two graphs was calculated for each resampling iteration, and since the distribution of LL differences tends to deviate from normality, an empirical one-tailed *p*-value was computed. We used one-tailed tests since we did not expect that a simpler model would ever demonstrate a significantly better fit than a more complex model. A one-tailed empirical *p*-value is a proportion of negative LL differences among all bootstrap replicates, where the first LL value corresponds to a less complex model and the second LL value to a more complex one. We note that the covariance matrix of *f*_*3*_-statistics, which is necessary for computing the model likelihood score, was calculated on the original non-resampled dataset. The model-comparison method is a part of a new implementation of AdmixTools (AdmixTools 2, https://github.com/uqrmaie1/admixtools).

## Acknowledgements

The article was prepared in the framework of a research grant funded by the Ministry of Science and Higher Education of the Russian Federation (grant ID: 075-15-2020-908). This work was supported by the Czech Ministry of Education, Youth and Sports from the Large Infrastructures for Research, Experimental Development and Innovations project “IT4Innovations National Supercomputing Center – LM2015070”. O.F., P.C., and P.F. were supported by the Institutional Development Program of the University of Ostrava. The study was funded by a subsidy from the Russian federal budget (project No. 075-15-2019-1879 “From paleogenetics to cultural anthropology: a comprehensive interdisciplinary study of the traditions of the peoples of transboundary regions: migration, intercultural interaction and picture of the world”). The work of SAS was carried out within the Research Topic 0065-2019-0007 of the Russian Federation Scientific state assignment.

