## Supplementary Text for "Circumpolar peoples and their languages: lexical and genomic data suggest ancient Chukotko-Kamchatkan–Nivkh and Yukaghir-Samoyedic connections"

### Supporting Information: text (linguistics)

### 1. Consonant classes

We use the same technique of consonant coding as described in Kassian, Starostin & Zhivlov 2015a.

All linguistic data in the present article are encoded in the unified transcription system of the *Global Lexicostatistical Database* project (http://starling.rinet.ru/new100/), which is generally based on the IPA alphabet, with several specific discrepancies, e.g., we use traditional *c č* for IPA *t͡s t͡ʃ*. See http://starling.rinet.ru/new100/UTS.htm

Further each proto-root is encoded according to its constituent consonant classes, that is, any given root is represented as a bi-consonantal skeleton with the shape *CC*. Since each consonant class was designed to include phonemes which mutate more frequently into each other during language evolution than into phonemes of other classes, two forms from the compared wordlists with identical *CC*-transcriptions have a higher chance to be historical cognates than forms whose *CC*-transcriptions differ. Our algorithm marks any pair of forms with the same *CC*-transcription as a *CC*-match, and other pairs as non-matching. The method of consonant classes is thus, on one hand, a crude variation on the measurement of Levenshtein distances and, on the other hand, is close to modeling the preliminary stage of real comparative-historical research, at least as far as criteria for eliciting potential etymological lexical matches between two languages are concerned.

We use the following consonant classes: Table S1.

Table S1. Consonant classes

| | |
|---|---|
| P-class (labials) | p b ɓ β f v… |
| T-class (dentals) | t d ɗ θ ð ɖ ʈ … |
| S-class (sibilant fricatives) | s z š ž… |
| Ʒ-class (sibilant affricates) | c ʒ č ǯ… |
| Y-class (palatal glides) | y… |
| W-class (labial glides) | w ʍ… |
| M-class (labial nasals) | m ɱ… |
| N-class (non-labial nasals) | n ɳ ɲ ŋ… |
| Q-class (lateral affricates) | ƛ … |
| R-class | r ɾ… |
| L-class | l ɭ ɬ ɫ… |
| K-class (velars & uvulars) | k g x ɣ q χ ʁ… |
| zero-class or H-class | ħ ʕ ʜ ʢ ʡ h ɦ ʔ and any vowels |

This proposed transcription system (*P T S Ʒ Y W M N Q R L K H*) is sufficient for encoding all wordforms or morphemes of any natural language that is included into comparison.

Elements of the *H*/zero-class and such features as coarticulation, prosody, phonation are notably deleted from the structure, with the exception of word-initial and word-final vowels and laryngeals which are coded as *H*.

Non-initial *Y* and *W* (weak glides) are treated as *H*, i.e., they are deleted in the medial position and coded as *H* in the final position.

As noted above, for the present study we use *CC*-transcription, reducing each wordform to its two first consonants.

Examples of how the consonant class encoding actually works are given in Table S2.

Table S2. Examples of transcription of consonant classes (hypothetical wordforms).

| **Wordform** | **Full consonant classes transcription** | ***CC*-transcription** |
|---|---|---|
| *tasam* | *TSM* | *TS* |
| *dʰüzo* | *TSH* | *TS* |
| *alaq* | *HLK* | *HL* |
| *ʔäʎx* | *HLK* | *HL* |
| *na* | *NH* | *NH* |
| *ŋoʔ* | *NH* | *NH* |
| *pkʰot* | *PKT* | *PK* |
| *baq’aƛ* | *PKQ* | *PK* |
| *wahat* | *WT* | *WT* |
| *ʍad* | *WT* | *WT* |
| *ka* | *KH* | *KH* |
| *kay* | *KH* | *KH* |
| *kawa* | *KH* | *KH* |
| *kat* | *KT* | *KT* |
| *kayat* | *KT* | *KT* |

#### 2. Weighted permutation test

When two isomorphic wordlists (with roots transcribed into *CC*-shapes) are compared with each other, one of the lists is randomly reshuffled, and the number of *CC*-matches is recorded for each new configuration. In the traditional (i.e., unweighted) permutation test, if the number of observed *CC*-matches between two wordlists is $X$, the number of random trials with $X$ or more *CC*-matches divided by the total number of trials produces the probability $p$ of getting at least the same number of $X$ matches as between the original wordlists (Oswalt 1970; Baxter & Manaster Ramer 2000; Kassian, Starostin & Zhivlov 2015a; 2015b).

In order to enhance the signal, we developed a weighted permutation procedure, where each Swadesh concept is assigned its own weight (or cost) in accordance with its typological stability.

It is commonly acknowledged that Swadesh concepts possess different average degrees of stability: some concepts are typologically more stable, i.e., words that designate these concepts usually last longer in the language, while other concepts are less stable, and the corresponding words disappear or change their meanings more frequently in the course of language evolution. Based on S. Starostin's (2007a) typological survey of language families of the Old World, we calculated the degree of stability for each concepts and used them as multiplication factors to increase the cost of *CC*-matches, so that a *CC*-match for a more stable concept is more expensive than a *CC*-match for a less stable concept (since the latter has a higher chance to represent a chance coincidence due to its general instability).

S. Starostin (2007a) offers statistical data on lexical stability of 110 Swadesh concepts in some families of the Old World: 132 Sino-Tibetan lects, 99 Austro-Asiatic lects, 54 Altaic lects, 94 Austronesian lects, 36 Australian lects, 26 Khoisan lects, 33 North Caucasian lects, 21 Dravidian lects, 97 Indo-European lects, 7 Kartvelian lects, 69 Afroasiatic lects, 47 Tai-Kadai lects, 17 Uralic lects, 14 Yeniseian lects. In total, S. Starostin's sample consists of 737 languages belonging to 14 language families.

S. Starostin (2007a) himself proposed a relatively complex and not intuitively transparent calculation of the stability index. Instead of this approach, we prefer to follow a simpler and more straightforward way proposed by Pozdniakov (2014). First, the "stability index" of each Swadesh item for each family is defined as $M/L$, where $L$ = the number of languages in the family and $M$ = the maximum number of languages within the group that use reflexes of the same root for the respective Swadesh meaning (e.g., the Slavic stability index for 'belly' is 0.38, since 5 out of 13 languages preserve reflexes of the same Proto-Slavic root **bryo:x*-); this part of the procedure is the same as in S. Starostin's original article. At the next step, for each Swadesh concept we take the arithmetic mean of its stability indexes in individual language families. The obtained number is the stability index of the given concept:

| | |
|---|---|
| 'I' | 0.805 |
| 'thou' | 0.797 |
| 'two' | 0.769 |
| 'eye' | 0.738 |
| 'we' | 0.725 |
| ... etc. | |

We present a spreadsheet showing stability indexes of Swadesh concepts as a separate MS Excel file. Our ranking is almost identical with Table 10 from Pozdniakov 2014 (minor rearrangements are due to rounding).

These indexes were further used as weights of individual Swadesh concepts. When two wordlists are compared, the sum of weights of all concepts with positive pairs (*CC*-matches in our cases) constitutes the total weight of that comparison. When the same slot is occupied by several synonyms (a normal situation in our study), we compare all possible pairs between two languages: if there is at least one matching pair, that pair is treated as positive.

For instance, the following positive pairs (CC-matches) were observed for the Haida and Nivkh wordlists (see SI for lexical data): Table S3.

Table S3. Positive pairs (*CC*-matches) between the Haida and Nivkh 110-item wordlists.

| concept | weight | Haida | Nivkh | CC-transcription |
|---|---|---|---|---|
| dog | 0.628 | *χa* | *ɢa* | *KH* |
| dry | 0.442 | *k'a:* | *qaw* | *KH* |
| sleep | 0.430 | *q'a* | *qo* | *KH* |
| that | 0.339 | *hu:* | *hu* | *HH* |

The total weight of matching pairs in the original non-reshuffled lists is 184 (multiplied by 100 and rounded for the sake of convenience).

The permutation test begins by randomly reshuffling one of the two lists (the Nivkh one in our case), checking the total weight for each new configuration. We routinely used 1.000.000 pseudo-random trials, but if all trials returned a smaller total weight than observed for the original lists, we reran the analysis with 10.000.000 trials.

If $s$ is the weight of the original wordlist comparison ($s$ = 184 in the Haida-Nivkh example), the probability $p_1$ of getting at least a weight $s$ as between the original wordlists is the number of trials with weight ≥ $s$ divided by the total number of trials.

The outcome of the Haida-Nivkh weighted permutation test looks as follows ($s$ is the weight):

s = 0: 51094 trial(s)
s = 22: 1018 trial(s)
s = 23: 1036 trial(s)
...
**s = 184: 3171 trial(s)**
s = 185: 3164 trial(s)
s = 186: 3090 trial(s)
...
s = 534: 1 trial(s)
s = 540: 1 trial(s)
s = 572: 1 trial(s)

It can also be depicted as a plot (Fig. S1).

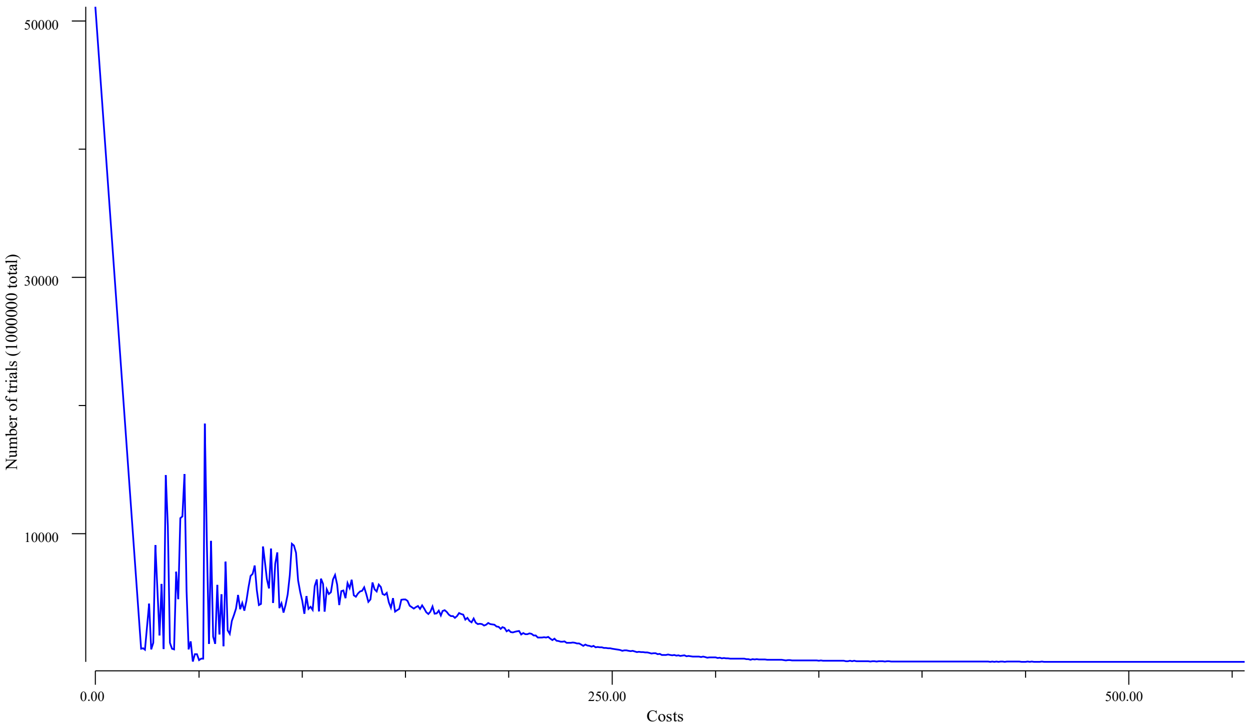

Fig. S1. Haida-Nivkh weighted permutation test (the Nivkh list is being reshuffled).

The number of trials that yielded $s \geq 184$ is 3,171 + 3,164 + ... + 1 = 187,327, thus $p_1$ = 0.187.

Then we repeat the permutation procedure, keeping the second list untouched whilst the first one (Haida in our case) is being reshuffled. This produces another probability estimate, $p_2$ (note that $p_1$ and $p_2$ are normally close to each other, but not expected to be equal). In the Haida-Nivkh case, $p_2 = 0.233$.

In order to reduce the chance of making a type 1 error, we accept the highest of two obtained $p$-values as the final estimate: the probability $P$ of getting the observed level of phonological matches between two lists by chance = $\max(p_1, p_2)$. In the Haida-Nivkh example, $P = 0.233$.

The software package used in the present study is available at https://github.com/oopcode/starling-permutation-test.

#### 3. Pairwise comparison between Yeniseian, Burushaski and all the Na-Dene wordlists

Tab. S4. Probabilities of phonetic matches between Yeniseian, Burushaski and the (proto-)languages that constitute the Na-Dene family obtained by the weighted permutation test.

| | **Tlingit** | **Eyak** | **Proto-Athabaskan** | **Proto-Athabaskan-Eyak** | **Proto-Athabaskan-Eyak + Tlingit** | **Proto-Burushaski** |
|---|---|---|---|---|---|---|
| **Proto-Yeniseian** | 0.557 | 0.023 | 0.013 | 0.006 | 0.014 | 0.009 |
| **Proto-Burushaski** | 0.364 | 0.847 | 0.066 | 0.289 | 0.247 | — |

Note the insignificant result of the Yeniseian-Tlingit pair ($p$ = 0.557), although comparison between Yeniseian and other non-Yeniseian lists yields $p < 0.05$.

#### 4. *CC*-matches in statistically significant pairs (Chukotian-Itelmen, Nivkh, Samoyed, Yukaghir, Yeniseian, Na-Dene, Burushaski)

Table S5. Positive pairs (*CC*-matches) between the “Proto-Chukotko-Kamchatkan” (cumulative Proto-Chukotian + Proto-Itelmen) and Proto-Nivkh 110-item wordlists.

| concept | “Proto-Chukotko-Kamchatkan” | Proto-Nivkh | CC-transcription |
|---|---|---|---|
| big | pəl | bil | *PL* |
| eat | nu | ɲi | *NH* |
| meat | tərɣe | dur | *TR* |
| new | tur- | ṭur- | *TR* |
| not | kä | qaw | *KH* |
| sit | təva | tiv | *TP* |
| smoke | t’i | taw | *TH* |
| star | äŋär | uɲ | *HN* |
| this | tiʔ | du | *TH* |
| tongue | il | hilɣ | *HL* |
| we | mur | mir | *MR* |

Table S6. Positive pairs (*CC*-matches) between the Proto-Samoyed and Proto-Yukaghir 110-item wordlists.

| concept | Proto-Samoyed | Proto-Yukaghir | CC-transcription |
|---|---|---|---|
| feather | tuǝ̆ | tiw | *TH* |
| I | mǝ̆ | mə | *MH* |
| mouth | aŋ | aŋa | *HN* |
| say | mɒn | mon | *MN* |
| that | ta | ta | *TH* |
| this | tǝ̆ | ti | *TH* |
| thou | tǝ̆ | tə | *TH* |
| we | me | mi | *MH* |
| worm | kǝ̆lǝ̆ | kelinčʸə | *KL* |

Table S7. Positive pairs (*CC*-matches) between the Proto-Yeniseian and “Proto-Na-Dene” (cumulative Proto-Athabaskan-Eyak + Tlingit) 110-item wordlists.

| concept | Proto-Yeniseian | “Proto-Na-Dene” | CC-transcription |
|---|---|---|---|
| ashes | qol | kʰéɬ’ | *KL* |
| big | qeʔ | keː | *KH* |
| fingernail | xiːɲe | χanc | *KN* |
| dry | qɔɢ | xuːk | *KK* |
| give | o | ʔa | *HH* |
| kill | xeːy | χeː | *KH* |
| liver | seŋ | sVnt’ | *SN* |
| louse | xəːke | kVks | *KK* |
| mouth | qowe | χ’é | *KH* |
| see | ɔŋ | ʔVːn | *HN* |
| stone | ciʔ | cʰVː | *CH* |
| that | ʔu | ʔVw | *HH* |
| go | hey | haː | *HH* |
| short | tuk | tik’ | *TK* |

Table S8. Positive pairs (*CC*-matches) between the Proto-Yeniseian and Proto-Burushaski 110-item wordlists.

| concept | Proto-Yeniseian | Proto-Burushaski | CC-transcription |
|---|---|---|---|
| dry | qɔɢ | qaq | *KK* |
| eat | siː | ʂi | *SH* |
| give | o | u | *HH* |
| kill | xeːy | qa | *KH* |
| name | ʔiɢ | ek | *HK* |
| that | ʔu | i | *HH* |

### 5. Basic information on the language families and groups

#### Yeniseian (George Starostin)

The Yeniseian family (Vajda 2001; Anderson 2004) consists of several languages, out of which only Ket is currently surviving, with approximately 200 remaining speakers in several villages located in the Middle Yenisei basin; however, several other members of the Yeniseian group have been attested as early as the 18th century, and evidence from history and toponymics suggests that the original extent of the family was much larger, reaching South Siberia and stretching almost to Lake Baikal. Besides Ket, abundant linguistic data have been preserved on Yugh (Sym), a language that was very closely related to Ket before its extinction at the end of the 20th century; somewhat less well described (primarily due to the research of M. Castrén) is Kott, extinct in the middle of the 19th century; and even scarcer and less reliable (though still historically priceless) data have been preserved on Arin and Pumpokol, two languages spoken further to the south and presumably extinct by the beginning of the 19th century.

Despite the fact that the majority of Yeniseian languages became extinct before their data could be captured by means of modern day linguistic fieldwork practices, data from the late 18th century survey sources on Arin and Pumpokol and particularly the data on Kott, collected by Castrén, permit proper comparative-historical research to be applied to the Yeniseian languages, and a detailed formal reconstruction of Proto-Yeniseian phonology and lexicon was laid out by Sergei Starostin (2007b [1982]), followed by a brief etymological dictionary of the reconstructed Proto-Yeniseian (Starostin 1995). Since then, a slightly amended alternative model for Proto-Yeniseian has been offered by Heinrich Werner (2002), who is currently working on an even more in-depth reconstruction in collaboration with Edward Vajda. Many of Werner's reconstructions are significantly different from S. Starostin's and have occasionally been criticized by the latter in his 2003–2004 notes in the database (Starostin 2005a).

The Yeniseian family as a whole is not too divergent, with much of the basic lexicon, root structure peculiarities, and overall phonological features of the proto-language well preserved in all of its daughter languages; however, the grammatical structure of Proto-Yeniseian is significantly harder to reconstruct due to its notorious complexity and lack of reliable data on such extinct languages as Arin and Pumpokol. Our glottochronological calculations, performed on carefully assembled 100-item Swadesh wordlists of the basic lexicon for the five Yeniseian languages, yield an approximate date of 700–500 BC for the disintegration of Proto-Yeniseian, and a tripartite structure of the family, with indisputable Ket-Yugh and Kott-Arin branches, and a questionable status for Pumpokol (probably somewhat closer to Ket-Yugh than to Kott-Arin, although this is questionable in light of numerous Yugh words formerly mislabeled as Pumpokol in older sources). Cf. the identical Yeniseian tree in Georg 2008.

Our protolanguage wordlist takes S. Starostin's phonological reconstruction as its starting point; however, Werner's alterations to the reconstructions are considered on a regular basis, and some modifications to the etymologies have also been suggested by G. Starostin (all such modifications are stated and justified in the notes section).

The present attempt at the reconstruction of a Swadesh wordlist for Proto-Yeniseian generally coincides with Starostin 2013a (several emendations are explicitly discussed below, most of them consist of an extra synonym added to the comparison in situations when two roots are found to have almost even chances of having expressed the required Swadesh meaning on the proto-level: 'breast', 'cold', 'to come', 'hand', 'to know', 'moon', 'round', 'short').

Seven Swadesh items are not reconstructible for Proto-Yeniseian (due to insufficient attestation or presence of new formations in the majority of lects): 'belly', 'to bite', 'green', 'seed', 'skin', 'yellow', 'snake'.

#### Na-Dene (Proto-Athabaskan, Eyak, Proto-Athabaskan-Eyak, Tlingit) (Alexei S. Kassian)

The general expert consensus on the Na-Dene family is that it consists of one large group, the Athabaskan languages (chronologically the Athabaskan group is probably slightly deeper than, for instance, the Germanic languages) and two outlier languages: Eyak and Tlingit. Out of these two, Eyak is definitely closer to Athabaskan than Tlingit (e.g., Krauss 1976; Kari 2010: 208), and the Athabaskan-Eyak relationship is visible to the naked eye. Tlingit, on the other hand, is much more distant, and since there are numerous points of uncertainty in phonological and morphological comparison of Tlingit with other Na-Dene languages, it is impossible to propose a reliable Proto-Athabaskan-Eyak-Tlingit phonological and semantic reconstruction at the current stage of research.

Thus, the Na-Dene section of our study consists of 4 Swadesh wordlists: a reconstructed list for Proto-Athabaskan, a synchronic list for Eyak, a reconstructed list for Proto-Athabaskan-Eyak, and a synchronic list for Tlingit.

Lexical data are taken from the GLD database: Athabaskan (Kassian 2011a, ongoing project), Eyak (Kassian 2011b), Tlingit (Kassian 2011c).

We proceed from the conservative model with a three-way division of Athabaskan: Pacific Coast, Apachean (Southern), and Northern (e.g., Kari 2010: 208). It is clear that the Northern group represents a complex tree with several subgroups and may even be polyphyletic. Nevertheless, such a rough three-way classification is enough for semantic reconstruction of the overwhelming majority of the Swadesh concepts.

Our phonological reconstruction of Proto-Athabaskan and Proto-Athabaskan-Eyak forms generally follows previous studies: Krauss & Leer 1981; Leer 1996; Leer 2008b; Leer 2010, and so on, as well as some proposals in Nikolaev 2014. We intentionally do not take into account some marginal sound correspondences between Proto-Athabaskan and Eyak, since these require additional investigation.

We do not go into details concerning exact phonological shapes of the reconstructed forms unless it could affect transcription of consonant classes. In particular, we often write simple **V* when it is difficult to reconstruct the appropriate vowel with any precision. However, we use the phonological transcription *t* *t*$^h$ *t’* for the traditional orthographic triad ⟨d t t’⟩. The Proto-Athabaskan palatalized velar row **k*$^y$ *k*$^{yh}$ *k*$^y$*’* *x*$^y$ *ŋ*$^y$ is to be reinterpreted as the plain velars **k* *k*$^h$ *k’* *x* *ŋ* (which are opposed to the uvular row **q* *q*$^h$ *q’* *χ*).

A crucial thing for our computational procedure is the reconstruction of Proto-Athabaskan-Eyak clusters with nasals, **nC*. Leer (2008a) proposed that the PAE sequence **VN* yielded the aspirated vowel phonation in Eyak, i.e., PAE **VN#* > Eyak *Ṽ*$^h$~*V*$^h$, **VNC* > Eyak *Ṽ*$^h$*C*~*V*$^h$*C*, and that the old nasalization is correspondingly the main source of the aspirated vowel in Eyak. E.g., Eyak *t=k*$^h$*ĩ*$^h$ ‘stick, wood’ / PA **tə=k*$^h$*ən* ‘stick, tree, wood’. Note that synchronous Eyak variations *Ṽ*~*V* (e.g., *=quht* ~ *=qũht* ‘knee’), the absence of nasalized *e* and many cases when PA nasals corresponds to Eyak plain vowels (Krauss & Leer 1981: 140) suggest that Eyak was documented at the moment when vowels began to lose the nasal coarticulation. In order to explain non-nasalized reflexes in Eyak, Leer (2008) additionally hypothesizes about rare non-homorganic clusters, PAE **VɲC* or **VmC*, but such a solution seems superfluous; for the present paper we reconstruct the only PAE nasal **n* for all cases of the Eyak aspirated vowels.

The synchronous Eyak vowel system, Table S9, supports Leer’s general idea. Eyak short vowels lack plain nasalized phonemes (***ã*), whereas long vowels lack aspirated phonemes (***aː*$^h$, ***ãː*$^h$). This may imply that Proto-Eyak short nasalized vowels yielded aspirated vowels **ã* > *ã*$^h$~*a*$^h$ (either nasalized or not in documented Eyak), whereas Proto-Eyak long nasalized vowels avoided aspiration (**ãː* is retained as is or denasalized > *aː*). Thus the first restriction for the Eyak vowel aspiration is the Proto-Eyak vowel length, i.e., aspiration does not affect Eyak vowels which retain PAE length. The second restriction is the Eyak vowel glottalization which prevented aspiration (*ã'* is retained as is or denasalized > *a'*).

Table S9. Types of Eyak vowels (after Krauss 1965: 169).

| | plain | aspirated | glottalized | nasalized | nasalized aspirated | nasalized glottalized |
|---|---|---|---|---|---|---|
| **short** | ə (< *a) | $a^h$ | a' | — | $ã^h$ | ã' |
| **long** | aː | — | aː' | ãː | — | ãː' |

Since sequences of the shape **CVNC* are not reconstructible for Proto-Athabaskan (except for morpheme boundaries, e.g., as in 'nose'), an important implication of the Leer's idea is that, if the Proto-Athabaskan sequence **CVC* corresponds to Eyak *CṼʰC* or simply *CVʰC* (with recent denasalization), the PAE form is to be reconstructed with a nasal cluster, **CVnC*. E.g., Eyak *łãʰt* 'smoke' and PA **łət* 'smoke' should go back to PAE **łant*, or Eyak *=saʰt* 'liver' and PA **=zət'* 'liver' should go back to PAE **sant'*. Additionally, as noted by Leer (2008a) the Eyak verbal paradigm does not allow to discriminate between verbal roots of the shapes **=CVːN* and **=CV(ː)* due to levelled TMA suffixes such as imperfective/perfective *-h* (< **ɲ*, Krauss & Leer 1981: 38-39), imperative *-ʔ* etc.

The second and more technical thing about the Proto-Athabaskan reconstruction of nasal consonants. There are at least two nasals reliably reconstructed for Proto-Athabaskan. The first one was probably **n*, but the phonetic nature of the second is debatable. The second nasal is traditionally interpreted as velar **ŋ* or **ŋʸ* (e.g., Krauss & Leer 1981), but later it was proposed to reinterpret it as palatal **ɲ* (e.g., Leer 2010). The choice between *ŋ* and *ɲ* is irrelevant for our computational procedure (*ŋ* and *ɲ* fall in the same consonant class), but the palatal interpretation **ɲ* seems slightly more apt in the light of internal and external comparison (Leer 2010: 172). In such a case, it is natural to interpret the third PA nasal, Krauss & Leer's (Krauss & Leer 1981: 14-15; Leer 2010: 172) **ŋ₂*, as **ŋ* since the shift *ŋ* > *m* (as in Pacific Coast Athabaskan) is normal cross-linguistically.

We follow Leer 2010 and interpret the Proto-Athabaskan labialized hushing phonemes (**čʷ, *šʷ* etc.) as retroflex (**č̢ *š̢* etc.).

#### Haida (Alexei S. Kassian)

Haida is a language isolate with two dialects: Southern and Northern. Our wordlist was adapted from Kassian 2011d; it is based on Southern forms since the Southern dialect is more archaic phonetically than the Northern one (Enrico 2005: viii). There is no need to define a reconstructed protolanguage level for a family with such small time depth. Instead, in those several cases where there are lexical discrepancies between the two dialects, we either use the more archaic term (if there are internal indications on the direction of semantic development) or use both words as synonyms. In order to accommodate the Haida data to the automated consonant classes analysis we write simple *a* for the specific Southern Haida lateralized vowel which is an allophone of /a/.

#### Eskimo-Aleut (Alexei S. Kassian)

The Eskimo sub-family consists of two main groups, Yupik and Inuit, within each of which there are several closely related languages. The recently extinct Sirenik language likely represents an outlier within the Yupik group.

Our Proto-Eskimo reconstruction is generally based on Fortescue et al.'s (2010) etymological dictionary and on a number of dictionaries and glossaries of individual languages such as Central Siberian Yupik (Menovshchikov 1988), Pacific Gulf Yupik (Leer 1979), Sirenik (Menovshchikov 1964), North Alaskan Inuit (MacLean 2014), Greenlandic Inuit (Fortescue 1984), and others.

The Aleut language consists of three primary dialects: Eastern, Atkan, and Attuan, which are close to each other. Our Proto-Aleut reconstruction is based on Bergsland's (1994; 1997) cumulative dictionary and grammar, as well as sources on individual dialects: Bergsland & Dirks 1978, Golovko 1994, Golovko, Vakhtin & Asinovskiy 2009.

Eskimo and Aleut are apparently genealogically related (Bergsland 1986; 1989; 1994; Fortescue 1998; Fortescue, Jacobson & Kaplan 2010; as plausibly advocated in Berge 2016; 2018 the genealogical relationship was later obscured by several waves of Eskimo-Aleut contacts which were accompanied by lexical loans in both directions), but our lexicostatistical analysis suggests that the Eskimo-Aleut relationship is so distant that it is essentially useless to reconstruct a joint Proto-Eskimo-Aleut wordlist for our current purposes.

#### Chukotian and Itelmen (Chukotko-Kamchatkan) (Alexei S. Kassian)

The Chukotian sub-family consists of four languages, which are relatively closely related to each other: Chukchi (with dialectal diversity), Kerek (poorly documented and practically extinct), Koryak, and Alutor.

Our Proto-Chukotian reconstruction is based on Fortescue's (2005) etymological dictionary as well as on main lexicographic sources on synchronic Chukchi (Inenlikei 2005; Moll & Inenlikei 1957; Skorik 1961; 1977), Koryak (Zhukova 1967; 1972; 1990) and Alutor (Kibrik, Kodzasov & Muravyeva 2004; Nagayama 2003).

The Itelmen a.k.a. Kamchatkan sub-family consists of three closely related languages: Western (the only one spoken today; it consists of two dialects: Sedanka and Xajrjuzovo a.k.a. Napana), Eastern, and Southern (both of these are extinct and have not been documented systematically).

Reconstruction of Proto-Itelmen is a non-trivial task, because 18$^{th}$-19$^{th}$ century data on the extinct Eastern and Southern Itelmen languages are not very reliable and consistent. This problem concerns both phonology and, crucially for our purposes, semantic definitions. In particular, it means that the current Proto-Itelmen reconstruction is inevitably Western-biased, mostly relying upon modern sources on Western Itelmen. We generally follow Mudrak's (2005; a preliminary version was published as Mudrak 2000; a more complex version is Mudrak 2008) reconstruction of Proto-Itelmen with minor emendations and/or simplifications, if needed. The synchronic Itelmen data are taken from Volodin & Khaloimova 2001; Volodin 1976; Ono 2003; Stebnitsky 1934 as well as from Fortescue 2005.

Genetic relationship between Chukotian and Itelmen is generally accepted by experts (Skorik 1958; Fortescue 1998; 2003; 2005; Kurebito et al. 2001; Mudrak 2000; with hesitation Volodin 1976). The opposite opinion was expressed by Worth (1962) and Volodin (1997; Georg & Volodin 1999: 224–228), who supposed that the observed Chukotian-Itelmen matches, which cover not only basic vocabulary but also some main grammatical exponents (Fortescue 2003), are contact loans from one language group to another. Worth-Volodin's scenario, however, clearly contradicts the theory of language contacts which predicts that cultural vocabulary is always borrowed first, whereas basic vocabulary and main grammatical exponents are most protected from borrowing (Thomason & Kaufman 1988).

It could be reasonable to use a Proto-Chukotko-Kamchatkan (Proto-Chukotian-Itelmen) wordlist instead of two distinct lists for Proto-Chukotian and Proto-Itelmen respectively, but there are too many unsolved obstacles in the Chukotian-Itelmen comparison at the current stage of research, making reliable phonological reconstruction of a Proto-Chukotko-Kamchatkan wordlist impossible.

#### Nivkh (Alexei S. Kassian)

The Nivkh family consists of four closely related languages, which are sometimes called dialects: Amur, East Sakhalin, South Sakhalin, and North Sakhalin (Gruzdeva 1998: 7; Fortescue 2016: 1). North Sakhalin is traditionally described as an intermediate lect between Amur and East Sakhalin that should imply contact influence on North Sakhalin on the part of either Amur or East Sakhalin (it is likely that Amur and North Sakhalin form a distinct clade, but North Sakhalin was later influenced by the East Sakhalin language).

Our Proto-Nivkh reconstruction is generally based on Fortescue's (2016) comparative dictionary and synchronic sources: Panfilov 1962; 1965; Peiros & Starostin 1986; Savelyeva & Taksami 1965; 1970; Taksami 1996. We reconstruct the Proto-Nivkh palatal obstruents as non-sibilant plosives **ȶ *ȡ* (for Fortescue's **c *d'*). In modern lects, they tend to shift into the sibilant affricate articulation such as *č*, but typologically the development **ȶ* > *č* seems more natural than *vice versa*. We assume that clusters of the shape **rK* can shift to *sK* in Amur and North Sakhalin, being retained with *r~r̥* in East and/or South Sakhalin, whereas clusters of the shape **zK* yield *sK* everywhere.

#### Samoyed (Mikhail Zhivlov)

Samoyed is the most divergent branch of the Uralic language family. There is no consensus on the internal classification of Samoyed languages. We provisionally accept E. Helimski's classification, according to which Northern Samoyed languages — Tundra and Forest Nenets, extinct Old Eastern Yurak (sometimes erroneously called Yurats), Tundra and Forest Enets and Nganasan — form a subgroup, while the so-called Southern Samoyed languages (Mator, Kamass, and Selkup) represent three independent branches of the Samoyed group (Helimski 1982). Nevertheless, any lexical isogloss unique to "Southern" languages can in principle result from a later areal development, so that attestation in at least one Northern and one "Southern" language is crucial for assuring that a word was present in Proto-Samoyed. Of course, a word with good etymological parallels in other branches of Uralic must be reconstructed for Proto-Samoyed even if it is attested only in one Samoyed language.

Our Proto-Samoyed reconstruction follows Janhunen 1977 with important additions and corrections by Helimski (1997: 68–70; 2005). However, we do not distinguish in our transcription between front and back reduced vowels, since this distinction is morphophonemic rather than phonological.

A set of synchronic Samoyed Swadesh wordlists is offered in Koryakov 2018.

#### Yukaghir (Mikhail Zhivlov)

Tundra and Kolyma Yukaghir are frequently called "dialects" of the Yukaghir "language". In fact, they are mutually incomprehensible; moreover, the percentage of common words in the Swadesh 100-word list between Tundra and Kolyma Yukaghir is approximately the same as between Russian and Latvian. On the other hand, extinct Chuvan and Omok, traditionally viewed as outliers in the Yukaghir family, are specifically (but not closely) related to Kolyma and Tundra languages respectively. Except for Omok, all old Yukaghir wordlists, recorded in 18th and 19th centuries, represent varieties specifically related to Kolyma Yukaghir (at least from the point of view of historical phonology). Nevertheless, a lexical parallel between one of these wordlists and modern Tundra Yukaghir is not necessarily an archaism going back to Proto-Yukaghir: it can also be a common areal development.

Semantic reconstruction of the Proto-Yukaghir basic lexicon encounters serious obstacles. We can reconstruct the Proto-Yukaghir word for a given Swadesh meaning: 1) if both

Kolyma and Tundra have the same word for this meaning; 2) if the word for this meaning in one of the languages is clearly secondary, e.g., it results from unidirectional semantic development like 'to put down' > 'to kill'. In most other cases, we are forced to list both Kolyma and Tundra words in Proto-Yukaghir phonological garb as technical synonyms.

Our phonological reconstruction of Proto-Yukaghir generally follows Nikolaeva 2006.

#### Burushaski (George Starostin)

Burushaski (Berger 1998; Driem 2001) is essentially a single macro-language, spoken by approximately 87,000 people in northern Pakistan (the Gilgit-Baltistan region), with two very closely related and mutually intelligible dialects (Hunza and Nagar) and a third one (Yasin) that is significantly more divergent and allegedly harder to understand for Hunza and Nagar speakers, although Hunza and Yasin still share about 94% common lexicon on the Swadesh wordlist, indicating a period of divergence that is unlikely to exceed 1,000 years. The formal status of Burushaski as an "isolate" is most likely due to displacement and assimilation of the original speakers of this taxon in the area, mainly by speakers of various branches of Indo-Iranian, from Dardic to Eastern Iranian and Indo-Aryan (Urdu); the Burushaski language itself has numerous borrowings from these languages, although its basic lexical and grammatical structure still survives.

The close proximity of the dialects makes the special reconstruction of a "Proto-Burushaski" somewhat superfluous; nevertheless, due to a few phonetic shifts and some lexical replacements in daughter dialects it is technically possible, and a first attempt was carried out by S. Starostin (2005b), who also analyzed some peculiarities of Burushaski morphophonology, for instance, setting up a special Proto-Burushaski lateral cluster **lt* for stems containing a phoneme that is realized as *t-* in the word-initial position and as *-lt-* word-medially (e. g. **ltap* 'leaf' → Yasin *tap*, but *du=ltap-i-* 'to wither'; **lten* 'bone' → Hunza *tin*, but *=ltin* with possessive prefixes). Considering that root-initial clusters in Burushaski are otherwise strictly prohibited, we reinterpret this cluster as a monophonemic lateral affricate **ƛ*.

The Burushaski database (Starostin 2013b) consists of two closely related lects: Yasin and Hunza. The Proto-Burushaski (scil. Proto-Yasin-Hunza) reconstruction is self-evident for the majority of Swadesh items, the problematic cases being: 'dry', 'feather', 'fish', 'name', 'to swim', 'tail', 'to go', 'salt', see individual notes on these concepts. Eight Swadesh items are not reconstructible for Proto-Burushaski, since they are either not documented or replaced with recent borrowings: 'all', 'bark', 'good', 'round', 'tree'.

#### 6. Linguistic comments on individual Swadesh forms

All linguistic data in the present article are encoded in the unified transcription system of the *Global Lexicostatistical Database* project (http://starling.rinet.ru/new100/), which is generally based on the IPA alphabet, with several specific discrepancies, e.g., we use traditional *c č* for IPA *ts tʃ*. See http://starling.rinet.ru/new100/UTS.htm

1. 'all (*omnis*)'.

**Proto-Yeniseian**. **bɨʎ-* (S. Starostin 1995: 211), not very reliable, since the form is properly reconstructible only on the Ket-Yugh level.

**Proto-Athabaskan**. An unstable item; the match Kato *ɬe-neʔ-haʔ* 'all' / Tanaina *ɬu-q'u* 'all' provides us with a possible candidate: **ɬV-*, which is supported by the Eyak cognate (*ɬi-ʔq'* 'all').

**Eyak.** *ɬi-ʔq'*, cognate to Athabaskan.

**Proto-Athabaskan-Eyak**. **ɬV-* (Athabaskan, Eyak).

**Proto-Eskimo**. **tama-ʁ* (Fortescue et al. 2010: 358) is retained in both branches.

**Proto-Aleut**. **huzu-* (Bergsland 1994: 452; Golovko 1994: 193) is retained in Eastern and Atkan as 'all (*omnis*) / all (*totus*)'. In Attuan, superseded with *čimika-χ*, whose Common Aleut meaning is 'whole' (Bergsland 1994: 143).

**Proto-Chukotian**. **əmə-* (Fortescue 2005: 342), meaning 'all (*omnis*) / all (*totus*)'. Retained in all languages except for Alutor. Perhaps cognate to the Itelmen term.

**Proto-Itelmen**. **mini-ɬ* (Volodin, Khaloimova 2001: 140; Fortescue 2005: 342), meaning 'all (*omnis*) / all (*totus*)'. Perhaps cognate to the Chukotian term.

**Proto-Nivkh**. **ṭik* (Fortescue 2016: 32; Savelyeva, Taksami 1965: 82, 97).

**Proto-Samoyed**. **tük-* (Janhunen 1977: 168) is attested in Northern Samoyed and Mator.

**Proto-Yukaghir**. Kolyma **čömu*, also attested in old wordlists (Nikolaeva 2006: 139–140) vs. Tundra **yawnV-* (Nikolaeva 2006: 186). Kolyma word is compared by Nikolaeva to Tungusic **čuŋnu* 'all, entirely', but borrowing from Tungusic is improbable on phonetic grounds.

**Proto-Burushaski**. Superseded with Indo-Aryan loans.

2. 'ashes'.

**Proto-Yeniseian**. **qol- ~ *qor-* (S. Starostin 1995: 263). Of all known terms for 'ashes', only Ket *qɔlɨn* lacks any internal etymologization, and may therefore be tentatively regarded as the optimal candidate for Proto-Yeniseian 'ashes' at the moment.

**Proto-Athabaskan**. Not reconstructible reliably, because 'ashes' is normally expressed with help of various words for 'sand', 'dirt', etc., frequently with the epithet 'of fire'.

**Eyak.** *cʰĩʔƛ'-k*.

**Proto-Athabaskan-Eyak**. **cʰVnƛ'* (Eyak + scarcely retained Proto-Athabaskan **cʰiːƛ'* 'hot coals'), apparently related to Tlingit *kʰéɬ'-t'* 'ashes'. For the correspondences such as Athabaskan-Eyak **c* / Tlingit *k*, Leer (2010: 178) reconstructs a PAET velar-palatal row **kʸ, kʸʰ, kʸ', xʸ*, although G. Starostin (2012: 130) tends to reinterpret it as a specific sibilant row **cʸ, cʸʰ, cʸ', sʸ*.

**Proto-Eskimo**. **aʁða* (Fortescue et al. 2010: 45) is retained as 'ashes' in Inuit, having shifted into such specific meanings as 'gunpowder', 'medicine', etc. in Yupik.

**Proto-Aleut**. **utxi-χ* (Bergsland 1994: 453; Golovko 1994: 213) is retained at least in Eastern and Atkan.

**Proto-Chukotian**. **piŋ-piŋ* (Fortescue 2005: 216), retained as 'ashes' in all languages except for Kerek. Cognate to the Itelmen term.

**Proto-Itelmen**. **piŋ-piŋ* (Volodin, Khaloimova 2001: 72; Fortescue 2005: 216). Cognate to the Chukotian term (if not a Chukotian loan). Tends to be superseded with a derivative from the durative verb **čin-zu-* 'to be burnt *vel sim.*' (Volodin, Khaloimova 2001: 102, 156).

**Proto-Nivkh**. **bləŋk* (Fortescue 2016: 23; Savelyeva, Taksami 1965: 163; Savelyeva, Taksami 1970: 263). Attested as 'ashes' in Amur (thus Savelyeva & Taksami), East Sakhalin and South Sakhalin. The second candidate is **hilm-r* ~ **him-r* (Fortescue 2016: 74), but this one seems weaker. Firstly, because **hi(l)m-r* is glossed specifically as Russian 'пепел', i.e. 'fine ashes', for Amur (Savelyeva, Taksami 1965: 282; Savelyeva, Taksami 1970: 428) and only as 'soot' for North Sakhalin (Peiros, Starostin 1986: 215). Secondly, because **hi(l)m-r* is a transparent derivate from the verb **hil-m-* 'to blaze, flare' (Savelyeva, Taksami 1970: 428).

**Proto-Samoyed**. **kimä* (Janhunen 1977: 70) is attested in Northern Samoyed and Selkup.

**Proto-Yukaghir**. **noŋqə* (Nikolaeva 2006: 309) means 'ashes' in Tundra and 'sand' in Kolyma. Kolyma word for 'ashes' is a Russian loan.

3. 'bark'.

**Proto-Yeniseian**. **ʔɨɢ-* ~ **xɨɢ-* (S. Starostin 1995: 196). The sole uncontested candidate for Proto-Yeniseian 'bark', lost in Kott and not attested in either Arin or Pumpokol.

**Proto-Athabaskan**. **t'uːč'* is attested in the Northern and Apachean groups; Pacific Coast **sVc'* 'bark' < 'skin'.

**Eyak.** *qʰaht-ɬ*, synchronously this is the basic term for 'bark', but its complex morphological structure could point to a new formation; the obsolete term *ɬãh* 'bark', which represents a bare root, probably has more chance to be a Proto-Eyak word for this meaning.

**Proto-Athabaskan-Eyak**. **t'uːč'* (Athabaskan), **ɬaːn* (Eyak).

**Proto-Eskimo**. **qaʔəʁu-* (Fortescue et al. 2010: 301) can mean specifically 'birch bark' in some lects, nevertheless the generic meaning 'bark' is attested in both branches, e.g., in Central Siberian Yupik (Menovshchikov 1988: 178) and North Alaskan Inuit (MacLean 2014: 887). Morphologically unclear.

**Proto-Aleut**. **ukałaχ* (Bergsland 1994: 428), morphologically unclear, means 'bark' at least in Eastern and Atkan. In both dialect tends to be superseded by **qačχ(i)-* (Golovko 1994: 218; Bergsland & Dirks 1978: 143), whose original meaning is 'skin' (Bergsland 1994: 292).

**Proto-Chukotian**. **ut=qulɣə-n* (Fortescue 2005: 311), retained in all languages. A compound of **ut(tə)-* 'tree' (q.v.) and **qulɣə-n* 'fish skin' (Fortescue 2005: 241).

**Proto-Itelmen**. **ilʔal* (Volodin, Khaloimova 2001: 163; Fortescue 2005: 311).

**Proto-Nivkh**. **oʁm* (Fortescue 2016: 126; Savelyeva, Taksami 1965: 187).

**Proto-Samoyed**. **kasɒ* (Janhunen 1977: 65) is attested in Northern Samoyed, Kamass and Selkup.

**Proto-Yukaghir**. Kolyma **qaːr* (Nikolaeva 2006: 379) vs. Tundra **sawa* (Nikolaeva 2006: 399). Same word as 'skin', q.v.

**Proto-Burushaski**. Not documented.

4. 'belly'.

**Proto-Yeniseian**. Not reconstructible (all languages have different equivalents, mostly transparent new formations).

**Proto-Athabaskan**. **=wet'* is retained in all three branches.

**Eyak.** *=kʰəmah.*

**Proto-Athabaskan-Eyak**. **wet'* (Athabaskan), **kʰʷVnV* (Eyak).

**Proto-Eskimo**. **aqya-* (Fortescue et al. 2010: 44), retained in all branches.

**Proto-Aleut**. **kimla-χ* (Bergsland 1994: 238; Bergsland & Dirks 1978: 44; Golovko 1994: 208), attested in all branches, although tends to be superseded with *sanʁu-χ* 'stomach' (Bergsland 1994: 352) or *ili-ða-χ* < **ili-* 'inside' (Bergsland 1994: 191).

**Proto-Chukotian**. **nanqə-n* (Fortescue 2005: 185), retained in all languages.

**Proto-Itelmen**. **qʷeli-tq* (Volodin, Khaloimova 2001: 153).

**Proto-Nivkh**. **ɢoʁ* (Fortescue 2016: 68; Savelyeva, Taksami 1965: 141), meaning 'belly / stomach'.

**Proto-Samoyed**. It is hard to choose between **nancə̑* (Janhunen 1977: 20) and **pärkä* (Janhunen 1977: 122). **nancə̑* is retained in Northern Samoyed, Mator, Kamass and Selkup, **pärkä* – in Mator and Selkup (with derivatives in Northern Samoyed). The semantic difference between the two words in Mator and Selkup is difficult to establish.

**Proto-Yukaghir**. **lʸir-il* (Nikolaeva 2006: 243–244) is attested in both Yukaghir languages. A possible alternative candidate, **monʸ-il* (Nikolaeva 2006: 274), also attested in both languages, rather means 'stomach (as an internal organ)'.

5. 'big'.

**Proto-Yeniseian**. **qeʔ* (S. Starostin 1995: 300). In its original form and meaning, the word is well preserved in Ket-Yugh, as well as Pumpokol (where *xäː-se* = Ket *qɛː-sʸi* 'big; chief', substantivized form), but seems to be absent as such in Kott and Arin.

**Proto-Athabaskan**. **=kʰʸaːχ, *kʰʸuχ* is retained in all three branches.

**Eyak.** *=ʔluw ~ =ʔnuw.*

**Proto-Athabaskan-Eyak**. **=kʰaːχ* (Athabaskan), **nuw* (Eyak).

**Proto-Eskimo**. **aŋə-* (Fortescue et al. 2010: 35), retained in all branches. Cognate to the Aleut term.

**Proto-Aleut**. **aŋu-na-l-* (Bergsland 1994: 91; Golovko 1994: 186), attested in all branches. Cognate to the Eskimo term.

**Proto-Chukotian**. **mäγəŋ-* (Fortescue 2005: 171), retained in all languages.

**Proto-Itelmen**. **pəl-* (Volodin, Khaloimova 2001: 136; Stebnitsky 1934: 102; Fortescue 2005: 420).

**Proto-Nivkh**. **bil-* (Fortescue 2016: 23; Savelyeva, Taksami 1965: 71).

**Proto-Samoyed**. **ɒr-kɒ* (Janhunen 1977: 19) is retained in Northern Samoyed, Mator, Kamass and Selkup. The word is derived from **ɒrə̑* 'magnitude'.

**Proto-Yukaghir**. **čomo-* (Nikolaeva 2006: 138–139) is retained in both Yukaghir languages.

6. 'bird'.

**Proto-Yeniseian**. **duma* (S. Starostin 1995: 225). The word is found in both Ket-Yugh and Kott and is clearly of Proto-Yeniseian provenance, although its original semantics may have been specifically restricted to 'small bird'.

**Proto-Athabaskan**. Not reconstructible, because normally the concept 'bird' is expressed with various descriptive expression, e.g., 'small animal', 'the one with feathers' and so on.

**Eyak.** *yəχ=tə=ɬə=k'aʔt'-χ*, a descriptive formation 'it flies around'.

**Proto-Athabaskan-Eyak**. Not reconstructible.

**Proto-Eskimo**. **təŋ-miɣ-aʁ* (Fortescue et al. 2010: 372), attested with the generic meaning 'bird' in both Yupik and Inuit; derived from **təŋ-miɣ-* 'to be flying'.

**Proto-Aleut**. **sa-χ* (Bergsland 1994: 342; Golovko 1994: 257), attested in all branches.

**Proto-Chukotian**. **pəčiqä* (Fortescue 2005: 219), retained as a basic term in all languages except for Chukchi. Chukchi *ɣatle* 'bird' < 'duck' (Fortescue 2005: 82).

**Proto-Itelmen**. **unʸa-* (Volodin, Khaloimova 2001: 93; Ono 2003: 100; Fortescue 2005: 186), meaning 'small bird' or generally 'bird'.

**Proto-Nivkh**. **bəy-ŋa* (Fortescue 2016: 28; Savelyeva, Taksami 1965: 350). Derived from **bəy-* 'to fly' q.v., final **-ŋa* may be **ŋa* 'animal' (Fortescue 2016: 117).

**Proto-Samoyed**. **sɒrmɒ* (Janhunen 1977: 136) is a general word for 'bird' and '(wild) animal'. No specific word for 'bird' can be reconstructed.

**Proto-Yukaghir**. **nontə* (Nikolaeva 2006: 309) is a general word for 'bird' and '(wild) animal' in Proto-Yukaghir (cf. a similar polysemy in Samoyed). In Kolyma Yukaghir, the word is retained in the meaning 'bird'; the Tundra Yukaghir reflex of **nontə* means 'wolf', but the older meaning is preserved in compound *nodod-uo* 'egg' (lit. 'bird's child'). The modern Tundra Yukaghir word for 'bird' is a compound 'thing with wings'.

7. 'to bite'.

**Proto-Yeniseian**. Not reconstructible due to insufficient attestation.

**Proto-Athabaskan**. The main candidate is **=ʁVč ~ *=ʁVš* attested in all three branches.

**Eyak.** *=qʰa,* the Eyak paragim can point to either a PAE **CV:*-root or a **CV:n*-root.

**Proto-Athabaskan-Eyak**. **=χVč ~ *=χVš* (Athabaskan), **=qʰa: ~ *=qʰa:n* (Eyak).

**Proto-Eskimo**. **kəɣə-* (Fortescue et al. 2010: 179), retained in both Yupik and Inuit. Cognate to the Aleut term.

**Proto-Aleut**. **kix-s-* (Bergsland 1994: 238; Golovko 1994: 280), attested in all branches. Cognate to the Eskimo term.

**Proto-Chukotian**. **yəɣu-* (Fortescue 2005: 119), retained in all languages.

**Proto-Itelmen**. **pəl-k-* (Volodin, Khaloimova 2001: 73, 166; Fortescue 2005: 166).

**Proto-Nivkh**. **har- ~ *haz-* (Fortescue 2016: 71; Savelyeva, Taksami 1965: 435).

**Proto-Samoyed**. **sacʑ* (Janhunen 1977: 136–137) is retained in Northern Samoyed and Selkup.

**Proto-Yukaghir**. Kolyma **toð-* (Nikolaeva 2006: 432) vs. Tundra **nenč-* (Nikolaeva 2006: 296). The Tundra word has a Kolyma cognate with the meaning 'to gnaw'.

8. 'black'.

**Proto-Yeniseian**. **tum-* (S. Starostin 1995: 289). Preserved in all daughter languages.

**Proto-Athabaskan**. **=ʂe̢ɲ* is retained in all three branches. The initial retroflex points to PAE **xʷ*.

**Eyak.** *=t'u:č'*, cab be borrowed from Tlingit *t'u:č'* 'black'.

**Proto-Athabaskan-Eyak**. **=xʷe̢ɲ* (Athabaskan), **=t'u:č'* (Eyak).

**Proto-Eskimo**. **qiʁ-nəʁ-* (Fortescue et al. 2010: 336), attested with the basic meaning '(to be) black' in Inuit and in the derived stem 'blue fox' in Yupik, its antiquity may be proven by the potential Aleut cognate: **qax-čax-* 'to be black', although the correspondence Eskimo **ʁ* / Aleut **x* is not fully regular. In some Yupik languages, expressions for '(to be) black' are derived from **taʁəʁ-nəʁ* 'darkness' (Fortescue et al. 2010: 363).

**Proto-Aleut**. **qax-čax-s-* (Bergsland 1994: 295; Golovko 1994: 287), attested in all branches.

**Proto-Chukotian**. **əv-* (Fortescue 2005: 348), retained in all languages.

**Proto-Itelmen**. **ktɣə-* (Volodin, Khaloimova 2001: 221; Volodin 1976: 320; Fortescue 2005: 354).

**Proto-Nivkh**. **wəl-wəl* (Fortescue 2016: 164; Savelyeva, Taksami 1965: 453; Savelyeva, Taksami 1970: 58). Other Amur forms, not included into Fortescue 2016, such as **biw-* 'black' (Savelyeva, Taksami 1970: 262), seem more marginal.

**Proto-Samoyed**. Not reconstructible: none of the roots used in this meaning in various Samoyed languages can be safely projected on the Proto-Samoyed level.

**Proto-Yukaghir**. Kolyma **em-Vwə* (Nikolaeva 2006: 157–158) vs. Tundra **toro-* (Nikolaeva 2006: 436).

9. 'blood'.

**Proto-Yeniseian**. **sur* (S. Starostin 1995: 278). Preserved in all daughter languages (but not attested in Pumpokol). The same root also served (already on the Proto-Yeniseian level) as the main derivational stem for the word 'red' q.v.

**Proto-Athabaskan**. **tẹł* is retained in all three branches.

**Eyak.** *təł*, cognate to Athabaskan.

**Proto-Athabaskan-Eyak**. **tẹł* (Athabaskan, Eyak).

**Proto-Eskimo**. **aðuɣ* (Fortescue et al. 2010: 5), retained in all branches.

**Proto-Aleut**. **a:max ~ *a:mɣi-χ* (Bergsland 1994: 63; Golovko 1994: 220), attested in all branches.

**Proto-Chukotian**. **mullə-mul* (Fortescue 2005: 178), retained in all languages. Reduplicated stem. Cognate to the Itelmen term.

**Proto-Itelmen**. **mɫim* (Volodin, Khaloimova 2001: 165; Fortescue 2005: 178). Cognate to the Chukotian term.

**Proto-Nivkh**. There are two candidates. First, **ŋar* (Fortescue 2016: 121; Savelyeva, Taksami 1965: 191; Savelyeva, Taksami 1970: 227), attested in Amur and North Sakhalin, meaning 'blood'. Second, **t̪oʁ* (Fortescue 2016: 35; Savelyeva, Taksami 1970: 227), meaning 'blood' in East and South Sakhalin, but 'juice, sap' in Amur and North Sakhalin. The root **ŋar* has the advantage because, firstly, it is possible that **ŋar* is retained in East Sakhalin in a derivative which means 'vein' (thus Taksami 1996: 116); secondly, the meaning 'sap' or 'resin' is still documented for **t̪oʁ* in some South Sakhalin sources.

**Proto-Samoyed**. **kəm* (Janhunen 1977: 65) is retained in all Samoyed languages.

**Proto-Yukaghir**. Kolyma **lepk-ul* (Nikolaeva 2006: 240–241) vs. Tundra **če:mə* (Nikolaeva 2006: 129). Cf. also the root **minčə* ~ **minčʸə* 'blood, bloody', attested in several old wordlists (Nikolaeva 2006: 269).

10. 'bone'.

**Proto-Yeniseian**. **ʔaʔd* ~ **xaʔd* (S. Starostin 1995: 178). Preserved in Ket-Yugh; not attested in Arin and Pumpokol. In Kott, the etymological parallel is *ar-aŋ* ~ *ar-aŋ-an* 'joint; limb', which may be analyzed as a former collective plural form: 'limb' = '(a set of) bones'.

**Proto-Athabaskan**. **c'en* is retained in all three branches.

**Eyak.** *c'əl*, cognate to Athabaskan.

**Proto-Athabaskan-Eyak**. **c'en* (Athabaskan, Eyak).

**Proto-Eskimo**. **nənə-ʁ* (Fortescue et al. 2010: 248) means 'bone' in Yupik and Sirenik, without Inuit cognates. The second candidate is **caHu-nəʁ* (Fortescue et al. 2010: 78) which means 'bone' in Inuit, not attested in Yupik; this one looks like a recent deverbative with the common nominalizer **-nəʁ*.

**Proto-Aleut**. **qaɣna-χ* (Bergsland 1994: 296; Golovko 1994: 219), attested in all branches.

**Proto-Chukotian**. **qətʁəm* (Fortescue 2005: 248), retained in all languages. Cognate to the Itelmen term.

**Proto-Itelmen**. **ktχʷəm* (Volodin, Khaloimova 2001: 164; Fortescue 2005: 248). Cognate to the Chukotian term.

**Proto-Nivkh**. **ŋa=ɲɣəv* (Fortescue 2016: 121; Savelyeva, Taksami 1965: 189). A prefixal element **ŋa-*, common for body part names, can be singled out.

**Proto-Samoyed**. **lɘ* (Janhunen 1977: 82), retained in all Samoyed languages, goes back to Proto-Uralic **lɨwi* 'bone'.

**Proto-Yukaghir**. **am-un* (Nikolaeva 2006: 102) is retained in both Yukaghir languages.

11. 'breast (generic or male), chest'.

**Proto-Yeniseian**. Ket-Yugh & Pumpokol **təga* (S. Starostin 1995: 284) is distinctly opposed to the Kott-Arin isogloss, reconstructible with difficulty: Kott *pa* and Arin *pʰi-* are hard to reconcile; perhaps the vowel fluctuation is due to different ways of contraction of an earlier cluster, e.g., < **paxV* or **pixV*. Which of the two should be considered the primary candidate for Proto-Yeniseian '(male) breast', remains uncertain; we include both into comparison.

**Proto-Athabaskan**. **č̨V:χ* is retained in all three branches at least in relic expressions. PA retroflex points to PAE **kʷ*.

**Eyak.** *=še:k'*.

**Proto-Athabaskan-Eyak**. **kʷV:χ* (Athabaskan), **še:k'* (Eyak).

**Proto-Eskimo**. **qatə-ɣ* (Fortescue et al. 2010: 316) means 'human breast, chest' in Yupik and Sirenik, having shifted into the meaning 'breastbone of bird' in Inuit. In the Inuit languages, 'chest' is expressed with either a new suffixed derivation from **qatəɣ* or a suffixed derivation from **caðə-* 'front of body' (Fortescue et al. 2010: 67).

**Proto-Aleut**. **simsi-* (Bergsland 1994: 361; Golovko 1994: 106, 199), attested in all branches. The second and more marginal candidate is **kači-χ* (Bergsland 1994: 220).

**Proto-Chukotian**. **mačve* (Fortescue 2005: 168), retained in all languages, frequently with suffix extensions.

**Proto-Itelmen**. Inherited terms are not documented reliably. Cf. the Chukotian loan *wayeter* ‘chest’ (Fortescue 2005: 322).

**Proto-Nivkh**. **ŋa=rɣər* (Fortescue 2016: 121; Savelyeva, Taksami 1965: 121; Savelyeva, Taksami 1970: 237). A prefixal element **ŋa-*, common for body part names, can be singled out.

**Proto-Samoyed**. **sünsə̆* (Janhunen 1977: 144), retained in Northern Samoyed and Mator, has an external cognate in Hungarian and goes back to Proto-Uralic **sʸünsʸi*.

**Proto-Yukaghir**. Kolyma **mel-uð* (Nikolaeva 2006: 263) vs. Tundra **sis-il* (Nikolaeva 2006: 407). **mel-uð* has more chances to be the Proto-Yukaghir word for ‘breast’, since it is attested in fragmentarily known Omok, a language sharing some important phonological innovations with Tundra Yukaghir. Still, we list both words.

12. ‘to burn (trans.)’.

**Proto-Yeniseian**. **qɔʔt* (S. Starostin 1995: 304). In the Kott-Arin branch, this word is preserved only as a nominal stem ‘fire’, where it has wiped out the original root for ‘fire’ (**boʔk* q.v.), although the exact situation in Arin is actually unknown (no equivalent for the verb ‘to burn’ attested in that branch).

**Proto-Athabaskan**. **=q’aːn* is retained in all three branches.

**Eyak.** *=q’a*, the Eyak paragim can point to either a PAE **CVː*-root or a **CVːn*-root.

**Proto-Athabaskan-Eyak**. **=q’aːn* (Athabaskan, Eyak), cognate to Tlingit *χ’aːn* ‘fire’.

**Proto-Eskimo**. **əkə-* (Fortescue et al. 2010: 110) means ‘to burn (intr.)’ in both Yupik and Inuit, the transitive stem ‘to burn (trans.)’ is derived with help of perfective/causative suffixes: usually **-uma-* in Yupik-Sirenik, **-t-* in Inuit. In some Yupik lects and Sirenik, ‘to burn (trans.)’ is expressed with the verb **pinə-ʁ-* ‘to fry out’ (Fortescue et al. 2010: 272; Menovshchikov 1988: 237).

**Proto-Aleut**. **hix-t-* (Bergsland 1994: 175), causative from **hix-* ‘to burn (intr.)’; cognate to the Aleut term, if *h-* is secondary. In Atkan, the transitive stem tends to be superseded with *a-ta-* ‘to burn (trans.)’ (Golovko 1994: 35) < **a-* ‘to blaze’ (Bergsland 1994: 5).

**Proto-Chukotian**. **ðən=känɣ-* (Fortescue 2005: 132) is a basic verb for ‘to burn (tr.)’ in Koryak (Zhukova 1990: 198) and Alutor (Kibrik et al. 2004: 421), derived from intransitive **känɣ-* ‘to burn (in a specific way)’ with the transitivizer **ðən-*. In Chukchi, the transitive stem **ðən=tləv-ät-* ‘to burn’ (Fortescue 2005: 69) is used < **təlvə-* ‘to burn (oneself)’ (Fortescue 2005: 296). Distinct from the basic Proto-Chukotian intransitive verb **ŋəl-ät-* ‘to burn’ (Fortescue 2005: 201).

**Proto-Itelmen**. **ən=q’wa-* (Volodin, Khaloimova 2001: 112, 153). Causative in **ən-* from intransitive **q’wa-* ‘to burn’. The basic verb for ‘to burn (intr.)’ is apparently **lu-* (Volodin, Khaloimova 2001: 47, 146; Fortescue 2005: 297).

**Proto-Nivkh**. **tuv-* (Fortescue 2016: 152; Savelyeva, Taksami 1965: 386; Savelyeva, Taksami 1970: 323). This is the basic verb for ‘to burn (tr.)’ at least in Amur. Distinct from **u-* ‘to burn (intr.)’ (Fortescue 2016: 155; Savelyeva, Taksami 1965: 118; Savelyeva, Taksami 1970: 392).

**Proto-Samoyed**. **kɒrə̆-* (Helimski 1997: 268–269) is attested only in Mator, but has an external etymology. According to Helimski, it can be compared to Proto-Uralic **karti-* ‘to roast, to burn’. Other roots, used in this meaning in Samoyed languages, cannot be safely projected to the Proto-Samoyed level.

**Proto-Yukaghir**. Kolyma **pentə-* (Nikolaeva 2006: 349–350) vs. Tundra **ent-* (Nikolaeva 2006: 162).

13. ‘fingernail’.

**Proto-Yeniseian**. **ʔiːɲe ~ *xiːɲe* (S. Starostin 1995: 195). Preserved only in Ket-Yugh. In Kott, probably replaced by *halčiːg* ‘hoof’ = Ket *qɔlʸesʸ*, Arin *kalɨs* ‘hoof’.

**Proto-Athabaskan**. **qVɲ* is retained in all three branches.

**Eyak.** *=yə=ɬ=χahc-ɬ*, initial *-yə-* means ‘hand’, final *-ɬ* is a common nominal suffix (originally instrumental).

**Proto-Athabaskan-Eyak**. We follow Leer 2010: 183 and treat Eyak *χahc* as an etymological cognate of Tlingit *χaːkʷ* ‘fingernail’. According to Leer 2008a, vowel aspiration in Eyak points to a nasal cluster, so the Proto-Athabaskan-Eyak-Tlingit form for ‘fingernail’ should be something like **χankʸ* > Proto-Athabaskan-Eyak **χanc* ‘fingernail’ (Proto-Athabaskan **qVɲ* ‘fingernail’ is thus an innovation of unclear origin).

**Proto-Eskimo**. **kuki-ɣ* (Fortescue et al. 2010: 197), meaning ‘nail, claw, hoof, a claw-like tool (e.g., stone head of scraper)’ in Inuit. In Yupik, the root is only retained in the suffixed stem **kukiɣ-kšak* which denotes a claw-like tool: ‘greenish stone used as chisel for ivory carving’, ‘grapnel hook’. In Yupik-Sirenik, ‘fingernail’ is expressed by **citu-ɣ* (Fortescue et al. 2010: 94), lacking Inuit cognates.

**Proto-Aleut**. **qaɣa-lʁi-χ* (Bergsland 1994: 295; Golovko 1994: 233), attested in all branches. Perhaps derived from **qaɣa-* ‘to knock, crack, make a sharp noise’ (Bergsland 1994: 294).

**Proto-Chukotian**. **väɣ* (Fortescue 2005: 313), retained in all languages.

**Proto-Itelmen**. **k’uxʷ-k’uxʷ* (Volodin, Khaloimova 2001: 178; Fortescue 2005: 363), a reduplicated stem.

**Proto-Nivkh**. **dak-ɲ* (Fortescue 2016: 38; Savelyeva, Taksami 1965: 243; Savelyeva, Taksami 1970: 367). Derived from **dak-* ‘to carve, make cuts in’ (Fortescue 2016: 37), see Panfilov 1962: 61 for the rare suffix **-ɲ*.

**Proto-Samoyed**. **kə̆tɒ* (Janhunen 1977: 55–56) is retained in all Samoyed languages.

**Proto-Yukaghir**. **önčʸ-il* (Nikolaeva 2006: 330) is retained in both Yukaghir languages.

14. ‘cloud’.

**Proto-Yeniseian**. **ʔas=pur* (S. Starostin 1995: 255). Preserved in all daughter languages (although not attested in Pumpokol). Structure-wise, the word is clearly a compound, in which the first part is the Proto-Yeniseian word for ‘sky’: Ket *eˑsʸ*, Yugh *es*, Kott *eːš*, Arin *es*, Pumpokol *eč* < Proto-Yeniseian **ʔes* (S. Starostin 1995: 188).

**Proto-Athabaskan**. Apachean-Northern **q’Vs* competes with pacific Coast **ʔVχ*. The external Eyak cognate (*q’ahs* ‘cloud’) speaks in favor of **q’Vs*.

**Eyak.** *q’ahs*, cognate to Athabaskan.

**Proto-Athabaskan-Eyak**. **q’ʷVns* (Athabaskan, Eyak). We follow Leer 2008a and reconstruct a medial nasal on account of vowel aspiration in Eyak.

**Proto-Eskimo**. **qilaɣ-ɬuʁ* (Fortescue et al. 2010: 331), literally ‘bad sky’ < **qilaɣ* ‘sky’ + **-ɬuɣ* ‘bad’. Formally this is the best candidate, because **qilaɣ-ɬuʁ* means ‘cloud’ everywhere in Yupik-Sirenik and in a dialect of Seward Peninsula Inuit. Another candidate is **nuviya-* (Fortescue et al. 2010: 266), which means ‘cloud’ everywhere in Inuit, lacking Yupik cognates.

**Proto-Eskimo**. **qilaɣ-ɬuʁ* (Fortescue et al. 2010: 331), literally ‘bad sky’ < **qilaɣ* ‘sky’ + **-ɬuɣ* ‘bad’. Formally this is the best candidate, because **qilaɣ-ɬuʁ* means ‘cloud’ everywhere in Yupik-Sirenik and in a dialect of Seward Peninsula Inuit. Another candidate is **nuviya-* (Fortescue et al. 2010: 266), which means ‘cloud’ everywhere in Inuit, lacking Yupik cognates. The Seward Peninsula Inuit form may be a contact-driven innovation, so **nuviya-* has a better chance to present a Proto-Eskimo term than **qilaɣ-ɬuʁ* which looks like a new formation. We treat both **qilaɣ-ɬuʁ* and **nuviya-* as synonyms.

**Proto-Aleut**. **inka-maːʁu-χ* (Bergsland 1994: 202; Golovko 1994: 234), attested in all branches. Derived from **inka-χ* ‘sky’ + **-maːʁu- ~ -miːʁu-* ‘?’.

**Proto-Chukotian**. **yəʁə-n* (Fortescue 2005: 124), retained in all languages.

**Proto-Itelmen**. **mizʸə-* (Fortescue 2005: 357), note the secondary initial *ŋ-* for expected *m-* in modern Western (Volodin, Khaloimova 2001: 65).

**Proto-Nivkh**. **laɣ* (Fortescue 2016: 92; Savelyeva, Taksami 1965: 248; Savelyeva, Taksami 1970: 157). Attested everywhere, although is East Sakhalin tends to be superseded with *čʰarŋi* 'cloud' (Savelyeva, Taksami 1970: 157) of unclear origin.

**Proto-Samoyed**. **tiə̆* (Janhunen 1977: 162) is retained in all Samoyed languages.

**Proto-Yukaghir**. Kolyma **qa:r* (Nikolaeva 2006: 379) vs. Tundra **sawa* (Nikolaeva 2006: 399). In Tundra Yukaghir, the meaning 'cloud' is expressed by the compound 'skin of the sky'; Kolyma Yukaghir has a compound of 'skin' with an otherwise unknown root (an old word for 'sky'?). The main word is thus identical to 'skin', q.v.

15. 'cold'.

**Proto-Yeniseian**. It is impossible to choose a single candidate for the meaning 'cold' in Proto-Yeniseian, since at least two choices have the exact same probability: Ket-Yugh **taʔy* (S. Starostin 1995: 280) and Kott-Arin **ǯVr₁- ~ *ǯVl-* (S. Starostin 1995: 311). The vocalism in the case of the latter is hard to recover due to morphological vowel gradation in the attested forms.

**Proto-Athabaskan**. **q'ʷVc'* is retained in all three branches.

**Eyak.** *=ƛ'e*, the Eyak paragim can point to either a PAE **CV:*-root or a **CV:n*-root.

**Proto-Athabaskan-Eyak**. **q'ʷVc'* (Athabaskan), **=ƛ'V ~ *=ƛ'Vn* (Eyak).

**Proto-Eskimo**. An unstable and not very reliably documented item. The best candidates are various derivatives from **nəŋə-* 'to be(come) cold' (Fortescue et al. 2010: 249) and **itðə-* 'cold' (Fortescue et al. 2010: 160), both roots are attested in Yupik and Inuit. It is likely that **nəŋə-* is typically applicable to weather, whereas **itðə-* normally means 'cold (of objects)', but it cannot be established with certainty without detailed synchronous descriptions. We treat both roots as synonyms.

**Proto-Aleut**. **qiŋa-na-* (Bergsland 1994: 325; Golovko 1994: 82, 284), attested in Eastern and Atkan, applicable to objects and weather, derived from the substantive **qiŋa-* 'cold'. Distinct from **aču-na-* 'cold (of weather)' attested in Atkan and Attuan (Bergsland 1994: 8).

**Proto-Chukotian**. **čəq-* (Fortescue 2005: 53), apparently a basic term at least in Chukchi (Inenlikei 2005), Koryak (Zhukova 1990: 65, 215) and Alutor (Kibrik et al. 2004: 523).

**Proto-Itelmen**. **ɬqa-* (Volodin, Khaloimova 2001: 47, 219; Fortescue 2005: 166).

**Proto-Nivkh**. **div- ~ *tiv-* (Fortescue 2016: 148; Savelyeva, Taksami 1965: 448; Savelyeva, Taksami 1970: 353). Applicable to both objects and weather.

**Proto-Samoyed**. **täksV- ~ *tätsV- ~ *täcṣV- ~ *tässV-* (Janhunen 1977: 159) is retained in all Samoyed languages.

**Proto-Yukaghir**. Kolyma **čel-* (Nikolaeva 2006: 128) vs. Tundra **qančʸ-* (Nikolaeva 2006: 377–378). **čel-* is also attested in Omok. Kolyma reflex of **qančʸ-* means 'to temper'. Since Yukaghir **qančʸ-* is borrowed from Proto-Samoyed **kə̆nsä-* 'to cool down' we do not include it in the Proto-Yukaghir list.

16. 'to come'.

**Proto-Yeniseian**. The Proto-Yeniseian form is not very well reconstructible, since the Ket-Yugh and Kott forms lack mutual etymologization. We may settle upon **=ət-* (preserved in Kott) as the best candidate, because of a highly non-trivial morphophonological structure of the Kott paradigm that can speaks in favor of its archaic nature. In the Ket-Yugh branch, the root **i-* is used as an element of the Ket-Yugh complex verb 'to come', but the secondary morphological nature of the Ket-Yugh expression makes **i-* a less probable candidate for the Proto-Yeniseian status.

**Proto-Athabaskan**. **=ha:* 'to go / to come' is retained in all three branches.

**Eyak.** *=a*, cognate to Athabaskan.

**Proto-Athabaskan-Eyak**. **=ha:* (Athabaskan, Eyak).

**Proto-Eskimo**. There are two equally probable candidates. The first one is **aɣə-ɣiʁ-* (Fortescue et al. 2010: 7) which means 'to come' in Inuit. In Yupik, only the additionally suffixed stem is retained with the meaning 'to approach from a distance, return, pass'. An archaic derivative from Eskimo **aɣə-* 'to go (over or past)'. The second candidate is **taɣi-* (Fortescue et al. 2010: 354), meaning 'to come' in Yupik. We treat **aɣə-ɣiʁ-* and **taɣi-* as synonyms.

**Proto-Aleut**. **haqa-l-* (Bergsland 1994: 93; Golovko 1994: 158), attested in all branches.

**Proto-Chukotian**. **yät-* (Fortescue 2005: 112), retained in all languages.

**Proto-Itelmen**. **k'oɬ-* (Volodin, Khaloimova 2001: 194; Fortescue 2005: 358).

**Proto-Nivkh**. **prə-* (Fortescue 2016: 137; Savelyeva, Taksami 1965: 334; Savelyeva, Taksami 1970: 291).

**Proto-Samoyed**. **toy-* ~ **tuy-* (Janhunen 1977: 164), retained in all Samoyed languages, goes back to Proto-Uralic **tuli-* 'to come'.

**Proto-Yukaghir**. **kel-* (Nikolaeva 2006: 205) is retained in both Yukaghir languages.

17. 'to die'.

**Proto-Yeniseian**. **qɔ* (S. Starostin 1995: 264). Preserved in all daughter languages.

**Proto-Athabaskan**. Not reconstructible, because normally the concept 'to die' is expressed with various polite and euphemistic expressions ('to happen', to sleep' etc.).

**Eyak.** *=sĩh.*

**Proto-Athabaskan-Eyak**. **=sVn* (Eyak).

**Proto-Eskimo**. **tuqu-* (Fortescue et al. 2010: 386), retained in all branches.

**Proto-Aleut**. **asχa-l-* (Bergsland 1994: 99; Golovko 1994: 281), attested in all branches.

**Proto-Chukotian**. **viʁ-* (Fortescue 2005: 318), retained in all languages.

**Proto-Itelmen**. **izʸʔa-* (Volodin, Khaloimova 2001: 217; Fortescue 2005: 359).

**Proto-Nivkh**. **mu-* (Fortescue 2016: 108; Savelyeva, Taksami 1965: 435; Savelyeva, Taksami 1970: 196). Cf. the homonymous verb **mu-* 'to become' (Fortescue 2016: 108), thus the Common Nivkh meaning 'to die' could be a result of polite usage.

**Proto-Samoyed**. **kɒə̆-* (Janhunen 1977: 56–57), retained in all Samoyed languages, goes back to Proto-Uralic **kali-* 'to die'.

**Proto-Yukaghir**. **yompə-* (Nikolaeva 2006: 194) is attested in modern Tundra Yukaghir as well as in several old wordlists. Connection with Kolyma *you* 'disease', accepted by Nikolaeva, is phonologically irregular. Modern Kolyma *amdə-* 'to die' is related to words meaning 'to lay down' etc. (Nikolaeva 2006: 102).

18. 'dog'.

**Proto-Yeniseian**. **čip*. Preserved in all daughter languages. The Kott-Arin forms are attested in conjunction with a desemanticized prefix (Kott *al=*, Arin *il=*, original vocalism unclear) that is also encountered in several other entries on the Swadesh wordlist ('bird', 'star'); this seems to have been a shared Kott-Arin innovation.

**Proto-Athabaskan**. **łeɲ* is retained in all three branches.

**Eyak.** *χəwa:*, morphologically unclear, looks like a secondary formation.

**Proto-Athabaskan-Eyak**. **łeɲ* (Athabaskan).

**Proto-Eskimo**. **qikmi-ʁ* (Fortescue et al. 2010: 331), retained as a basic term in both Yupik and Inuit.

**Proto-Aleut**. **ayku-χ* (Bergsland 1994: 120), attested in Eastern and Atkan.

**Proto-Chukotian**. **qətʁə-n* (Fortescue 2005: 247), retained in all languages. Cognate to the Itelmen term.

**Proto-Itelmen**. **qosχ* (Volodin, Khaloimova 2001: 207; Fortescue 2005: 248). Cognate to the Chukotian term.

**Proto-Nivkh**. **ɢa-nŋ* (Fortescue 2016: 65; Savelyeva, Taksami 1965: 396; Savelyeva, Taksami 1970: 140). As proposed in Panfilov 1962: 51, the rare suffix **-nŋ* is to be singled out on the basis of the cognate Amur form *qa-χ* ‘leading dog of a team in dog races at the bear festival’, further cf. **wo* ‘village’ / **wo-nŋ* ‘villager’ (Fortescue 2016: 163).

**Proto-Samoyed**. **wǝn* (Janhunen 1977: 173–174) is retained in all daughter languages. This word is possibly a loan from Tocharian (cf. Tocharian B ⟨kweṃ⟩ ‘dog’, acc.sg.).

**Proto-Yukaghir**. Kolyma **pumpə-l* (Nikolaeva 2006: 370) vs. Tundra **qapnʸə* (Nikolaeva 2006: 379). Words for ‘dog’ in both modern Yukaghir languages are innovations. Modern Kolyma *towkə* (earlier *towoka, toboko, tabaka*) is borrowed from Russian *sobaka* ‘dog’ (Nikolaeva 2006: 408). Jochelson (1926: 326) lists Kolyma *pubel* ‘dog’ with the note: “ancient word, now *toboko*”. The same word is attested in several old wordlists. It seems clear that **pumpə-l* is the original Kolyma word for ‘dog’, replaced by a Russian borrowing. The word *qapnʸe*, attested in an old wordlist as a word for ‘dog’, is used in modern Tundra as a curse and as a part of compound *qapnʸe-burie* ‘currants’ (lit. ‘dog’s berry’). It must be noted that the English gloss ‘infection, contagion (also used as a word of abuse)’ in (Nikolaeva 2006: 379) is a mistake: it is explicitly stated in (Kurilov 2001: 249), the source of Tundra data in (Nikolaeva 2006), that the word is used only as a mild swear-word. Kurilov glosses it by the Russian swear-word *зараза*, which literally means ‘contagion’: this polysemy is nowhere attested in Yukaghir. The origin of Modern Tundra *laːma* ‘dog’ (Nikolaeva 2006: 232) is not known, but it is clear that this word replaced an earlier word for ‘dog’, *qapnʸe*.

19. ‘to drink’.

**Proto-Yeniseian**. **=op* (S. Starostin 1995: 202). Best attested in Kott, as well as in a part of the Ket-Yugh paradigm.

**Proto-Athabaskan**. **=naːŋ* is retained in all three branches; we interpret Krauss & Leer’s (1981: 70) *$ŋ_2$ as **ŋ*.

**Eyak.** *=tə=la*, the Eyak paragim can point to either a PAE **CVː*-root or a **CVːn*-root.

**Proto-Athabaskan-Eyak**. **=naːŋ* (Athabaskan, Eyak), cognate to Tlingit *=naː* ‘to drink’, but details are not entirely clear.

**Proto-Eskimo**. **əmə-ʁ-* (Fortescue et al. 2010: 120), retained in all branches, the same stem as ‘water’ (q.v.).

**Proto-Aleut**. **taːŋa-χ* (Bergsland 1994: 392; Golovko 1994: 244), attested in all branches, the same stem as ‘water’ (q.v.).

**Proto-Chukotian**. **iw=ɣiči-* (Fortescue 2005: 105), retained in all languages, except for Kerek. The second candidate is **pəl-* (Fortescue 2005: 221), whose meaning is rather to be reconstructed as ‘to drink up’.

**Proto-Itelmen**. **ɣil-* (Volodin, Khaloimova 2001: 186; Fortescue 2005: 360).

**Proto-Nivkh**. **da-* (Fortescue 2016: 37; Savelyeva, Taksami 1965: 294).

**Proto-Samoyed**. **ǝ-r-* (Janhunen 1977: 21–22), retained in Nenets, Mator and Selkup, goes back to Proto-Uralic **yɨγi-* ‘to drink’. Replaced in some daughter languages by **witV-*, derived from **wit* ‘water’.

**Proto-Yukaghir**. Kolyma **ončə-* (Nikolaeva 2006: 330–331) vs. Tundra **law-* (Nikolaeva 2006: 236). Cf. also the root **liŋkə-* ‘to drink’ (Nikolaeva 2006: 255), attested in Chuvan and Omok wordlists.

20. ‘dry’.

**Proto-Yeniseian**. **qɔy-* ~ **qɔɢ-* (S. Starostin 1995: 265). Preserved in all daughter languages except for the Kott dialect described by Castrén.

**Proto-Athabaskan**. Two roots compete with each other: **=qaɲ* and **=cʰaːɣʸ* / **=cʰaːk’ʸ*.

**Eyak.** *=tə=ɬ=ʔeht.*

**Proto-Athabaskan-Eyak**. **=qaɲ*, **=cʰaːk’* (Athabaskan), **=ʔVnt* (Eyak).

**Proto-Eskimo**. An unstable and not very reliably documented item. The best candidate is **kinə-ʁ-* (Fortescue et al. 2010: 191), which means ‘to be(come) dry’ in Yupik and ‘to filter out’, ‘to melt away’, ‘to have run off (water)’ in Inuit. In Inuit, the meaning ‘(to be) dry’ can be expressed with *pali-ʁ-* 'to be tanned by the sun, sunburnt' (Fortescue et al. 2010: 271) or **panə-ʁ* 'to dry out, starve' (Fortescue et al. 2010: 272).

**Proto-Aleut**. **qaka-* (Bergsland 1994: 300; Golovko 1994: 273), attested in all branches.

**Proto-Chukotian**. **kərɣə-* (Fortescue 2005: 150), retained as basic 'dry' in all languages except for Kerek. Cognate to the Itelmen term.

**Proto-Itelmen**. **k'izʸɣi-* (Volodin, Khaloimova 2001: 211; Fortescue 2005: 151). Cognate to the Chukotian term.

**Proto-Nivkh**. There are two candidates. First, **ṭe-* ~ **te-* (Fortescue 2016: 31; Savelyeva, Taksami 1965: 414; Savelyeva, Taksami 1970: 446), meaning '(to be) dry' in Amur and perhaps South Sakhalin. Second, **qaw-* (Fortescue 2016: 140; Savelyeva, Taksami 1970: 446), meaning '(to be) dry' in East and North Sakhalin. We are forced to treat them as synonyms.

**Proto-Samoyed**. **kɒsə̆-* (Janhunen 1977: 60–61), retained in Northern Samoyed and Mator, goes back to Proto-Uralic **kosʸki* 'dry'.

**Proto-Yukaghir**. Kolyma **keːlə-* (Nikolaeva 2006: 204) vs. Tundra **sɨl-* (Nikolaeva 2006: 421–422). Tungusic **sile-*, compared to Yukaghir **sɨl-* in (Nikolaeva 2006: 422), means 'dew', not 'dry'.

**Proto-Burushaski**. In both Yasin and Hunza dialects the adjectival meaning 'dry' is expressed by forms derived from the Common Burushaski verb **buɻ-* 'to be dry / to be thirsty', but morphological models are not identical between the two dialects, making it likely that we are dealing with parallel new formations in both cases. Instead, we fill the slot with **qaq-* which is an alternate Yasin term for 'dry / thirsty' (synchronic difference between the two Yasin words for 'dry' is unclear); in Hunza it is only attested with the meaning 'hungry'. Thus the most likely scenario is that **qaq-* meant 'dry (adj.) / thirsty' in Proto-Burushaski as opposed to the verb **buɻ-* 'to be dry / to be thirsty'; in Yasin, **qaq-* is still retained with its original meaning 'dry / thirsty', competing with new deverbal formations from **buɻ-* 'to be dry / to be thirsty'; in Hunza, **qaq-* was superseded with the verb **buɻ-* and has itself shifted to the meaning 'hungry'.

21. 'ear'.

**Proto-Yeniseian**. **ʔɔdɢe* (S. Starostin 1995: 198). Preserved in all daughter languages except for Kott (*kaloːx*, most likely borrowed from a Turkic source). The reconstruction shape is somewhat problematic; however, the reconstruction **ʔɔdɢe*, despite the uniqueness of its medial cluster, accounts for most of the resulting diversity in daughter languages (Ket *ɔgdɛ*, Yugh *ɔχtiŋ*, Arin *utqʸöːn-oŋ*, Pumpokol *atkin*).

**Proto-Athabaskan**. **čeχ* is retained in all three branches.

**Eyak.** *=čehχ*, cognate to Athabaskan.

**Proto-Athabaskan-Eyak**. **čVnχ* (Athabaskan, Eyak). We follow Leer 2008a and reconstruct a medial nasal on account of vowel aspiration in Eyak.

**Proto-Eskimo**. **ciɣ-un* (Fortescue et al. 2010: 82), retained in all branches. Deverbative form an unattested verb with the instrumental suffix *-*un*.

**Proto-Aleut**. **tut-usi-χ* (Bergsland 1994: 411; Golovko 1994: 283), attested in all branches. Derived from **tut-* 'to hear' (q.v.) with the instrumental suffix *-*Vsi*.

**Proto-Chukotian**. **vilu* (Fortescue 2005: 317), retained in all languages. Cognate to the Itelmen term.

**Proto-Itelmen**. **elwe-* (Volodin, Khaloimova 2001: 218; Fortescue 2005: 317). Cognate to the Chukotian term.

**Proto-Nivkh**. **noz* (Fortescue 2016: 112; Savelyeva, Taksami 1965: 440; Savelyeva, Taksami 1970: 213), attested in Amur and North Sakhalin. In East and South Sakhalin, it was superseded with **m-la* 'ear' (Fortescue 2016: 106; Taksami 1996: 52), which is likely to be derived from **mə-* 'to hear / to listen' q.v.

**Proto-Samoyed**. **kɒw* (Janhunen 1977: 62) is retained in all Samoyed languages.

**Proto-Yukaghir**. **unəmə* (Nikolaeva 2006: 444) is retained in both Yukaghir languages.

22. 'earth (soil)'.

**Proto-Yeniseian**. **baʔŋ* (S. Starostin 1995: 205). Preserved in all daughter languages.

**Proto-Athabaskan**. Two roots compete with each other: **ɲenʔ* and **ɬVč*. The external Eyak cognate (*ʔãh* 'earth') speaks in favor of **ɲenʔ*.

**Eyak.** *ʔãh*, cognate to Athabaskan with **nʸ > 0*.

**Proto-Athabaskan-Eyak**. **ɲVn* (Athabaskan, Eyak).

**Proto-Eskimo**. **nuna* (Fortescue et al. 2010: 262), retained in Yupik and Inuit, meaning 'soil / land / ground'. Cognate to the Aleut term.

**Proto-Aleut**. **tana-χ* (Bergsland 1994: 388; Golovko 1994: 119, 212), attested in all branches, meaning 'soil / land / ground'. Cognate to the Eskimo term.

**Proto-Chukotian**. **nutä-lq-ən* (Fortescue 2005: 189), retained in all languages with polysemy 'soil / ground', derived from **nutä* 'land'.

**Proto-Itelmen**. **ktxu-m* ~ **kčtxu-m* (Volodin, Khaloimova 2001: 35, 157; Fortescue 2005: 361), meaning specifically 'soil'. Distinct from **sʸimt* 'land' (Volodin, Khaloimova 2001: 157; Fortescue 2005: 373), which can, however, acquire the meaning 'soil' in some lects, thus Volodin 1976: 105, 153, 169; Ono 2003: 98.

**Proto-Nivkh**. **miv* (Fortescue 2016: 105; Savelyeva, Taksami 1965: 162, 325; Savelyeva, Taksami 1970: 187), meaning 'earth (in general)'.

**Proto-Samoyed**. **yɒə̑* (Janhunen 1977: 36–37) is retained in the meaning 'earth (soil)' in Northern Samoyed, Mator and Kamass. The root apparently also meant 'sand', q.v.

**Proto-Yukaghir**. **lewe:* (Nikolaeva 2006: 241–242) is retained in Kolyma as the word for 'earth, land'. Tundra Yukaghir has another word for 'earth': **luk-ul* (Nikolaeva 2006: 252), but preserves **lewe:* in the meaning 'land, nature'. Pace (Nikolaeva 2006: 242), for semantic reasons **lewe:* can hardly be viewed as a loan from Tungusic **lebe:(n)* 'swamp, marsh'.

23. 'to eat'.

**Proto-Yeniseian**. **si:-* (S. Starostin 1995: 274, **siɢ-*; later amended to **si:-* in Starostin 2005a). Preserved as a verbal root with its original basic meaning in Ket-Yugh and possibly in Arin and Pumpokol, but only in derived stems in Kott (where the old verbal root 'to eat' is still preserved in the nominal derivate *ši-g* 'food, meal').

**Proto-Athabaskan**. **=ha:ɲ* (or the secondary variant **=ya:ɲ*) is retained in all three branches.

**Eyak.** *=a*, cognate to Athabaskan, the Eyak paragim can point to either a PAE **CV:*-root or a **CV:n*-root.

**Proto-Athabaskan-Eyak**. **=ha:ɲ* (Athabaskan, Eyak).

**Proto-Eskimo**. **nəʁə-* (Fortescue et al. 2010: 252), retained in all branches.

**Proto-Aleut**. **qa-l-* (Bergsland 1994: 289; Golovko 1994: 206), attested in all branches. The same root as **qa-χ* 'fish / meal' q.v. Distinct from more marginal **inu-l-* 'to eat', **inu-χ* 'piece of food' (Bergsland 1994: 203).

**Proto-Chukotian**. **nu-* (Fortescue 2005: 188), retained in all languages. Cognate to the Itelmen term.

**Proto-Itelmen**. **nu-* (Volodin, Khaloimova 2001: 152; Fortescue 2005: 188). Cognate to the Chukotian term.

**Proto-Nivkh**. **ɲi-* (Fortescue 2016: 115; Savelyeva, Taksami 1965: 138; Savelyeva, Taksami 1970: 96).

**Proto-Samoyed**. **ə̆m-* (Janhunen 1977: 15), **por-* (Janhunen 1977: 127–128). Proto-Samoyed apparently had two verbs for 'to eat'. The first, **ə̆m-* (< Proto-Uralic **imi-* 'to suck'), is preserved in most daughter languages, while the second, **por-* (< Proto-Uralic **puri-* 'to gnaw, to bite') is retained only in Mator. However, derivatives from both roots – **ə̆m-sɒ* 'meat, food' and **por-sɒ* 'fish flour' can be safely reconstructed for Proto-Samoyed. This fact suggests that both **ə̆m-* and **por-* meant ‘to eat’ in Proto-Samoyed, possibly depending on the type of food.

**Proto-Yukaghir**. **leɣ-* (Nikolaeva 2006: 237–238) is retained in both Yukaghir languages.

24. ‘egg’.

**Proto-Yeniseian**. **yeʔy* (S. Starostin 1995: 232 as **yeŋ* ~ **yɔŋ*). Preserved (often in a morphologically modified form) in most Yeniseian records, with the exception of Kott. The current reconstruction is primarily based on the Ket-Yugh paradigm (sg. **ɛʔy*, pl. **e-ŋ*), but word-initial **y-* is necessary to account for the related *dʲa-nan* ‘roe’ in Kott.

**Proto-Athabaskan**. **=ʁe:ʂ* is retained in all three branches. The final retroflex points to PAE **xʷ*.

**Eyak.** *=tə=ʔuht-k*, looks like a deverbative, so may represent a new formation.

**Proto-Athabaskan-Eyak**. **χe:xʷ* (Athabaskan).

**Proto-Eskimo**. **manniɣ* (Fortescue et al. 2010: 208), retained in all branches. Distinct from **pəkyu* (Fortescue et al. 2010: 278), whose proto-meaning was apparently ‘wild eggs’.

**Proto-Aleut**. **sa:m̥la-χ* (Bergsland 1994: 351; Golovko 1994: 290), attested in all branches.

**Proto-Chukotian**. **liɣ-liɣ* (Fortescue 2005: 159), retained in all languages. Reduplicated stem. Cognate to the Itelmen term.

**Proto-Itelmen**. **lɣi-lɣi* (Volodin, Khaloimova 2001: 224; Fortescue 2005: 159). Cognate to the Chukotian term.

**Proto-Nivkh**. **ŋoyeq* (Fortescue 2016: 125; Savelyeva, Taksami 1965: 467; Savelyeva, Taksami 1970: 232). Morphologically unclear, perhaps unrelated to **ŋoy* ‘bough / penis’.

**Proto-Samoyed**. **mə̆nɒ* (Janhunen 1977: 86), retained in Enets, Nganasan and Kamass (Selkup reflex of this word means ‘penis’), goes back to Proto-Uralic **muna* ‘egg’.

**Proto-Yukaghir**. **aylʸ* (Nikolaeva 2006: 98). Modern Kolyma Yukaghir *yayčə* ‘egg’ is a Russian loanword, modern Tundra has a compound *nodod-uo* (lit. ‘bird’s child’). The original word for ‘egg’ is attested in a number of old wordlists.

25. ‘eye’.

**Proto-Yeniseian**. **de-s* (S. Starostin 1995: 220). Preserved in all daughter languages. In S. Starostin’s reconstruction, final **-s* is interpreted as a fossilized singulative suffix, a fuller variant of which may also be seen in **xu-sa* ‘one’ q.v. and several other archaic nominal stems (e. g. ‘stone’ q.v.). This argumentation is solidly supported by Arin *tie-ŋ*, which probably preserves a trace of the archaic paradigm: sg. **de-s*, pl. **de-ŋ* (the latter form shifted to **des-ŋ* in Proto-Ket-Yugh by analogy).

**Proto-Athabaskan**. **na:χ* ~ **ne:χ* is retained in all three branches. Krauss and Leer (1981: 60-62) propose a very complex solution in the light of the Degexit’an (Ingalik) form *=ma:q* ‘eye’ and the potential Tlingit *comparandum wa:q* ‘eye’. Actually, however, Degexit’an retains regular reflexes of **na:χ* in compounds and incorporation (*na:χ-* ~ *no:χ-* ~ *nʊχ-* and *na:-* ‘eye’), whereas *=ma:q* is a rare term with a specific meaning ‘animal’s eye’. Thus Degexit’an *=ma:q* and Tlingit *wa:q* can be simply unrelated to PA **na:χ* ‘eye’.

**Eyak.** *=la:χ*, cognate to Athabaskan.

**Proto-Athabaskan-Eyak**. **na:χ* (Athabaskan, Eyak).

**Proto-Eskimo**. **əðə* (Fortescue et al. 2010: 106), retained in all branches. Cognate to the Aleut term.

**Proto-Aleut**. **ða-χ* (Bergsland 1994: 158; Golovko 1994: 197), attested in all branches. Cognate to the Eskimo term.

**Proto-Chukotian**. **ləlä* (Fortescue 2005: 163), retained in all languages. Perhaps historically a reduplicated plural stem **lə-lä* as follows from the cognate Itelmen root **lo-*.

**Proto-Itelmen**. **lo-ŋ* (Volodin, Khaloimova 2001: 145; Fortescue 2005: 163). Final **-ŋ* is the singulative exponent. Reduplicated plural: **lu-l-*. Cognate to the Chukotian term.

**Proto-Nivkh**. **ɲŋaɣ* (Fortescue 2016: 116; Savelyeva, Taksami 1965: 116; Savelyeva, Taksami 1970: 221). Morphologically unclear.

**Proto-Samoyed**. **sə̆ymä* (Janhunen 1977: 132), retained in all daughter languages, goes back to Proto-Uralic **silmä* ‘eye’.

**Proto-Yukaghir**. Kolyma **waŋ-čʸə* (Nikolaeva 2006: 452) vs. Tundra **yöː-ntə* (Nikolaeva 2006: 191). Kolyma word for ‘eye’ is related to the verb ‘to look for, to seek’; Tundra word for ‘eye’ is derived from **yöː-* ‘to see’.

26. ‘fat’.

**Proto-Yeniseian**. **gɨʔd* (S. Starostin 1995: 228). Preserved in all daughter languages where attested, but not found in Arin and Pumpokol.

**Proto-Athabaskan**. **q’aχ* is retained in all three branches.

**Eyak.** *=q’əχ*, cognate to Athabaskan.

**Proto-Athabaskan-Eyak**. **q’aχ* (Athabaskan, Eyak).

**Proto-Eskimo**. **uqðu-ʁ* (Fortescue et al. 2010: 412), retained in all branches. Its narrow meaning can be ‘blubber, seal oil’.

**Proto-Aleut**. **čaðu-χ* (Bergsland 1994: 124; Golovko 1994: 142, 208), attested in all branches.

**Proto-Chukotian**. **äčʁə-n* (Fortescue 2005: 25), retained in all languages except for Kerek.

**Proto-Itelmen**. **kʷəč’x* (Volodin, Khaloimova 2001: 153; Fortescue 2005: 363).

**Proto-Nivkh**. **tom* (Fortescue 2016: 150; Savelyeva, Taksami 1965: 141; Savelyeva, Taksami 1970: 382).

**Proto-Samoyed**. **yür* (Janhunen 1977: 50), **tuyt ~ *cu̯yt ~ *tuycɩ~ *cu̯ycɩ*(Janhunen 1977: 166). The semantic difference between the two Proto-Samoyed words for ‘fat’ is not clear. The first one, **yür*, is a Turkic loanword, the second one probably is the original Samoyed word for ‘fat’.

**Proto-Yukaghir**. Kolyma **poŋičə* (Nikolaeva 2006: 360) and **suwo-npə* (Nikolaeva 2006: 421) vs. Tundra **nʸan-* (Nikolaeva 2006: 288), related to Kolyma words ‘fish oil’ and ‘to overeat’. The semantic difference between the two Kolyma words for ‘fat’ is not clear from the available lexicographic sources, so we are forced to list three roots here.

27. ‘feather’.

**Proto-Yeniseian**. **ʔaːsi* (S. Starostin 1995: 205). Preserved in all daughter languages (but not attested in Pumpokol). Vocalic correspondences are unclear, but the data suggest that, most likely, the stem-final **-i* has influenced the root vocalism in the Kott-Arin branch (**ʔaːsi* > **ʔiːsi*).

**Proto-Athabaskan**. **t’aː* is retained in all three branches.

**Eyak.** *t’ahl* ‘feather / leaf’, apparently the original meaning was ‘leaf’ (q.v.) with the later shift > ‘feather’ according to the areal isogloss.

**Proto-Athabaskan-Eyak**. **t’aː* (Athabaskan).

**Proto-Eskimo**. **culu-ɣ* (Fortescue et al. 2010: 100), retained in all branches. In Inuit, it shifted into the specific meaning ‘large feather, wing feather’, being superseded with **məl-quʁ* ‘body hair, fur’ (Fortescue et al. 2010: 216), which acquires the polysemy ‘feather (in general); body hair, fur’ in Inuit.

**Proto-Aleut**. **haka-χ* (Bergsland 1994: 42; Golovko 1994: 243), attested in all branches.

**Proto-Chukotian**. **teŋə-lŋən* (Fortescue 2005: 284), retained in all languages, meaning ‘wing feather’. Distinct from **ɣəð-ɣəð* ‘fur’ (Fortescue 2005: 64) which acquires the polysemy ‘fur / down, feather (in general)’ in Koryak-Alutor.

**Proto-Itelmen**. **sʸisʸi* (Fortescue 2005: 60), in some sources documented with polysemy ‘wing / feather’. In modern Western, shifted into the meaning ‘wing’ (Volodin, Khaloimova 2001: 83), having been superseded with **čeɬx* ‘fur / down’ which is currently glossed as ‘fur / feather / down’ (Volodin, Khaloimova 2001: 104, 186; Fortescue 2005: 64).

**Proto-Nivkh**. **dup-r* (Fortescue 2016: 47; Savelyeva, Taksami 1965: 291; Savelyeva, Taksami 1970: 365). See Panfilov 1962: 59; Savelyeva, Taksami 1970: 530– 531 for the suffix **-r*.

**Proto-Samoyed**. **tuǝ̆* (Janhunen 1977: 166), retained in Northern Samoyed, Mator and Selkup, is related to Finno-Ugric **tulka* 'feather / wing'.

**Proto-Yukaghir**. **tiw-il* (Nikolaeva 2006: 431–432) is attested in Kolyma Yukaghir and a number of old worlists. Words for 'feather' are absent from Kurilov 2001, the main lexicographic source for Tundra Yukaghir.

**Proto-Burushaski**. *$p^h$olʁo* was borrowed as a marginal term for 'feather' in some Shina dialects (note that the basic Shina word for this meaning has Indo-Aryan etymology, Anton Kogan, p.c.).

28. 'fire'.

**Proto-Yeniseian**. **boʔk* (S. Starostin 1995: 212). Preserved in Ket-Yugh and Pumpokol. In Kott and Arin (probably, in Proto-Kott-Arin), replaced by a nominalization of Proto-Yeniseian **qɔʔt* 'to burn' q.v.

**Proto-Athabaskan**. *$q^{hw}$enʔ* is retained in all three branches.

**Eyak.** *tǝ=q'aː-k*, derived from *=q'a* 'to burn', so should represent a new formation.

**Proto-Athabaskan-Eyak**. *$q^{hw}$en* (Athabaskan).

**Proto-Eskimo**. **ǝk-nǝʁ* (Fortescue et al. 2010: 111), retained in all branches. Looks like a deverbative from **ǝkǝ-* 'to burn' with the common nominalizer *-*nǝʁ*. On the other hand, irregular dropping of final *-ʁ* is some lects can point to an independent stem **ǝknǝ-ʁ* which was latter reanalyzed as a suffixed deverbative (thus Oleg Mudrak, p.c.).

**Proto-Aleut**. **qiɣna-χ* (Bergsland 1994: 320; Golovko 1994: 235), attested in all branches.

**Proto-Chukotian**. **milɣǝ-* (Fortescue 2005: 176), retained as the basic term in Chukchi (one of the non-east dialects, Moll, Inenlikei 1957: 77), Koryak (Zhukova 1990: 57, 163), Alutor (Kibrik et al. 2004: 455). Cognate to the Itelmen term.

**Proto-Itelmen**. **xi=mɬx* (Volodin, Khaloimova 2001: 180; Fortescue 2005: 176). Partial reduplication; cognate to the Chukotian term. In some lects, superseded with **piŋ-č* (Fortescue 2005: 364).

**Proto-Nivkh**. **tu-ɣr* ~ **tuɣ-r* (Fortescue 2016: 152; Savelyeva, Taksami 1965: 255; Savelyeva, Taksami 1970: 384). Either derived from **tuv-* 'to burn (tr.)' q.v. with the rare suffix *-*ɣr* (for which see Fortescue 2016: 175; Panfilov 1962: 62; Savelyeva, Taksami 1970: 523), although the loss of *-v-* seems irregular. Or derived from a root **tuɣ-* '?' with the rare suffix *-*r* (for which see Panfilov 1962: 59; Savelyeva, Taksami 1970: 530– 531). Phonetic similarity with Proto-Tungusic **toɣa* 'fire' (> Orok *tawa*, Evenki *toɣo*, Nanai *tao*, all 'fire') seems accidental, since the scenario Tungusic > Nivkh does not explain the Nivkh non-productive suffixation.

**Proto-Samoyed**. **tuy* (Janhunen 1977: 166), retained in all Samoyed languages, goes back to Proto-Uralic **tuli* 'fire'.

**Proto-Yukaghir**. **yeŋkilǝ* 'fire' (Nikolaeva 2006: 188) is attested in Jochelson's Kolyma materials, as well as in a number of old wordlists. Although modern Kolyma and Tundra Yukaghir have reflexes of **loč-il* 'fire / firewood' (Nikolaeva 2006: 245–246) as the main word for fire, **yeŋkilǝ* 'fire' is attested both in Old Kolyma wordlists and in Omok. According to Jochelson, in Kolyma Yukaghir "[l̥]o′čil is a modern term which also means fuel and, more generally, any material for a fire or the hearth. Yegi′l̥e is an ancient word" (Jochelson 1926: 141). We tentatively reconstruct an opposition **yeŋkilǝ* 'fire' vs. **loč-il* 'firewood' in Proto-Yukaghir with a homoplastic development 'firewood' > 'fire / firewood' in both modern languages.

29. 'fish'.

**Proto-Yeniseian**. **ciːk* (S. Starostin 1995: 214). Preserved in the original meaning in Kott-Arin and in Pumpokol. In Ket-Yugh, replaced in the meaning 'fish' with **ʔiˑs* 'meat' and only preserved in the meaning 'snake'. The shift chain 'meat' > 'fish / meat', 'fish' > 'snake' is, overall, the most economic solution, given the distribution of cognates in daughter languages.

**Proto-Athabaskan**. **łuːq'* is retained in all three branches.
**Eyak.** *tʰəʔ-yaʔ*, literally 'the one in water', a transparent new formation.
**Proto-Athabaskan-Eyak**. **łuq'* (Athabaskan).
**Proto-Eskimo**. **iqałuɣ* (Fortescue et al. 2010: 154), retained in all branches. Looks like an old suffixed derivation, cf. **-łuɣ* 'bad' (Fortescue et al. 2010: 453).
**Proto-Aleut**. **qa-χ* (Bergsland 1994: 289; Golovko 1994: 262), attested in all branches. Polysemy: 'fish / meal'. The same root as **qa-l-* 'to eat' q.v.
**Proto-Chukotian**. **ənnə* (Fortescue 2005: 345), retained in all languages. Cognate to the Itelmen term.
**Proto-Itelmen**. **ənč* (Volodin, Khaloimova 2001: 201; Fortescue 2005: 345). Cognate to the Chukotian term.
**Proto-Nivkh**. **t̥o* (Fortescue 2016: 34; Savelyeva, Taksami 1965: 375; Savelyeva, Taksami 1970: 450).
**Proto-Samoyed**. **kɒlä* (Janhunen 1977: 59), retained in all Samoyed languages, goes back to Proto-Uralic **kala* 'fish'.
**Proto-Yukaghir**. Kolyma **an-il* ~ **wan-il* (Nikolaeva 2006: 105) vs. Tundra **olʸoɣə* (Nikolaeva 2006: 325).
**Proto-Burushaski**. **ɕʰumo* was borrowed in Shina (not *vice versa* since Shina *čʰumu* 'fish' lacks Indo-Aryan etymology).

30. 'to fly (sg.)'.
**Proto-Yeniseian**. **=doːq* (S. Starostin 1995: 223). Preserved in Ket-Yugh, but not in Kott; not attested in either Arin or Pumpokol. The root **=doːq* is still attested in Kott in the meaning 'to jump'.
**Proto-Athabaskan**. **=t=ʔaq* is retained in all three branches.
**Eyak.** *=tə=l=k'aʔt'*.
**Proto-Athabaskan-Eyak**. **=ʔaq* (Athabaskan), **=k'Vt'* (Eyak).
**Proto-Eskimo**. **təŋə-* (Fortescue et al. 2010: 372), retained in all branches. In Inuit (or already in Proto-Inuit?) tends to shift into the meaning 'to fly up, fly away', having been superseded with the durative stem **təŋ-miɣ-* 'to be flying'.
**Proto-Aleut**. **iɣa-χta-l-* (Bergsland 1994: 175; Golovko 1994: 222), attested in all branches. A durative from **iɣa-l-* 'to start to fly, fly up'.
**Proto-Chukotian**. **ðiŋä-* (Fortescue 2005: 59), retained as generic 'to fly' in Koryak, having shifted into such specific meanings as 'to fly up' or 'to fly off' in other languages (where the durative semantics 'to fly' is now expressed by various suffixed stems derived from **ðiŋä-*). Cognate to the Itelmen term.
**Proto-Itelmen**. **sʸiŋ-* (Volodin, Khaloimova 2001: 167; Fortescue 2005: 59). Cognate to the Chukotian term.
**Proto-Nivkh**. **bəy-* (Fortescue 2016: 28; Savelyeva, Taksami 1965: 198; Savelyeva, Taksami 1970: 277).
**Proto-Samoyed**. **tey-* (Janhunen 1977: 161–162), retained in all languages save Kamass, goes back to Proto-Uralic **ʂelki-* 'to fly'.
**Proto-Yukaghir**. Kolyma **mere-* (Nikolaeva 2006: 266) vs. Tundra **čen-* (Nikolaeva 2006: 129).

31. 'foot'.
**Proto-Yeniseian**. **bul* (S. Starostin 1995: 213). Preserved everywhere except for Pumpokol, where **bul* 'foot' *may* have been replaced with *an-* = Arin *an* 'thigh', Kott *aːn-ar* 'thigh' (there is a probability that Pumpokol *an-iŋ* really means 'legs', whereas the proper word for 'feet' was not recorded).
**Proto-Athabaskan**. **qʰe* is retained in all three branches.
**Eyak.** *=k'ʷahš*.

**Proto-Athabaskan-Eyak**. **qʰe* (Athabaskan), **k'ʷVnš* (Eyak; we follow Leer 2008a and reconstruct a medial nasal on account of vowel aspiration in Eyak).

**Proto-Eskimo**. **itəɣ-aʁ* (Fortescue et al. 2010: 160), retained in all branches. Literally 'toe cap-like' from **itəɣ* 'toe cap, tip of boot'.

**Proto-Aleut**. **kita-χ* (Bergsland 1994: 241; Golovko 1994: 233), attested in all branches.

**Proto-Chukotian**. **kətka* (Fortescue 2005: 154), retained in all languages with polysemy 'foot / leg'. Cognate to the Itelmen term.

**Proto-Itelmen**. **ktxa-* (Volodin, Khaloimova 2001: 178; Fortescue 2005: 154), polysemy 'foot / leg'. Cognate to the Chukotian term.

**Proto-Nivkh**. **ŋa=ṭɣ* (Fortescue 2016: 117; Savelyeva, Taksami 1965: 243; Savelyeva, Taksami 1970: 238). Polysemy: 'foot / leg'. A prefixal element **ŋa-*, common for body part names, can be singled out. Distinct from a more specific and rarely used term **ŋa=zl* 'sole of foot' (Fortescue 2016: 124; Savelyeva, Taksami 1970: 234).

**Proto-Samoyed**. **ɒy* (Janhunen 1977: 17) is retained in all daughter languages save Selkup.

**Proto-Yukaghir**. Kolyma **noy-l* (Nikolaeva 2006: 306) vs. Tundra **öɣur-* (Nikolaeva 2006: 320), related to Kolyma word for 'ski with fur on the bottom'.

32. 'full'.

**Proto-Yeniseian**. **ʔute* (S. Starostin 1995: 201). Preserved in all daughter languages where attested, but not found in Arin or Pumpokol.

**Proto-Athabaskan**. **=wen* is retained in all three branches.

**Eyak.** *təq-i-taʔ ...-tə-ʔya*, a transparent new formation based on the preverb *təq* 'upstream, upriver' and the postposition *-taʔ* 'arrival at'.

**Proto-Athabaskan-Eyak**. **=wen* (Athabaskan).

**Proto-Eskimo**. An unstable item. The best candidate seems to be **cilə-* (Fortescue et al. 2010: 86), which is attested with the meaning '(to be) full' in both Yupik (Central Siberian Yupik, Menovshchikov 1988: 127, 214) and Inuit (North Alaskan Inuit, MacLean 2014: 307). In other Inuit, it shifts into the neighboring meanings 'to have a full stomach, suffer from indigestion' or 'to leave the rest (having had enough)'.

**Proto-Aleut**. **čχa-l-* (Bergsland 1994: 134; Golovko 1994: 249), attested in all branches.

**Proto-Chukotian**. **yərʁ-* (Fortescue 2005: 123) means 'to fill (up)'. At least the Chukchi (Inenlikei 2005) and Alutor (Kibrik et al. 2004: 414) basic expressions for 'full' are derived from this verb.

**Proto-Itelmen**. **txnu- ~ *t=xnu-* (Volodin, Khaloimova 2001: 35, 89, 190) 'to fill (trans.)' from which the expression for 'full' is derived. Initial **t* can be an old causative exponent.

**Proto-Nivkh**. **ṭar-* (Fortescue 2016: 30; Savelyeva, Taksami 1965: 313; Savelyeva, Taksami 1970: 443).

**Proto-Samoyed**. **tärə̑* (Janhunen 1977: 158), retained in Selkup as the word for 'full' (Northern Samoyed cognates mean 'interior'), goes back to Proto-Uralic **täwði* 'full' (Aikio 2002: 31–34).

**Proto-Yukaghir**. **poto-* (Nikolaeva 2006: 363) is attested in Tundra Yukaghir. Kolyma *qodo-nʸe-y* 'full' is derived from *qodo* 'contents, handful' and further related to the verb *qodo:-* 'to lie down' (Nikolaeva 2006: 220–221).

33. 'to give'.

**Proto-Yeniseian**. There are altogether three different roots / stems attested with the meaning 'to give' in various Yeniseian languages. Of these: (1) Kott-Arin **=pen-* (always functions as the second root in a composite stem, with differing first elements) is compared by S. Starostin (2005) with Yugh *=fin* in the composite verbal stem *χɜdʸiŋ=fin ~ χɜdʸiŋ=fan* 'to give back; to give away'; however, external Ket evidence shows that it is Yugh *χɜdʸiŋ-*, not *=fin*, that carries the main lexical meaning of 'give back, give away' (= Ket *qɜr-am ~ qɜr-aŋ* id.). Considering that in Kott and Arin, *=pen-* is also not found on its own, it is more likely that the verb was a general "directional" auxiliary in Proto-Yeniseian rather than an original ‘to give’. (2) Ket-Yugh *=aq*, likewise, is a verbal root with much broader semantics than ‘to give’. (3) Consequently, the only verbal stem that is attested *exclusively* in the meaning ‘to give’ is Ket-Yugh **n=...=o*. Furthermore, its highly unusual shape (monovocalic root + very rare directional prefix) is an additional indirect hint at archaicity.

**Proto-Athabaskan**. **=ʔaː*, a classificatory verb ‘to handle smth. / to be in position (of smth.)’, retained in all three branches.

**Eyak.** *=tʰa*, a generic classificatory verb ‘to handle smth. / to be in position (of smth.)’; since the second classificatory verb which is also widely used, but nevertheless not so generally is *=ʔa* (cognate to Athabaskan **=ʔaː*), it is likely that **=ʔa* was the Proto-Athabaskan-Eyak verb used in the neutral expression ‘to handle smth. / to be in position (of smth.)’ (particularly ‘to give’), gradually superseded with *=tʰa* in Eyak.

**Proto-Athabaskan-Eyak**. **=ʔaː* (Athabaskan).

**Proto-Eskimo**. **tunə-* (Fortescue et al. 2010: 381), retained in all branches.

**Proto-Aleut**. **aχ-s- ~ *uχ-s-* (Bergsland 1994: 31; Golovko 1994: 37, 200), attested in all branches. Polysemy: ‘to put, place / to give’; can be an old derivative from **a-/u-* ‘to be’.

**Proto-Chukotian**. **yəl-* (Fortescue 2005: 119), retained in all languages. Cognate to the Itelmen term.

**Proto-Itelmen**. **zʸil-* (Volodin, Khaloimova 2001: 147; Fortescue 2005: 119). Cognate to the Chukotian term.

**Proto-Nivkh**. **kim-* (Fortescue 2016: 86; Savelyeva, Taksami 1965: 124; Savelyeva, Taksami 1970: 92).

**Proto-Samoyed**. **mi-* (Janhunen 1977: 94), **tɒ-* (Janhunen 1977: 94). Judging by the situation in Tundra Nenets, the two verbs for ‘give’ were used depending on the person of recipient: **mi-* (< Proto-Uralic **meɣi-* ‘to give’) was used with 2nd and 3rd person recipient, while **tɒ-* was required in sentences with 1st person recipient.

**Proto-Yukaghir**. **tant-* ‘give (to a speech act participant)’ (Nikolaeva 2006: 426), **key-* ‘give (to a 3rd person)’ (Nikolaeva 2006: 203). Both verbs are retained in both Yukaghir languages.

34. ‘good’.

**Proto-Yeniseian**. **haq-* (S. Starostin 1995: 230). Preserved in all daughter languages (but not attested in Pumpokol).

**Proto-Athabaskan**. **=žʉː ~ *=žuː* is retained in all three branches. See Krauss 1977: 33 for the discrepancy in reflexes between **žˌ* and **ž*. Since the fricative retroflex **šˌ* (**žˌ*) tends to merge with the ordinary shibilant **š* (**ž*), the original shape of the PA form should be **=žʉː*. The PA retroflex hussing points to PAE **xʷ*.

**Eyak.** *=cuː*, goes back to PAET **kʸuː*, apparently unrelated to Proto-Athabaskan **=žʉː* ‘good’ (Leer 2010: 177).

**Proto-Athabaskan-Eyak**. **=xʷuː* (Athabaskan), **=cuː* (Eyak).

**Proto-Eskimo**. An unstable and not very reliable documented item which cannot be reconstructed with certainty.

**Proto-Aleut**. **iʁama-na-l-* (Bergsland 1994: 184), attested in Eastern and Atkan. In modern Atkan, shifted to the narrower meaning ‘good (of person)’ (Golovko 1994: 48), whereas the meaning ‘good (of object)’ is now expressed with *suɣða-na-l-* ‘good, beautiful, nice, pretty’ (Golovko 1994: 113, 284), whose original meaning should be ‘notable, distinguished; beautiful, fancy’ (Bergsland 1994: 375).

**Proto-Chukotian**. **mäl-* (Fortescue 2005: 171), a basic root for ‘good’ at least in Chukchi, Alutor and perhaps in Kerek. Cognate to the Itelmen term. Distinct from **täŋ-* (Fortescue 2005: 281) whose meaning is rather to be reconstructed as ‘good, kind, nice’.

**Proto-Itelmen**. **mel-* (Volodin, Khaloimova 2001: 58; Fortescue 2005: 172). Cognate to the Chukotian term.

**Proto-Nivkh**. **ur-* (Fortescue 2016: 157; Savelyeva, Taksami 1965: 448; Savelyeva, Taksami 1970: 395).
**Proto-Samoyed**. **sǝ̆mɒ* (Janhunen 1977: 132–133) is retained in Northern Samoyed and Selkup.
**Proto-Yukaghir**. **omo-* (Nikolaeva 2006: 327–328) is retained in both Yukaghir languages.
**Proto-Burushaski**. *ʃua* is likely borrowed from Dardic (cf. Shina *sʸoː*, Palula *šuːo* 'good').

35. 'green'.
**Proto-Yeniseian**. Not reconstructible. The item is poorly attested in all extinct languages; not a single match between two different languages can be detected; and there are reasons to assume that the meaning 'green' was not lexically distinct even in Proto-Ket-Yugh.
**Proto-Athabaskan**. **=cʰuχ* 'green / yellow' is retained in Pacific Coast and Northern.
**Eyak.** *tiːyaʔ-kaʔ*, literally 'salt water-like', a transparent new formation.
**Proto-Athabaskan-Eyak**. **=cʰuχ* (Athabaskan), although the Athabaskan color term may also be secondary, if it is cognate to Eyak *=cʰeʔq'* 'urine'.
**Proto-Eskimo**. **cuŋa-* (Fortescue et al. 2010: 101), a substantive 'gall, bile' from which expressions for '(to be) blue/green' are derived in the majority of lects.
**Proto-Aleut**. **čið-ʁi-l-* (Bergsland 1994: 135; Golovko 1994: 212), attested in all branches, meaning 'green/blue'.
**Proto-Chukotian**. **wǝt-* (Fortescue 2005: 337), retained as a basic root for 'green' in Chukchi and Koryak (Alutor 'green' < 'gall'), cognate to the Itelmen term. The Proto-Chukotian word for 'leaf' (q.v.) is based on this root. Comparison with Itelmen suggests that the color semantics might be primary for Proto-Chukotian, whereas the botanic meaning is secondary.
**Proto-Itelmen**. **ʌɬ-* (Volodin, Khaloimova 2001: 157; Fortescue 2005: 337), meaning 'green/blue' or even 'green/blue/yellow', although the meaning 'yellow' is poorly documented. Cognate to the Chukotian term.
**Proto-Nivkh**. **tǝɣ- ~ *dǝɣ-* (Fortescue 2016: 154). Polysemy 'green / blue'. In Amur, retained with the meaning 'blue' (Savelyeva, Taksami 1965: 387; Savelyeva, Taksami 1970: 367), whereas 'green' is expressed with the new formation *ɲlays-vala-* (Savelyeva, Taksami 1965: 161; Savelyeva, Taksami 1970: 217), literary 'greenery-colored'.
**Proto-Samoyed**. **tǝ̆ŋkV- ~ *cǝ̆ŋkV-* 'green / blue' (Helimski 1997: 356) is retained in Forest Nenets and Mator.
**Proto-Yukaghir**. **qomo-* (Nikolaeva 2006: 385) is attested in Tundra Yukaghir. Kolyma *nʸoŋoː-* 'green, blue' is a Tungusic loanword (Nikolaeva 2006: 308).

36. 'hair'.
**Proto-Yeniseian**. **cǝŋe* (S. Starostin 1995: 213). Preserved in all daughter languages.
**Proto-Athabaskan**. **=ʁa-ʔ* is retained in all three branches, meaning generic 'hair, fur'.
**Eyak.** *leːɬ*, meaning specifically 'head hair'.
**Proto-Athabaskan-Eyak**. **=χVw* (Athabaskan). Tlingit *=χaːw* 'hair (generic)' and Eyak *=χuʔ* 'fur, body-hair' prove that **=χVʔʷ* should be a proto-term for '(head) hair', whereas Eyak *leːɬ* 'head hair' is an innovation.
**Proto-Eskimo**. **nuya-ʁ* (Fortescue et al. 2010: 267), retained in all branches, meaning 'head hair'. Distinct from **mǝl-quʁ* 'body hair, fur' (Fortescue et al. 2010: 216).
**Proto-Aleut**. **imli-χ* (Bergsland 1994: 198; Golovko 1994: 192), attested in all branches, meaning 'head hair'. Distinct from **čŋa-χ* 'body hair, fur' (Bergsland 1994: 147).
**Proto-Chukotian**. Compound **kǝð=wir* (Fortescue 2005: 143) is retained in all languages with the meaning 'hair (generic)'. The first morpheme **kǝð* may denote something related to head, it is also attested in such compounds as **kǝð-ðel* 'forehead' (Fortescue 2005: 143), **kǝrǝ-tkǝn ~ *kǝðǝ-tkǝn* 'top' (Fortescue 2005: 152), **kǝð-täl* 'braid, plait' (Fortescue 2005: 143). If so, the second element should be the main meaningful root here.
**Proto-Itelmen**. **k'ʷimi* (Volodin, Khaloimova 2001: 141; Fortescue 2005: 367), meaning 'hair (generic)'.

**Proto-Nivkh**. **ŋa=mɣ* (Fortescue 2016: 120; Savelyeva, Taksami 1965: 91; Savelyeva, Taksami 1970: 236). Attested at least in Amur (*ŋəŋg*) and East Sakhalin (*ŋamx*), meaning 'hair (generic, normally applicable to human)'. A prefixal element **ŋa-*, common for body part names, can be singled out. Tends either to be superseded with or to contaminate with **ŋa=vrki* 'fur (of animal)' (Fortescue 2016: 120; Savelyeva, Taksami 1970: 233).

**Proto-Samoyed**. **ɜptə̆* (Janhunen 1977: 21), retained in all Samoyed languages, goes back to Proto-Uralic **ɨpti* 'hair'.

**Proto-Yukaghir**. **monilə* (Nikolaeva 2006: 258) is retained in both Yukaghir languages.

37. 'hand'.

**Proto-Yeniseian**. The meaning 'hand' is notoriously unstable in Yeniseian languages: almost every language has its own etymological equivalent (sometimes two!), and most of the etymological connections are problematic. The best chances lie with the pairing of Ket *hɜŋn* and Arin *p*$^{yh}$*yaga* (= *pega*), which allows S. Starostin (1995: 254) to reconstruct the protoform as **pVg-*. The semantic matching is exact, and the correspondences are generally reconcilable. However, there is some doubt as to whether the Ket word is indeed the primary equivalent for 'hand', and, subsequently, this would influence Proto-Yeniseian semantics. One should probably also make a note of Ket *l*$^{y}$*aʔŋ* 'hand', with a parallel in Arin: *lan-t*$^{y}$*uːŋ ~ l*$^{y}$*an-puy* 'wing' (Starostin 2005a); the semantic shift 'hand' > 'wing' is theoretically possible.

**Proto-Athabaskan**. **=laː* is retained in all three branches.

**Eyak.** *=yə=q'aʔc'* 'hand'; however internal Eyak evidence such as *č'ãːʔ* '5' points out that *=č'el-ih ~ č'ãː-* 'arm' previously denoted 'hand / arm'.

**Proto-Athabaskan-Eyak**. **łaː* (Athabaskan), **č'Vn* (Eyak).

**Proto-Eskimo**. **aðɣa-* (Fortescue et al. 2010: 4), retained in all branches.

**Proto-Aleut**. **ča-χ* (Bergsland 1994: 123; Golovko 1994: 262), attested in all branches.

**Proto-Chukotian**. Browsing through available sources suggests that **mənɣə* (Fortescue 2005: 184) is the basic term with polysemy 'hand / arm' in all languages. The Proto-Chukotian expression for '10' (Fortescue 2005: 184) is based on this root. Distinct from the specific term **käɣə* (Fortescue 2005: 129), meaning 'palm (of hand)'. The latter is probably cognate to the Itelmen word for 'hand'.

**Proto-Itelmen**. **xk'i-* (Volodin, Khaloimova 2001: 98, 201; Ono 2003: 106; Fortescue 2005: 129), meaning 'hand / arm'. Distinct from **manzə-* 'palm (of hand)' (Volodin, Khaloimova 2001: 56; Ono 2003: 106).

**Proto-Nivkh**. **damk* (Fortescue 2016: 39; Savelyeva, Taksami 1965: 374; Savelyeva, Taksami 1970: 368), glossed as 'hand' by Fortescue, but it has the generic meaning 'hand / arm' at least in Amur and North Sakhalin. Distinct from **dot* (Fortescue 2016: 45; Savelyeva, Taksami 1965: 374; Savelyeva, Taksami 1970: 360), which is glossed as 'arm' by Fortescue, but at least in Amur it means specifically 'forearm'.

**Proto-Samoyed**. **utɒ* (Janhunen 1977: 30) is retained in all Samoyed languages. Judging by its phonology, the word cannot be inherited from Proto-Uralic.

**Proto-Yukaghir**. **tolon-* (Nikolaeva 2006: 434), is attested in modern Tundra Yukaghir as well as in two old Kolyma wordlists. The original opposition **tolon-* 'hand' vs. **n*$^{y}$*uŋkən* 'arm' (Nikolaeva 2006: 315) is preserved in Tundra, whereas modern Kolyma *nugen* means both 'hand' and 'arm'.

38. 'head'.

**Proto-Yeniseian**. **ciʔɢe* (S. Starostin 1995: 214). Preserved in Ket-Yugh (although mostly replaced in modern Ket), Kott, and possibly Arin (or at least one of the Arin dialects).

**Proto-Athabaskan**. **=c*$^{h}$*i-ʔ* is retained in all three branches.

**Eyak.** *=lə=q*$^{h}$*ah ~ =šaːw*, synchronously there are two common expressions for 'head' in Eyak (*=šaːw* can be a Tlingit loan, but it is not certain). It seems, however, that the old root for 'head' is retained as *=c*$^{h}$*ĩʔ* 'neck' (q.v.), because, firstly, *c*$^{h}$*ĩʔ* as a qualifier prefix or in some fossilized constructions means 'head', secondly, it corresponds to Athabaskan *=cʰi-ʔ 'head', if one assumes a *n*-extension in Proto-Eyak: *cʰĩʔ* < **cʰi-n*. As noted in Leer 2010: 179, the same *n*-extension is observed for this root in the compound stems in Carrier (Athabaskan) and Tlingit (if Tlingit *šá* 'head' is related).

**Proto-Athabaskan-Eyak**. **cʰi* ~ **cʰi-n* (Athabaskan). Leer 2010: 179 compares Tlingit *šá* 'head' and reconstructs PEAT *kʸʰeŋʔ*. But if the Tlingit form is indeed related, PAE $c^h$ / Tlingit *š* is a very specific row of correspondences, distinct from the more reliable row PAE $c^h$ / Tlingit $k^h$ (presumably < PAET **kʸʰ*), and it remains unclear why we should merge these two row in a single proto-phoneme (as *per* Leer) and, second, why the consistent sibilant reflexes should point to a guttural proto-phoneme.

**Proto-Eskimo**. **nayə-quʁ* (Fortescue et al. 2010: 243), retained in Yupik and Inuit. The suffix *-*quʁ* means 'smth. associated with or attached to smth.' (Fortescue et al. 2010: 467).

**Proto-Aleut**. **kamɣi-χ* (Bergsland 1994: 226; Golovko 1994: 198), attested in all branches.

**Proto-Chukotian**. **läwət* (Fortescue 2005: 158), retained in all languages.

**Proto-Itelmen**. **ktxi-* (Volodin, Khaloimova 2001: 145; Fortescue 2005: 152), tends to be superseded (see Ono 2003: 104; Fortescue 2005: 368) with derivatives from **k'ʷimi* 'hair' q.v.

**Proto-Nivkh**. **d̥oŋ-kr* (Fortescue 2016: 53; Savelyeva, Taksami 1965: 117; Savelyeva, Taksami 1970: 351). Cf. **d̥oŋa-* 'to turn away from' (Savelyeva, Taksami 1970: 351).

**Proto-Samoyed**. **ɒywɒ* (Janhunen 1977: 17), retained in Northern Samoyed and Mator, goes back to Proto-Uralic **oywa* 'head'.

**Proto-Yukaghir**. **yo:* (Nikolaeva 2006: 190) is retained in both Yukaghir languages.

39. 'to hear'.

**Proto-Yeniseian**. *=*ta* (S. Starostin 1995: 291). Preserved in all daughter languages (not attested in Pumpokol, but the root may be present in the form *hiti-fun* 'to be silent', literally 'without hearing').

**Proto-Athabaskan**. The main candidates are *=*c'a:ɲ* and *=*c'aq* ~ *=*c'aχ*. Apparently *=*c'aq* (and further *=*c'aχ*) originates from the aspectual suffixed stem *=*c'a:ɲ-q* with the regular loss of nasal (Leer 2008b).

**Eyak.** *=tə=ɬə=č'a:q'*, it is unclear whether the Eyak root is related to Athabaskan *=*c'a:ɲ* and *=*c'a:nʸ-q* or not.

**Proto-Athabaskan-Eyak**. *=*c'a:ɲ* (Athabaskan), *=*č'a:q'* (Eyak).

**Proto-Eskimo**. **tuca-ʁ* (Fortescue et al. 2010: 376), it means 'to hear' in Inuit, having shifted into more specific meanings such as 'to learn, hear from, find out' or 'to hear and comprehend' in Yupik; cognate to the Aleut term. Distinct from **naɣa-t-* 'to listen' (Fortescue et al. 2010: 226), which acquired the meaning 'to hear' in some (not all) Yupik lects. Finally it is not excluded that a genuine root for either 'to hear' or 'to listen' is retained in **ciɣ-un* 'ear' (q.v.) which contains the instrumental suffix *-*un*, cf. the same pattern in Aleut.

**Proto-Aleut**. **tut-s-* (Bergsland 1994: 410; Golovko 1994: 127, 268), meaning 'to hear'; in modern lects superseded with the suffixed stem *tut-a-l-* 'to hear / to listen', attested in all branches. Cognate to the Eskimo term.

**Proto-Chukotian**. **valom-* (Fortescue 2005: 313), retained in all languages, morphologically unclear; can be related to Itelmen **ilwsʸ-* 'to hear'. Distinct from **palom-tel-* 'to listen' (Fortescue 2005: 208), which partially contaminated with **valom-* perhaps already in Proto-Chukotian as well in some attested languages such as Alutor. Additionally one can suspect contamination with **vilu* 'ear' q.v.

**Proto-Itelmen**. **ilwsʸ-* (Volodin, Khaloimova 2001: 119; Fortescue 2005: 369), meaning 'to hear / to listen'. Resembles **elwe-* 'ear' q.v. Can be related to Chukotian **valom-* 'to hear'.

**Proto-Nivkh**. **mə-* (Fortescue 2016: 109; Savelyeva, Taksami 1965: 393; Savelyeva, Taksami 1970: 201), meaning 'to hear / to listen'.

**Proto-Samoyed**. **yüntə̆-* (Janhunen 1977: 49) is retained in most Samoyed languages.

**Proto-Yukaghir**. **með-* (Nikolaeva 2006: 261). Although formally reconstructible to Proto-Yukaghir level, this word is apparently an early borrowing from Tungusic.

40. 'heart'.

**Proto-Yeniseian**. **pu* (S. Starostin 1995: 251). Preserved in Ket-Yugh and Pumpokol. It is not a certified fact, despite the confidence in S. Starostin 1995: 251, that Ket-Yugh / Pumpokol **pu* 'heart' is etymologically connected with **pɨy* 'belly' q.v., despite phonetic similarity and semantic proximity. For the time being, it is preferable to judge it as an individual root with a precise Swadesh meaning ('heart') and not a member of any Proto-Yeniseian "word-family".

**Proto-Athabaskan**. **čę̣ːy* is retained in all three branches. The initial retroflex points to PAE **kʷ*.

**Eyak.** *=ʔuq-ɬ*, final *-ɬ* is a common nominal suffix (originally instrumental).

**Proto-Athabaskan-Eyak**. **kʷeːy* (Athabaskan), **ʔuq* (Eyak).

**Proto-Eskimo**. **uŋu-ma-n* (Fortescue et al. 2010: 410), derived from **uŋu-ma-* 'to be alive' with the instrumental suffix **-(u)n*. Formally this is the best candidate, well attested in all branches. In some Yupik lects, it was superseded with the stem **irca-quʁ* (Fortescue et al. 2010: 157).

**Proto-Aleut**. **kanuː-χ* (Bergsland 1994: 229; Golovko 1994: 265), attested in all branches.

**Proto-Chukotian**. **liŋ-liŋ* (Fortescue 2005: 159), reduplicated stem, retained in all languages. Cognate to the Itelmen term.

**Proto-Itelmen**. **liŋ-* (Volodin, Khaloimova 2001: 204; Fortescue 2005: 159). Cognate to the Chukotian term.

**Proto-Nivkh**. **ŋiv* (Fortescue 2016: 125; Savelyeva, Taksami 1965: 385; Savelyeva, Taksami 1970: 232).

**Proto-Samoyed**. **säyə̆*, retained in all Samoyed languages, goes back to Proto-Uralic **sʸäðä* 'heart'.

**Proto-Yukaghir**. **suɣo-nčʸə* (Nikolaeva 2006: 404) is retained in both Yukaghir languages. Our reconstruction differs from Nikolaeva's one (**sewe-*), being closer to actual reflexes of this root in daughter languages.

41. 'horn'.

**Proto-Yeniseian**. **qɔʔ* (S. Starostin 1995: 303). Preserved in all daughter languages where attested, but not found in Arin or Pumpokol.

**Proto-Athabaskan**. **teː* is retained in all three branches.

**Eyak.** *=təleh*, morphologically looks like a compound *=tə-leh*, whose first element corresponds to Athabaskan **teː*, whereas the second one remains unclear.

**Proto-Athabaskan-Eyak**. **teː* (Athabaskan, Eyak?).

**Proto-Eskimo**. **naɣðuɣ* (Fortescue et al. 2010: 227), attested in Inuit only, morphologically unclear, can be a descriptive form with the archaic and non-productive suffix **-yuɣ* ~ **-ðuɣ* 'thing resembling smth.' (Fortescue et al. 2010: 481). It was superseded with **ciʁu-nəʁ* in Yupik-Sirenik (Fortescue et al. 2010: 93), a transparent new formation < **ciʁu-* 'to cover' + nominalizer **-nəʁ*.

**Proto-Aleut**. **tumɣa-χ* (Bergsland 1994: 405), attested in Eastern and Atkan, polysemy 'walrus tusk / horn, antlers'.

**Proto-Chukotian**. **rətnə* (Fortescue 2005: 259), retained in all languages. Cognate to the Itelmen term.

**Proto-Itelmen**. **əntəŋ* (Volodin, Khaloimova 2001: 200; Fortescue 2005: 259). Cognate to the Chukotian term.

**Proto-Nivkh**. **murki* (Fortescue 2016: 109; Savelyeva, Taksami 1965: 372; Savelyeva, Taksami 1970: 197).

**Proto-Samoyed**. **amtǝ̆* (Janhunen 1977: 20) is retained in all Samoyed languages.

**Proto-Yukaghir**. **önm-uð* (Nikolaeva 2006: 332–333) is retained in both Yukaghir languages.

42. 'I'.

**Proto-Yeniseian**. **ʔaʒ* (S. Starostin 1995: 185). Preserved in all daughter languages (Ket-Yugh **ʔad*, Kott, Arin *ai*). Reconstruction of the final consonant is questionable. The correspondence is interpreted by S. Starostin as a reflexation of the rare Proto-Yeniseian phoneme **-ʒ* in word-final position, but in reality it is practically indistinguishable from word-final **-ǯ*, so that the reconstruction might ultimately be amended to **ʔaǯ*. However, the final consonant was almost certainly an affricate of some kind.

**Proto-Athabaskan**. **šiː ~ *šį̆ː* is retained in all three branches. The majority of languages point to PA **š*, although some languages have the same reflex as of **š*$_{\iota}$(Krauss 1977: 33).

**Eyak.** *xuː*, apparently related to **šiː ~ *š*$^{w}$*iː* 'I' < PAE **x*$^{w}$*iː*, although the expected reflex of PAE **x*$^{w}$ should be PA **š*$_{\iota}$ (or **š*$^{w}$ in a more traditional notation, Leer 2010). Krauss (1977), however, demonstrated that in the case of the fricative, simple **š* is a more frequent Athabaskan outcome of PAE **x*$^{w}$ than **š*$_{\iota}$

**Proto-Athabaskan-Eyak**. **x*$^{w}$*iː* (Athabaskan-Eyak).

**Proto-Eskimo**. **uva=ŋa ~ *uvi=ŋa* (Fortescue et al. 2010: 418), retained in all branches. In some Yupik lects, it was simplified and lost the nasal element. Apparently the same pattern as in Aleut, i.e., a desemanticized proclitic **uva-* attached to the meaningful pronominal morphemes: **uva=ŋa* 'I', **uva=ku-t* 'we' (q.v.). The same **-ŋa* is used as the 1st p. sg. subject exponent in verbal forms (Fortescue et al. 2010: 489). Cognate to the Aleut pronominal morpheme.

**Proto-Aleut**. **ti=ŋ* (Bergsland 1994: 397; Bergsland 1997: 57), attested in all branches. The meaningful element is *-ŋ*, whereas *ti- ~ txi-* is a proclitic attached to the 1st, 2nd and reflexive 3rd person pronouns (Bergsland 1997: 56). The same **-ŋ* is used as the 1st p. sg. exponent in verbal forms. Cognate to the Eskimo pronominal morpheme.

**Proto-Chukotian**. **kǝm-* (Fortescue 2005: 146), retained in all languages. Cognate to the Itelmen term.

**Proto-Itelmen**. **kǝma* (Volodin, Khaloimova 2001: 236; Fortescue 2005: 147). Cognate to the Chukotian term.

**Proto-Nivkh**. **ɲi* (Fortescue 2016: 114; Gruzdeva 1998: 25; Panfilov 1962: 231).

**Proto-Samoyed**. **mǝ̆-n* (Janhunen 1977: 86), retained in all Samoyed languages, goes back to Proto-Uralic **mi-n* 'I'.

**Proto-Yukaghir**. **mǝ-t* (Nikolaeva 2006: 267) is retained in both Yukaghir languages.

43. 'to kill'.

**Proto-Yeniseian**. **ʔeːy ~ *xeːy* (S. Starostin 1995: 190). Attested only in Ket-Yugh; not attested in either Arin or Pumpokol, and most likely replaced in Kott. Since the word is known only from Ket-Yugh, word-initial zero could just as well have been **x-*.

**Proto-Athabaskan**. **=ʁeː* is retained in all three branches.

**Eyak.** *=še*, the Eyak paragim can point to either a PAE **CVː*-root or a **CVːn*-root.

**Proto-Athabaskan-Eyak**. **=χeː* (Athabaskan), **=šV ~ *=šVn* (Eyak).

**Proto-Eskimo**. **tuqu-t-* (Fortescue et al. 2010: 386), retained in all branches. A regular causative from **tuqu-* 'to die' (q.v.).

**Proto-Aleut**. **asχa-t-* (Bergsland 1994: 100; Golovko 1994: 279), attested in all branches. A regular causative from **asχa-l-* 'to die' (q.v.).

**Proto-Chukotian**. **tǝm-* (Fortescue 2005: 297), retained in all languages. Cognate to the Itelmen term.

**Proto-Itelmen**. **ɬmǝ-* (Volodin, Khaloimova 2001: 53, 216; Fortescue 2005: 297). Cognate to the Chukotian term.

**Proto-Nivkh**. **ku-* (Fortescue 2016: 88; Savelyeva, Taksami 1965: 429; Savelyeva, Taksami 1970: 90).

**Proto-Samoyed**. **kɒ̆ə̆-tɒ-* (Janhunen 1977: 57), retained in all daughter languages, is a causative of Proto-Samoyed **kɒə̆-* 'to die' (< Proto-Uralic **kali-* 'to die').

**Proto-Yukaghir**. **punʸ-* (Nikolaeva 2006: 370) is attested in Tundra Yukaghir. The meaning of Kolyma *kudedə-* 'to kill' is evidently secondary: its Tundra cognate means 'to put down' (Nikolaeva 2006: 220–221).

44. 'knee'.

**Proto-Yeniseian**. **baʔt* (S. Starostin 1995: 206). Preserved everywhere except for Kott; not attested in Pumpokol. Replaced in Kott with *arša*, an etymologically obscure form. The root **baʔt* per se must have had the general meaning 'joint' in Proto-Yeniseian, but this does not technically prevent us from setting up **baʔt* as the main bearer of the meaning 'knee' as well for Proto-Yeniseian.

**Proto-Athabaskan**. **=qVt'* is retained in all three branches.

**Eyak.** *=quht ~ =qũht*, cognate to Athabaskan.

**Proto-Athabaskan-Eyak**. **qVnt'* (Athabaskan, Eyak), we follow Leer 2008a and reconstruct a medial nasal on account of vowel nasalization and aspiration in Eyak.

**Proto-Eskimo**. **ciɣəð-quʁ* (Fortescue et al. 2010: 81), retained in all branches. The final element is the suffix **-quʁ* 'smth. associated with or attached to smth.'. Cognate to the Aleut term via metathesis.

**Proto-Aleut**. **čiðiɣi-χ* (Bergsland 1994: 136; Golovko 1994: 218), attested in Atkan and, with an additional suffix, in Eastern and Attuan. Cognate to the Eskimo term via metathesis.

**Proto-Chukotian**. **ŋəra-* (Fortescue 2005: 202), retained in all languages except for Kerek.

**Proto-Itelmen**. **sʸizʸə-* (Volodin, Khaloimova 2001: 162; Fortescue 2005: 373).

**Proto-Nivkh**. **biɣ-tV* (Fortescue 2016: 22; Savelyeva, Taksami 1965: 183; Savelyeva, Taksami 1970: 262).

**Proto-Samoyed**. **puə̆* (Janhunen 1977: 130), retained in all languages save Kamass, is related to Finno-Ugric words for 'knee'.

**Proto-Yukaghir**. **poɣo- ~ *poŋqə-* (Nikolaeva 2006: 354) is attested in Kolyma Yukaghir. Tundra word for 'knee' means literally 'joint bone'.

45. 'to know'.

**Proto-Yeniseian**. Dubious reconstruction: Ket-Yugh **ʔit-* is probably the best candidate for the PY slot, but even in Ket-Yugh the complex structure of this verb is not thoroughly understood, and it has no external parallels. Kott *ŋ=aːliga*, structured more like a nominal than a verbal formation, is even more obscure.

**Proto-Athabaskan**. The main candidates are: **=c'Vt* (Pacific Coast), **=nVy* (Northern), **=zen* (Apachean). Out of them, Apachean **=zen* < 'to think' as proven by its cognates in other branches, whereas **=nVy* denotes a wide range of sensory perceptions and activities, it is thus likely that 'to know' was not the primary meaning of **=nVy*. So **=c'Vt* is the best candidate.

**Eyak.** *=ka,* cognate to Tlingit *=keː* 'to understand' and potentially to PA *=čax ~ =čąx* 'to try, test'.

**Proto-Athabaskan-Eyak**. **=c'Vt* (Athabaskan), **=kaː* (Eyak).

**Proto-Eskimo**. **əli-ci-ma-* (Fortescue et al. 2010: 116), retained in all branches; final **-ma-* is the perfective suffix. The same root is used in such Eskimo stems as **əli-ma-* 'to be knowledgeable; to be apprehensive' or **əli-t-* 'to learn' (Fortescue et al. 2010: 115). Also negated forms of **nału-* 'to not know' (Fortescue et al. 2010: 231) can be used for the positive meaning 'to know'.

**Proto-Aleut**. **iða-χta-laka-* (Bergsland 1994: 171; Golovko 1994: 49, 213), attested in Atkan and Attuan, meaning 'to know a situation or fact / to understand'. Cf. some Bergsland's examples: "did you know that it is foggy?", "his wife knows for sure that he'll be back", etc. The complex stem *iða-χta-laka-* literally means 'not to have (smth.) unknown' with the negative exponents *-laka-* and the bound root *iða-* 'unknown, unclear *vel sim.*' (Bergsland 1994: 171). Distinct from the verb *haqa-t-a-l-* (Bergsland 1994: 94; Golovko 1994: 158, 213), which means 'to know, be acquainted with object/person, have skill to' in Atkan and Attuan. Cf. some Bergsland's examples: "I don't know him", "he does not know how to hunt seals", etc. The stem *haqa-t-a-l-* is derived from causative *haqa-t-* 'to bring, give; to come upon, find; to find out, learn', finally from *haqa-l-* 'to come' (Bergsland 1994: 93). Note that in the Eastern dialect, only *haqa-t-a-l-* is used for both meaning, apparently this situation is secondary.

**Proto-Chukotian**. *ləɣi ləŋ-* (Fortescue 2005: 162), an analytic construction with *ləɣi* 'known' and auxiliary *ləŋ-* 'to consider as', retained in all languages as the basic expression for 'to know (in general)'.

**Proto-Itelmen**. *xaq* AUX (Volodin, Khaloimova 2001: 157; Fortescue 2005: 373). An analytic construction 'to know (in general)', lit. 'to be *xaq*'.

**Proto-Nivkh**. *haym-* (Fortescue 2016: 70; Savelyeva, Taksami 1965: 162; Savelyeva, Taksami 1970: 470).

**Proto-Samoyed**. *tänä-mä-* (Janhunen 1977: 157) is attested in most daughter languages. The word is derived from Proto-Samoyed *tänä-* 'to remember'.

**Proto-Yukaghir**. Kolyma *ley-*, whose Tundra cognate means 'to remember' (Nikolaeva 2006: 238) vs. Tundra *kuril$^{y}$-* (Nikolaeva 2006: 229).

46. 'leaf'.

**Proto-Yeniseian**. *yə:pe* (S. Starostin 1995: 232). Preserved in all daughter languages.

**Proto-Athabaskan**. *=t'a:n-ʔ* is retained in all three branches.

**Eyak.** *t'ahɬ*, cognate to Athabaskan, historically should be analyzed as *t'ah-ɬ* with the common nominal suffix *-ɬ*.

**Proto-Athabaskan-Eyak**. *t'a:n* (Athabaskan, Eyak).

**Proto-Eskimo**. An unstable item. The best candidate is *pəɬu* (Fortescue et al. 2010: 279), which is attested as 'leaf' in both Yupik and Inuit. Theoretically it may be cognate to Aleut *huli-χ* 'leaf'.

**Proto-Aleut**. An unstable item. The best candidate is *huli-χ* attested in the Eastern dialect from the 18th c. on and in Attuan (Bergsland 1994: 435). In archaic Atkan, another word, *siʁli-χ* 'leaf', is attested from the 18th c. on (Bergsland 1994: 359). Cf. also the modern term with irregular sound fluctuation *yuʁli-χ ~ yuχli-χ ~ yuli-χ* 'leaf' (Bergsland 1994: 465; Golovko 1994: 223).

**Proto-Chukotian**. *wət-wət* (Fortescue 2005: 337), retained in all languages except for Kerek. Reduplicated stem, based on the root 'green'.

**Proto-Itelmen**. *palla-* (Volodin, Khaloimova 2001: 168; Fortescue 2005: 374).

**Proto-Nivkh**. *ɖom-r* (Fortescue 2016: 53; Savelyeva, Taksami 1965: 199; Savelyeva, Taksami 1970: 350). Attested in all languages. The second candidate is *blaŋ* 'leaf' which has a narrower distribution (Fortescue 2016: 23).

**Proto-Samoyed**. *yapä* (Janhunen 1977: 41) is retained in Northern Samoyed, Kamass and Selkup.

**Proto-Yukaghir**. Kolyma *piw-il* (Nikolaeva 2006: 353) vs. Tundra *pöɣ-il* (Nikolaeva 2006: 354). The modern Kolyma term for 'leaf' goes back to *polčičə* (Nikolaeva 2006: 356–357), but at least four old Kolyma wordlists from 18th and 19th centuries have reflexes of *piw-il* (whose modern reflexes in Kolyma and Tundra mean 'needle of a coniferous tree') glossed as 'leaf'. We take it as an indication that *piw-il* was replaced by *polčičə* in modern Kolyma. Tundra *pug-il* (< *pöɣ-il*) is glossed as 'leaf; widow(er)', but this is most likely a chance homonymy.

47. ‘to lie’.

**Proto-Yeniseian**. **=qɔt* (S. Starostin 1995: 183 as **ʔaq-ɔt-*, with probably incorrect segmentation). Preserved in Ket-Yugh and Kott. In Arin, definitely preserved in the meaning ‘sleep’ q.v., but not attested in the meaning ‘lie’; uncertain situation in Pumpokol (Pumpokol *ak* is an unclear form without any obvious parallels). The verb **=qɔt* was most likely polysemous in Proto-Yeniseian, meaning both ‘to lie’ and ‘to sleep’. The paradigm must have been suppletive, since Ket-Yugh **=dam-* in plural forms corresponds to Kott *=tam-* in such forms as *dʸ=a=tam-an-toŋ* ‘we lie / we sleep’, etc. The opposition **=qɔt* [sg.] / **=dam-* [pl.] is thus safely reconstructible, although Kott shows no signs of the directional prefix *t=*, obligatory in Ket-Yugh.

**Proto-Athabaskan**. **=tʰeː-(ɲ)* is retained in all three branches.

**Eyak.** *=tʰe*, cognate to Athabaskan.

**Proto-Athabaskan-Eyak**. **=tʰeː* (Athabaskan, Eyak).

**Proto-Eskimo**. **in-naʁ* ~ **iŋ-naʁ* (Fortescue et al. 2010: 149), retained in all branches. The final element is the assimilated durative suffix known in many forms: **-aʁ, *-ðaʁ, *-laʁ, *-maʁ, *-taʁ*, etc. In Yupik, it usually means ‘to lie, lie down’. In the majority of Inuit lects, shifted into such specific meanings as ‘lie down on side’ or ‘to go to bed’, having been superseded with **nala-* (Fortescue et al. 2010: 229), whose original meaning is not entirely clear (‘to be lying down (of plant)’?).

**Proto-Aleut**. **quyu-ʁi-l-* (Bergsland 1994: 340; Golovko 1994: 222), attested as a generic verb for ‘to lie (of human)’ in Eastern (from the 19th c. on) and Atkan (from the 18th c. on). Derived from **quyu-l-* ‘to go to bed, lie down’. The second and less probable candidate is **aŋa-mi-l-* ‘to lie down, be in lying position (on the side, or in general)’ (Bergsland 1994: 83) which is a synonym of **quyu-ʁi-lix*, but is restricted to the Eastern dialect; derived from **aŋa-* ‘side, lateral part’.

**Proto-Chukotian**. **rəl-tel-* (Fortescue 2005: 256), retained as basic ‘to lie’ in all languages except for Kerek. Cognate to the Itelmen term.

**Proto-Itelmen**. **sʸol-* (Volodin, Khaloimova 2001: 167; Fortescue 2005: 256). Cognate to the Chukotian term.

**Proto-Nivkh**. **bor-* (Fortescue 2016: 25; Savelyeva, Taksami 1965: 197; Savelyeva, Taksami 1970: 267), polysemy ‘to lie / to lie down’.

**Proto-Samoyed**. **kɨy-tV-* (Helimski 1997: 280–281), going back to Proto-Uralic **kuyi-* ‘to lie’, is retained in Mator.

**Proto-Yukaghir**. **qont-oː-* (Nikolaeva 2006: 220–221), retained in both Yukaghir languages, is derived from the root **köntə-*. The word is connected to Proto-Samoyed **kont-ö-* ‘to sleep’ (Janhunen 1977: 73), but the direction of borrowing is not clear (for phonological reasons common inheritance is unlikely in this case).

48. ‘liver’.

**Proto-Yeniseian**. **seŋ* (S. Starostin 1995: 272). Preserved only in Ket-Yugh. Kott *šičil* and Arin *sal* are most likely related and go back to Kott-Arin **sisal* ‘internal organ’, a form with no transparent internal etymology and vague semantics. In this context, Ket-Yugh **seŋ* is a more reliable candidate for Proto-Yeniseian ‘liver’.

**Proto-Athabaskan**. **=zet’* is retained in all three branches.

**Eyak.** *=saht*, cognate to Athabaskan.

**Proto-Athabaskan-Eyak**. **sVnt’* (Athabaskan, Eyak). We follow Leer 2008a and reconstruct a medial nasal on account of vowel aspiration in Eyak.

**Proto-Eskimo**. **təŋu-ɣ* (Fortescue et al. 2010: 373), retained in all branches.

**Proto-Aleut**. **aːʁi-χ* (Bergsland 1994: 38; Golovko 1994: 243), attested in all branches.

**Proto-Chukotian**. **ponta* (Fortescue 2005: 218), retained in all languages. Cognate to the Itelmen term.

**Proto-Itelmen**. **ponta* (Volodin, Khaloimova 2001: 186; Fortescue 2005: 218). Cognate to the Chukotian term.

**Proto-Nivkh**. **div-r* (Fortescue 2016: 42; Savelyeva, Taksami 1965: 292; Savelyeva, Taksami 1970: 353). Looks like a deverbative in **-r*.

**Proto-Samoyed**. **mitǝ̆* (Janhunen 1977: 93–94), retained in all Samoyed languages save Mator, goes back to Proto-Uralic **miksa* 'liver'.

**Proto-Yukaghir**. Kolyma **kuðenčʸǝ* (Nikolaeva 2006: 225) vs. Tundra **alʸaːyǝ* (Nikolaeva 2006: 100).

49. 'long'.

**Proto-Yeniseian**. **ʔux-* (S. Starostin 1995: 201). Preserved in all daughter languages, but not attested in Pumpokol.

**Proto-Athabaskan**. **=ɲeːs* is retained in all three branches.

**Eyak.** *=ʔaːw.*

**Proto-Athabaskan-Eyak**. **=ɲeːs* (Athabaskan), **=ʔaːw* (Eyak).

**Proto-Eskimo**. **takǝ-* (Fortescue et al. 2010: 355), retained in all branches.

**Proto-Aleut**. **aðu-* (Bergsland 1994: 14; Golovko 1994: 202), attested in all branches.

**Proto-Chukotian**. **iwlǝ-* (Fortescue 2005: 106), retained as basic 'long' in all languages except for Kerek. Cognate to the Itelmen term.

**Proto-Itelmen**. **iwl-* (Volodin, Khaloimova 2001: 149; Fortescue 2005: 106). Cognate to the Chukotian term (if not a Chukotian loan).

**Proto-Nivkh**. **gǝl-* (Fortescue 2016: 64; Savelyeva, Taksami 1965: 128; Savelyeva, Taksami 1970: 126).

**Proto-Samoyed**. **yɒmpǝ̆* (Janhunen 1977: 37) is retained in all daughter languages save Nganasan.

**Proto-Yukaghir**. **čit-nǝ-* (Nikolaeva 2006: 134) is retained in both Yukaghir languages.

50. 'louse'.

**Proto-Yeniseian**. **ʔǝːke* ~ **xǝːke* (S. Starostin 1995: 192). Preserved in both of the primary Yeniseian branches (including Ket-Yugh and Kott). Lack of Arin parallels means that either **ʔ-* or **x-* were present in the word-initial position.

**Proto-Athabaskan**. **yaʔ* ~ **yaː* is retained in all three branches.

**Eyak.** *kuks-k.*

**Proto-Athabaskan-Eyak**. **yaː* (Athabaskan), **kVks* (Eyak).

**Proto-Eskimo**. **kuma-ɣ* (Fortescue et al. 2010: 198), retained in all branches.

**Proto-Aleut**. **kitu-χ* (Bergsland 1994: 242), attested in Eastern and Atkan.

**Proto-Chukotian**. **mǝl-mǝl* (Fortescue 2005: 183), reduplicated stem, retained in all languages. Cognate to the Itelmen term.

**Proto-Itelmen**. **mil-mil* (Ono 2003: 102; Fortescue 2005: 183), reduplicated stem. Cognate to the Chukotian term.

**Proto-Nivkh**. An unclear situation. There are two terms attested in Amur and North Sakhalin, namely morphologically unclear **amrak* 'head louse' (Fortescue 2016: 13; Savelyeva, Taksami 1965: 95; Savelyeva, Taksami 1970: 32) and **dar* 'body louse' (Fortescue 2016: 40; Savelyeva, Taksami 1970: 344). The third term **hirk* 'louse' is attested in East and South Sakhalin (Fortescue 2016: 74; Savelyeva, Taksami 1970: 344), it can be a generic term for 'louse', although, strictly speaking, the data in Savelyeva, Taksami 1970 suggest the specific meaning 'body louse' for **hirk*. The proto-term for 'nit' is derived from the latter root: **hirk-r* 'nit' (Fortescue 2016: 75). Provisionally we treat **amrak* and **hirk* as synonyms.

**Proto-Samoyed**. **ɒncu̯* (Janhunen 1977: 18) is attested in Forest Nenets, Enets, Nganasan, Kamass and Selkup, whereas **pǝ̆nsV* (Helimski 1997: 246) is attested in Tundra Nenets and Mator with derivatives in Forest Nenets and Enets. It seems clear that both roots were present in Proto-Samoyed, but the semantic difference between them is not clear.

**Proto-Yukaghir**. **peme ~ *pime* (Nikolaeva 2006: 348) is retained in both Yukaghir languages.

51. ‘man (male human being)’.

**Proto-Yeniseian**. **pixe* (S. Starostin 1995: 249). Preserved in all daughter languages where attested, but not found in Pumpokol, and dubious in Arin.

**Proto-Athabaskan**. **te=ne:* ‘man / person’ is the main candidate (the initial element is a desemanticized nominal prefix).

**Eyak.** *łi=la:ʔ*, morphologically unclear, either a deverbative from *=la:ʔ* ‘?’ (thus a secondary formation) or the second element is a direct cognate to Athab. **=ne:*.

**Proto-Athabaskan-Eyak**. **nV:* (Athabaskan, Eyak?).

**Proto-Eskimo**. **aŋu-n* (Fortescue et al. 2010: 38), retained in all branches, although it was superseded with **inʸu-ɣ* ‘person’ (q.v.) in some Yupik lects.

**Proto-Aleut**. **taya-ʁu-χ* (Bergsland 1994: 395; Golovko 1994: 227), attested in all branches, the starting root is unclear, perhaps literally ‘the one with many *taya*’.

**Proto-Chukotian**. **qəlavol* (Fortescue 2005: 243), retained as a basic word for ‘man’ in all languages except for Koryak. Morphologically unclear.

**Proto-Itelmen**. **iχłχ* (Fortescue 2005: 244; Volodin, Khaloimova 2001: 172; Ono 2003: 110).

**Proto-Nivkh**. There are two terms for ‘man’ competing with each other. First, Amur and North Sakhalin **ut-kun* (Fortescue 2016: 158; Savelyeva, Taksami 1965: 214; Savelyeva, Taksami 1970: 397), formally looks like a regular plural form in **-kun*, thus can be an old collective term, or it can be related to the anthroponymic suffixes of the shape *-kun* for which see Panfilov 1962: 52 (for the starting root cf. **ut* ‘body’, Fortescue 2016: 163). Second, East and South Sakhalin **ar-mət̆* (Fortescue 2016: 16; Savelyeva, Taksami 1970: 397), derived from **ar* ‘male’ (Fortescue 2016: 15), the second element is probably related to the verb *mu-* ‘to become’, thus Panfilov 1962: 81. We treat **ut-kun* and **ar-mət̆* as synonyms.

**Proto-Samoyed**. **tepä* (Janhunen 1977: 163) is retained in Mator, Kamass and Selkup. Nganasan has a derivative with the meaning ‘boy’. In Northern Samoyed this word was replaced by Proto-Samoyed **kɒə̑-sɒ* ‘person’.

**Proto-Yukaghir**. **köy* (Nikolaeva 2006: 215–216) is retained in both Yukaghir languages.

52. ‘many’.

**Proto-Yeniseian**. **bəy-* (S. Starostin 1995: 209). Preserved in Kott, but possibly still active in its original meaning in mid-19th century Ket as well. Attestation of Ket *bзäyäm* ‘many’ in Castrén’s records, clearly related to Kott *payaŋ*, shows that the modern Ket descendant of this proto-item, *bзyaŋ* ‘enough’, may have undergone a semantic shift.

**Proto-Athabaskan**. **ła:ɲ* is retained in all three branches.

**Eyak.** *=t’uʔ*, note the Athabaskan verb **=t’e:* ‘to be thus, be in circumstances of’ on which expressions for ‘many’ are based in some Athabaskan languages. Theoretically Eyak *=t’uʔ* and Athabaskan **=t’e:* can be cognates.

**Proto-Athabaskan-Eyak**. **ła:ɲ* (Athabaskan), **=t’V* (Eyak).

**Proto-Eskimo**. **amə-cu- ~ *amə-lʁa-* (Fortescue et al. 2010: 24–25), retained in all branches, although suffix extensions are not entirely clear.

**Proto-Aleut**. **qala-χ* (Bergsland 1994: 302), attested in Eastern and Atkan; a more widely used expression is suffixed **qala-ʁi-l-* (Bergsland 1994: 302; Golovko 1994: 74), attested in all branches. Tends to be superseded with the new formation **hasi-na-l-* ‘to be many’ (Bergsland 1994: 101; Golovko 1994: 161), derived from the bound root **hasi-* ‘crowd *vel sim.*’.

**Proto-Chukotian**. **mək-* (Fortescue 2005: 181), retained in all languages. Distinct from **ŋənvəq* ‘much’ (Fortescue 2005: 202).

**Proto-Itelmen**. **pto-* (Volodin, Khaloimova 2001: 73; Volodin 1976: 320; Fortescue 2005: 226). Glossed specifically as ‘much’ in Fortescue 2005, but apparently it is a common basic term for ‘many / much’, thus Volodin 1976: 320. Another candidate is **meyim ~*

*meyiŋ (Volodin, Khaloimova 2001: 58; Fortescue 2005: 171), explained as 'enough, many/much, thoroughly' in Volodin 1976: 344. Note that Fortescue supposes that *meyim ~ *meyiŋ is a loan from Chukotian *mäyəŋ- 'big' (q.v.), but it is not very likely due to semantic difference.

**Proto-Nivkh**. *mal-ɣo- (Fortescue 2016: 101; Savelyeva, Taksami 1965: 210; Savelyeva, Taksami 1970: 173; Taksami 1996: 147). Attested everywhere; a basic expression for '(to be) many' at least in Amur. The second candidate in *dam- (Fortescue 2016: 38; Savelyeva, Taksami 1970: 342), which is attested with the meaning '(to be) many' in Amur, North and East Sakhalin.

**Proto-Samoyed**. *oykkɒ ~ *oytkɒ ~ *oycḳɒ ~ *oyskɒ (Janhunen 1977: 29) is retained in most daughter languages.

**Proto-Yukaghir**. Kolyma *ninkə- (Nikolaeva 2006: 303) vs. Tundra *poy-oː- (Nikolaeva 2006: 355).

53. 'meat'.

**Proto-Yeniseian**. *ʔise (S. Starostin 1995: 194). Preserved in all daughter languages except for Pumpokol, where it was replaced with *cič* = Kott *šig* 'food' < Proto-Yeniseian *siː-k 'food', a nominal derivative from *siː- 'to eat' q.v.

**Proto-Athabaskan**. *=cʰeɲʔ is retained in all three branches.

**Eyak.** =cʰeʔ, cognate to Athabaskan, final -ʔ prevents the expected aspiration of *e*.

**Proto-Athabaskan-Eyak**. *cʰeɲ (Athabaskan, Eyak).

**Proto-Eskimo**. *nəqə (Fortescue et al. 2010: 251), retained in all branches, usually polysemy: 'food / meat'.

**Proto-Aleut**. *ulu-χ (Bergsland 1994: 436; Golovko 1994: 228), attested in all branches.

**Proto-Chukotian**. *tərɣe-tər (Fortescue 2005: 301), partial reduplication, a basic term at least in Chukchi (Inenlikei 2005) and Alutor (Kibrik et al. 2004: 558). Superseded with *kinuŋi (Zhukova 1990: 155; Fortescue 2005: 138) in Koryak, whose Alutor cognate means specifically 'reindeer meat' (Nagayama 2003: 267).

**Proto-Itelmen**. *tχal-tχal (Volodin, Khaloimova 2001: 173; Fortescue 2005: 302), reduplicated stem.

**Proto-Nivkh**. *dur ~ *ḍur (Fortescue 2016: 47; Savelyeva, Taksami 1965: 216; Savelyeva, Taksami 1970: 373).

**Proto-Samoyed**. *ɒyɒ (Janhunen 1977: 17) preserves the sense 'meat / flesh' in Kamass and Selkup. In Northern Samoyed this word retains only the meaning 'flesh', being replaced in the sense 'meat' by *ə̆m-sɒ (originally 'food', derived from *ə̆m- 'to eat'). In Mator, the root *ɒyɒ is also replaced by *ə̆m-sɒ, being preserved only in the derivate 'raw'.

**Proto-Yukaghir**. *čuː-l (Nikolaeva 2006: 143) is retained in both Yukaghir languages.

54. 'moon'.

**Proto-Yeniseian**. The etymon *suy (S. Starostin 1995: 204) is seen in Kott-Arin and Pumpokol; in Ket-Yugh, the equivalent of 'moon' is *qip, of unclear origin; the most tempting solution would be to identify it with *qib 'grandfather', but the idea runs into significant phonetic problems, unless one can come up with a satisfactory solution for the irregular devoicing of the final consonant in the word for 'moon'. We include both words into comparison for extra safety.

**Proto-Athabaskan**. *ša̧ː 'sun / moon' is retained in all three branches, although, in the meaning 'moon', this root tends to be superseded with various descriptive formation.

**Eyak.** qʰə=χah, perhaps contains the same root as ʔiš=χah 'round howl, (round-bottomed) mixing-howl'.

**Proto-Athabaskan-Eyak**. Not reconstructible, because both Athabaskan and Eyak terms seem secondary. A possible candidate is PAE *nen (> PA *nen 'month', Eyak *leh* 'year'), since 'moon' is the main semantic source for the meaning 'month' and it is not a rare situation when an innovative term acquires the meaning 'moon', whereas the old term is retained as 'month'.

**Proto-Eskimo**. **tanqi-* (Fortescue et al. 2010: 360), retained in all branches. Formally this is the best candidate, apparently with Proto-Eskimo polysemy 'light, to be bright / moon'.

**Proto-Aleut**. **tuɣi-ða-χ* (Bergsland 1994: 402; Golovko 1994: 224), attested in all branches.

**Proto-Chukotian**. **yəʁ=ilɣən* (Fortescue 2005: 124), retained in all languages. Literally 'cloud's whiteness'(?) with **yəʁə-n* 'cloud' (q.v.) and **ilɣə* 'white' (q.v.).

**Proto-Itelmen**. Apparently the stem **kulač* (Fortescue 2005: 393), which is normally attested with the meaning 'sun', is to be reconstructed with the areal polysemy 'sun / moon'. Firstly, the synchronic polysemy 'sun / moon' is documented for Eastern Itelmen. Secondly, in both Eastern and Southern, 'moon' can be expressed with the collocation 'night **kulač*' *vel sim*. In Western, inherited **kulač* is retained as 'sun', whereas 'moon' is denoted with either the Chukotian loans such as *yeʔalɣən* ~ *yeʔalŋin* 'moon' (Ono 2003: 116; Fortescue 2005: 124) or the Russian loan *mesic* 'moon' (Volodin, Khaloimova 2001: 168).

**Proto-Nivkh**. **loŋ* (Fortescue 2016: 98; Savelyeva, Taksami 1965: 202; Savelyeva, Taksami 1970: 163).

**Proto-Samoyed**. **kɨy* (Janhunen 1977: 69), retained in Nganasan, Mator and Kamass, goes back to Proto-Uralic **kɨwi* 'moon'.

**Proto-Yukaghir**. **kininčʸə* (Nikolaeva 2006: 212) is retained in both Yukaghir languages.

55. 'mountain'.

**Proto-Yeniseian**. **rʸɨʔǯ* (S. Starostin 1995: 267). The situation with Proto-Yeniseian 'mountain' is quite complex. Ket-Yugh **qaʔy* 'mountain; steep bank' corresponds to Kott *xey* ~ *kʰey* 'back side of axe / knife'; the same root is most likely present in Kott *xeː-leːx* ~ *kʰeː-leːg* 'back side of mountain'. The semantic development 'mountain' > 'side of axe / knife' is suspicious; a more likely common invariant would be 'elevation', 'protruding part', etc., implying that the primary semantics of 'mountain' for this root on the Proto-Yeniseian level is not likely. In Ket-Yugh, the word was probably originally applied to 'cliffs' or 'steep riverbanks', then extended to denote 'wood-covered mountains' as well. The original word for 'wood-covered mountain' (the default kind of mountain for Yeniseian territory) must have been **rʸɨʔǯ*. Arin *kar* 'mountain' is isolated in Yeniseian and has no etymological connections whatsoever.

**Proto-Athabaskan**. Northern and Apachean **ceɬ* is the best candidate. Descriptive new formation in the Pacific Coast branch.

**Eyak.** *ʔiƛʼ*.

**Proto-Athabaskan-Eyak**. **ceɬ* (Athabaskan), **ʔiƛʼ* (Eyak).

**Proto-Eskimo**. **iŋʁi-ʁ* (Fortescue et al. 2010: 150), retained in all branches.

**Proto-Aleut**. **kiːʁuː-si-χ* (Bergsland 1994: 238; Golovko 1994: 67), attested in all branches.

**Proto-Chukotian**. **ŋäy* (Fortescue 2005: 194), retained in all languages. Cognate to the Itelmen term.

**Proto-Itelmen**. **ŋey-ŋe* (Volodin, Khaloimova 2001: 145; Volodin 1976: 31, 150, 323; Fortescue 2005: 194). Reduplicated stem. Cognate to the Chukotian term.

**Proto-Nivkh**. **bal* (Fortescue 2016: 20; Savelyeva, Taksami 1965: 118; Savelyeva, Taksami 1970: 250). In Amur, it means specifically 'mountain covered with forest / forest'. Distinct from **ʈir* (Fortescue 2016: 33) which denotes a forestless area: 'forestless mountain' in Amur, 'field' in East Sakhalin.

**Proto-Samoyed**. **wɒrɒ*, retained in Enets, Nganasan and Kamass, goes back to Proto-Uralic **wara* 'mountain'.

**Proto-Yukaghir**. Kolyma **peː* 'mountain / big stone', possibly connected with the Proto-Samoyed word for 'stone' (Nikolaeva 2006: 344–345) vs. Tundra **anaː* (Nikolaeva 2006: 105–106).

56. ‘mouth’.

**Proto-Yeniseian**. **qowe* (S. Starostin 1995: 302). Preserved in all daughter languages.

**Proto-Athabaskan**. **=za-ʔ* and **=ta-ʔ* are the main candidates with the protolanguage opposition ‘interior mouth’ / ‘exterior mouth’.

**Eyak.** *=saʔ*, cognate to Athabaskan.

**Proto-Athabaskan-Eyak**. **sa:* (Athabaskan, Eyak).

**Proto-Eskimo**. **qanə-ʁ* (Fortescue et al. 2010: 309), retained in all branches, frequently with polysemy ‘mouth / to speak’.

**Proto-Aleut**. **aɣi-lʁ-* ~ **aɣa-lʁ-* (Bergsland 1994: 23; Golovko 1994: 261), attested in all branches, sometimes with polysemy: ‘mouth / door’. Derived from **aɣi-l-* ‘to open one’s mouth; to yawn’.

**Proto-Chukotian**. **rək-ərŋə-n* (Fortescue 2005: 256), retained in all languages. The final element is the suffix **-ɣərŋə-n* (cf. Fortescue 2005: 408), attested in some nouns with the semantics of hole.

**Proto-Itelmen**. **qasχ* (Volodin, Khaloimova 2001: 201; Fortescue 2005: 378).

**Proto-Nivkh**. Two terms for ‘mouth’ can be reconstructed: **al* ‘interior mouth’ (Fortescue 2016: 10; Savelyeva, Taksami 1970: 463) and **amɣ* ‘exterior mouth’ (Fortescue 2016: 12; Savelyeva, Taksami 1965: 373; Savelyeva, Taksami 1970: 464). Phonetic similarity with Proto-Tungusic **amŋa* ‘mouth’ (> Orok *amŋa*, Evenki *amŋa*, Nanai *aŋma* ~ *amɢa*, all ‘mouth’) can be accidental.

**Proto-Samoyed**. **aŋ* (Janhunen 1977: 20), retained in all Samoyed languages, goes back to Proto-Uralic **aŋi* ‘mouth / opening’.

**Proto-Yukaghir**. **aŋa* (Nikolaeva 2006: 106) is retained in both Yukaghir languages. It is cognate to, or borrowed from, Proto-Samoyed **aŋ* ‘mouth’.

57. ‘name’.

**Proto-Yeniseian**. **ʔiɢ* (S. Starostin 1995: 193). Preserved in all daughter languages, although not attested in Arin (the Pumpokol form could also, in theory, be Yugh rather than Pumpokol). Root-final **-ɢ* reconstructed based on its complete disappearance in Ket-Yugh (**i:*) but preservation in Kott (*ix*).

**Proto-Athabaskan**. **=u:=že-ʔ* is retained in all three branches.

**Eyak.** *wə=šeh*, cognate to Athabaskan and to Tlingit *sa:* ‘name’, although the Eyak aspirated phonation should point to an additional *n*-suffix in Proto-Eyak.

**Proto-Athabaskan-Eyak**. **we=še* (Athabaskan, Eyak).

**Proto-Eskimo**. **atə-ʁ* (Fortescue et al. 2010: 55), retained in all branches. Cognate to the Aleut term.

**Proto-Aleut**. **asa-χ* (Bergsland 1994: 96; Golovko 1994: 215), attested in all branches. Cognate to the Eskimo term.

**Proto-Chukotian**. **nənnə* (Fortescue 2005: 191), retained in all languages.

**Proto-Itelmen**. **χela-ŋ* (Volodin, Khaloimova 2001: 159; Fortescue 2005: 379).

**Proto-Nivkh**. **qa* (Fortescue 2016: 138; Savelyeva, Taksami 1965: 169; Savelyeva, Taksami 1970: 146).

**Proto-Samoyed**. **nim* (Janhunen 1977: 102), retained in all Samoyed languages, goes back to Proto-Uralic **nimi* ‘name’.

**Proto-Yukaghir**. Kolyma **nʸi-u:* (Nikolaeva 2006: 312) vs. Tundra **kiriyə* (Nikolaeva 2006: 213). Kolyma word is traditionally compared with Proto-Uralic **nimi* ‘name’ and its Samoyed reflex **nim*. Early attestations of the Yukaghir word include *nim* in the 17th century Yukaghir translation of the Lord’s prayer and the 18th and 19th century forms like *nywa, niiv* and *niw*. Modern Kolyma has *nʸu:*. Based on the earliest attestation, Nikolaeva postulates a development **nime* > *niwe* > *niw* > *nʸu:*. The reconstruction with **m* cannot be correct, because there are plenty of Proto-Yukaghir words with word-internal **m*, which is regularly preserved in all daughter idioms. It is much more probable that the form *nim* results from a misprint or some other kind of error. We suggest that *n$^y$u:* can be etymologized as a derivative from the Proto-Yukaghir verb **n$^y$e:-* 'call, call by name' (Nikolaeva 2006: 292), formed with the deverbal noun suffix *-u:*. Older forms like *niw* preserve the root vowel (some derivatives from **n$^y$e:-* have an allomorph **n$^y$i-*). Thus, despite superficial similarity, the Kolyma word for 'name' has nothing to do with Uralic **nimi*.

**Proto-Burushaski**. We reconstruct the Proto-Burushaski form as **=yek* ~ **=ek* (> Yasin *=y'ek*, Hunza *='ik*) for S. Starostin's **yek*, since initial **y-* is generally expected to be retained in Hunza.

58. 'neck'.

**Proto-Yeniseian**. **kəqənt* (S. Starostin 1995: 237). Preserved in Ket-Yugh. In Kott-Arin, replaced with **puyme* ~ **puymur*, of unclear origin. The reason why the Ket-Yugh word is seen as more archaic is the Kott parallel in *agantan* 'collar' (< **kagantan* with dissimilation): the semantic development 'neck' > 'collar' is typologically normal, whereas the opposite would be quite strange. Due to its sheer length, Proto-Yeniseian **kəqənt* must have contained a suffix, although the element **-nt* is hardly segmentable as a productive derivative morpheme on any level.

**Proto-Athabaskan**. **q'$^w$as* is retained in all three branches.

**Eyak.** *=c$^h$īʔ*, the original meaning was apparently 'head' (q.v.), the Proto-Eyak term for 'neck' is unknown, cf. *=tə=q'əc'* 'collar' which looks similar to PA **q'$^w$as* 'neck'.

**Proto-Athabaskan-Eyak**. **q'$^w$as* (Athabaskan).

**Proto-Eskimo**. **uya-* (Fortescue et al. 2010: 420), scarcely retained with the meaning 'neck (non-anatomic, e.g., neck of bottle)'. In Proto-Yupik, the stem **uya-quʁ* 'neck' was derived (suffix **-quʁ* means 'smth. associated with or attached to smth.'). Cognate to the Aleut term. In Proto-Inuit, superseded with a derivative **quŋ-uciʁ* of unclear origin (Fortescue et al. 2010: 345).

**Proto-Aleut**. **uyu-χ* (Bergsland 1994: 457; Golovko 1994: 288), attested in all branches. Cognate to the Eskimo term.

**Proto-Chukotian**. **ləʁitən* (Fortescue 2005: 167), retained in all languages. Morphologically unclear. Can be cognate to the Itelmen term.

**Proto-Itelmen**. **xeyte-n* (Volodin, Khaloimova 2001: 222; Fortescue 2005: 167). Can be cognate to the Chukotian term.

**Proto-Nivkh**. **qor* (Fortescue 2016: 142; Savelyeva, Taksami 1965: 459; Savelyeva, Taksami 1970: 151).

**Proto-Samoyed**. **wayk-kə̆* (Janhunen 1977: 173), retained in all languages save Selkup. Replaced in Selkup by reflexes of Proto-Samoyed **soy* 'throat' (Janhunen 1977: 142). The word **wayk-kə̆* is etymologically a dual form of **wayk* 'shoulder' (< Proto-Uralic **wolka* 'shoulder'), itself preserved only in Selkup, being replaced in most other languages by **mə̆rkä* 'shoulder'.

**Proto-Yukaghir**. **n$^y$om-il* (Nikolaeva 2006: 307) is retained in both Yukaghir languages.

59. 'new'.

**Proto-Yeniseian**. **giʔ* (S. Starostin 1995: 227). Preserved in Ket-Yugh and Kott, not attested in Arin and Pumpokol.

**Proto-Athabaskan**. Not reconstructible, because 'new' is normally expressed with the help of various morphological structures based on roots with the meanings 'now', 'right now', 'recent' which looks like new formations.

**Eyak.** *q'a:-ya:*, a transparent new formation based on *q'ah* 'already, finally, now, by now'.

**Proto-Athabaskan-Eyak**. Not reconstructible.

**Proto-Eskimo**. **nuta-ʁ-aʁ* (Fortescue et al. 2010: 265), retained in all branches. Derived from the verb **nuta-ʁ-* 'to renew'.

**Proto-Aleut**. **taɣa-ða-l-* (Bergsland 1994: 380; Golovko 1994: 233), attested in Eastern and Atkan.

**Proto-Chukotian**. **tur-* (Fortescue 2005: 291), retained in all languages.

**Proto-Itelmen**. Not documented reliably or superseded with a Russian loan. Cf. **neʔn* 'now' from which an adjective 'new' is likely derived at least in one lect (Fortescue 2005: 380).

**Proto-Nivkh**. **ţur-* (Fortescue 2016: 36; Savelyeva, Taksami 1965: 243; Savelyeva, Taksami 1970: 454).

**Proto-Samoyed**. **nʸɒrpV* (Helimski 1997: 315) is retained in Enets and Mator. No other word for 'new' is reconstructible for Proto-Samoyed.

**Proto-Yukaghir**. Kolyma **inlʸə* (Nikolaeva 2006: 175–176) vs. Tundra **minčʸərpə-* (Nikolaeva 2006: 269).

60. 'night'.

**Proto-Yeniseian**. **siɢ* (S. Starostin 1995: 274). Preserved in all daughter languages. Word-final **-ɢ* is reconstructed primarily on the basis of its deletion in Ket-Yugh **si:* (the other uvular consonants are usually preserved).

**Proto-Athabaskan**. Two main candidates are **ƛ'eʔ* (Pacific Coast, Apachean) and **tʰec* (Northern).

**Eyak.** *χəƛ'* 'night / darkness', cf. the paronymous verb *=ɬ=χeʔƛ'* 'to get dark, night falls', cognate to PA **χeƛ'* / **χeɬ* 'to be dark'. Since the semantic shift '(to be) dark' > 'night' seems more normal than *vice verso*, it is likely the 'night' is the secondary meaning for Eyak; the same semantic derivation is found in some Athabaskan, e.g., Sarsi *xìɬ* 'night'.

**Proto-Athabaskan-Eyak**. **ƛ'eʔ, *tʰec* (Athabaskan).

**Proto-Eskimo**. **unnu-ɣ* (Fortescue et al. 2010: 407), retained in Yupik-Sirenik as 'night', having shifted in Inuit into the meaning 'evening', the original semantics is retained in Inuit verbal formations 'to become night', 'to spend the night', 'to work during the night'.

**Proto-Aleut**. **amax ~ *amɣi-χ ~* (Bergsland 1994: 62; Golovko 1994: 27, 233), attested in all branches. Tends to be superseded with **ðaya-χ* 'late evening' (Bergsland 1994: 160; Golovko 1994: 44).

**Proto-Chukotian**. **nəki-nək* (Fortescue 2005: 189), reduplicated stem, retained in all languages. Cognate to the Itelmen term.

**Proto-Itelmen**. **nki-nk* (Volodin 1976: 154; Fortescue 2005: 189). Reduplicated stem. Cognate to the Chukotian term.

**Proto-Nivkh**. **urk* (Fortescue 2016: 157; Savelyeva, Taksami 1965: 244; Savelyeva, Taksami 1970: 396).

**Proto-Samoyed**. **pi* (Janhunen 1977: 123) is retained in all Samoyed languages.

**Proto-Yukaghir**. Kolyma **em-il* (Nikolaeva 2006: 157–158) vs. Tundra **čiŋičə-l* (Nikolaeva 2006: 133).

61. 'nose'.

**Proto-Yeniseian**. **xaŋ* (S. Starostin 1995: 295). Preserved in Kott and Pumpokol; in Ket-Yugh, the old word for 'nose' was replaced with **ʔɔlɨn*, and in Arin, with *ar-quy*, where *-quy* = 'hole', as in *tim-quy* 'window'.

**Proto-Athabaskan**. **=nə=čʰĩ:xʸ < *=nə=čʰen-xʸ* (Krauss & Leer 1981: 115) is retained in all three branches. The stem is derived from PA *=čʰen* 'to smell' with the help of the auxiliary morpheme **nə* 'face' and the repetitive aspect suffix. Such a derivation should be relatively recent, since the nasalization is retained is some languages (old PAE **nC*-clusters normally lose the nasal element).

**Eyak.** *=ni:k'*, cognate to PA **=nə=ni:k'-* 'nostril', Tlingit *=ni:x'* 'to smell'.

**Proto-Athabaskan-Eyak**. **ni:k'* (Eyak).

**Proto-Eskimo**. **qəŋa-ʁ* (Fortescue et al. 2010: 325), retained in all branches.

**Proto-Aleut**. **anʁ-usi-* (Bergsland 1994: 76; Golovko 1994: 233), attested in all branches. Derived from **anʁ-* 'to breath'.

**Proto-Chukotian**. There are two candidates, intertwined between languages and dialects. First, **yeqa* (Fortescue 2005: 113), a basic term for 'human nose' in Chukchi and the Palana dialect of Alutor, meaning 'tip, end' in Koryak. Second, **qiŋ* (Fortescue 2005: 235), a basic term for 'nose (generic) / beak' in Koryak, Kerek and Alutor, meaning specifically 'nose (of animal)' in Chukchi. It is likely that the opposition **yeqa* 'nose (of human)' / **qiŋ* 'nose (of animal)' is to be reconstructed for Proto-Chukotian. In modern lects, **qiŋ* tends to acquire the generic meaning 'nose', superseding **yeqa*.

**Proto-Itelmen**. **qeqə-* (Volodin, Khaloimova 2001: 178; Fortescue 2005: 235).

**Proto-Nivkh**. **wiɣ* (Fortescue 2016: 163; Savelyeva, Taksami 1965: 244; Savelyeva, Taksami 1970: 55).

**Proto-Samoyed**. **pɨyɒ* ~ **puyɒ* (Janhunen 1977: 122–123) is retained in all daughter languages save Nganasan.

**Proto-Yukaghir**. **yoŋq-ul* (Nikolaeva 2006: 196) is retained in both Yukaghir languages.

62. 'not'.

**Proto-Yeniseian**. **wən* (S. Starostin 1995: 294 as **wə-*). Preserved in all daughter languages. Initial **w-* is reconstructed by S. Starostin on the basis of the voiced stop (or nasal *m*, assimilated from **b* under the influence of the following *n*) reflexation in all languages and dialects (Ket *bən^y*, etc.).

**Proto-Athabaskan**. The best candidate is the prefix **tuː-* attested as the main exponent of negation of assertion in the Pacific Coast and Apachean branches as well as in the Northern branch as a relic. Besides, there are attested very complex systems of negation in Northern lects. So we add four verbal morphemes that are likely to be reconstructed at least for the Proto-Northern level: the prefixes **iːʔ-* (perfective), **s-* (imperfective), the enclitic **=eː*, the proclitic **leʔ*.

**Eyak.** *ti-k' ...-q*, it is not entirely clear how the negative particle *tik'* is to be analyzed. The final -*k'* is apparently the morpheme *k'u* ~ *k'ə* — a negative prefix of interrogative pronouns. In this case the initial *ti-* resembles the Athab. negation **tuː* (< **tV-wV*?). Provisionally we accept this match and posit the Proto-Athabaskan-Eyak negative exponent **tV-*.

**Proto-Athabaskan-Eyak**. **tV-* (Athabaskan, Eyak).

**Proto-Eskimo**. **=nʁi-* (Fortescue et al. 2010: 460), retained as the suffix of negation of assertion in Yupik and Inuit.

**Proto-Aleut**. **=laka-* (Bergsland 1994: 518; Bergsland 1997: 84; Golovko 1994: 298), suffix used in the present tense. With other verbal forms, the enclitic **u=lax* is used (Bergsland 1994: 483; Bergsland 1997: 103; Golovko 1994: 293), originating from **a-/u-* 'to be' and the negative exponent **-lax*.

**Proto-Chukotian**. The basic negation of assertion is to be reconstructed as the verbal confix **ä-...-kä* (Fortescue 2005: 402), as it is attested in everywhere.

**Proto-Itelmen**. The basic negation of assertion is the particle **qaʔm* plus the verbal suffix **-aq* (Volodin 1976: 271–272; Fortescue 2005: 421).

**Proto-Nivkh**. According to Panfilov 1965: 158–159; Gruzdeva 1998: 44–45, the basic exponent of negation of assertion is the verb **qaw-* 'to be not' (Fortescue 2016: 140), which can function either as a copula or a verbal affix (Fortescue 2016: 171). Distinct from the prohibitive particle **ta* (Fortescue 2016: 144; Gruzdeva 1998: 34).

**Proto-Samoyed**. **e-* (Janhunen 1977: 26). Negative verb, inherited from Proto-Uralic and retained in all daughter languages.

**Proto-Yukaghir**. **əl* (Nikolaeva 2006: 155–156). Negative proclitic, retained in both Yukaghir languages.

63. 'one'.

**Proto-Yeniseian**. **qu-s-a* (S. Starostin 1995: 306). Preserved in all daughter languages. Forms such as Ket-Yugh **qɔʔ-k* 'one (animate)' clearly imply that **-s(a)* was a suffixal element in Proto-Yeniseian. Word-final **-a* is a morpheme common for most of Proto-Yeniseian numerals; as for the component *-*s*-, it may be compared with the singulative suffix *-*s* that is segmented out of archaic nominal stems such as 'eye' q.v. or 'stone' q.v.

**Proto-Athabaskan**. **łVq'*-, in many lects, the root was fused with a prefix or reanalyzed towards the shape *ł(V)*.

**Eyak.** *łĩhq-ih.*

**Proto-Athabaskan-Eyak**. **łVnq'*-, we follow Leer 2008a and reconstruct a medial nasal on account of vowel nasalization and aspiration in Eyak (Eyak *łĩhq* < **łĩhq'* according to Leer's rule).

**Proto-Eskimo**. **ataʁ-uciʁ* (Fortescue et al. 2010: 54), retained in all branches. Final element is probably the instrumental suffix *-*uciʁ*. Cognate to the Aleut form.

**Proto-Aleut**. **ataqa-* (Bergsland 1994: 106, 570; Golovko 1994: 236), attested in all branches. Cognate to the Eskimo form.

**Proto-Chukotian**. **ənnän* (Fortescue 2005: 345), retained in all languages. Morphologically unclear, probably derived from **ən-no* 'he, she, it' (Fortescue 2005: 342).

**Proto-Itelmen**. **nizəq* ~ **lizəq* (Fortescue 2005: 380), retained in Eastern and Southern Itelmen. In Western Itelmen, superseded with *qn-iŋ* '1', derived from **qun* 'once' (Volodin, Khaloimova 2001: 181; Fortescue 2005: 241).

**Proto-Nivkh**. **ɲə* (Fortescue 2016: 117, 178; Panfilov 1962: 181, 214–215; Gruzdeva 1998: 24).

**Proto-Samoyed**. **o-p* (Janhunen 1977: 28) is retained in all daughter languages except Selkup that has another derivative of the same root **o-*.

**Proto-Yukaghir**. **irk-* (Nikolaeva 2006: 177) is retained in Kolyma. Tundra **maːrqə-* (Nikolaeva 2006: 259) with its atypical long vowel in a closed syllable, according to Nikolaeva, can be a contraction of affirmative proclitic **mə-* + **irk-*.

64. 'person'.

**Proto-Yeniseian**. **keʔt* (S. Starostin 1995: 236). Preserved in all daughter languages. The word had a suppletive plural on the Proto-Yeniseian level, reconstructed as **ǯeʔŋ* (S. Starostin 1995: 309), probably the original plural of an unpreserved singular **ǯeʔ* 'person'.

**Proto-Athabaskan**. **te=neː* 'man / person' is the main candidate.

**Eyak.** *tə=χũh*, morphologically unclear.

**Proto-Athabaskan-Eyak**. **nVː* (Athabaskan).

**Proto-Eskimo**. **inʸu-ɣ* (Fortescue et al. 2010: 150), retained in all branches.

**Proto-Aleut**. **anʁ-aʁi-χ* ~ **anʁ-aʁi-na-χ* (Bergsland 1994: 74; Golovko 1994: 286), attested in all branches. Derived from **anʁ-aʁi-l-* 'to be alive' < **anʁ-* 'to breath'. Also **taya-ʁu-χ* 'man' (q.v.) can be used for generic 'person'.

**Proto-Chukotian**. **ʁuyä-mtä-wi-łʁə-n* (Fortescue 2005: 269), a complex stem apparently with descriptive semantics, although details are not entirely clear. It is reconstructed on the basis of the attested Koryak, Kerek and Alutor words for 'person'. The similar Chukchi term *ʔorawetlʔa-n* 'person' is probably a result of contamination with *ʔorawer* 'openly, visibly'. Can be cognate to the Itelmen term.

**Proto-Itelmen**. **kč'a-mzʸa-n-łx* (Volodin, Khaloimova 2001: 221; Ono 2003: 110; Fortescue 2005: 371). Morphological details are not entirely clear, it can be cognate to or at least calqued from the Chukotian term.

**Proto-Nivkh**. **ɲiɣ-vŋ* (Fortescue 2016: 115; Savelyeva, Taksami 1965: 452; Savelyeva, Taksami 1970: 210). It is likely that the Nivkh term for 'person' (self-designation of the Nivkhs) originally means 'the one living in a territory called *Niɣ*'.

**Proto-Samoyed**. **kɒə̆-sɒ* (Janhunen 1977: 61) retains the meaning 'person' in Mator and Kamass. In Northern Samoyed **kɒə̆-sɒ* changed its meaning to 'man'. The word goes back to Proto-Uralic **kali-sʸa* 'person' (preserved also in Mansi), derived from Proto-Uralic **kali-* (> Samoyed **kɒə̆-*) 'to die'.

**Proto-Yukaghir**. Kolyma **soromə* (Nikolaeva 2006: 415) vs. Tundra **köntə* (Nikolaeva 2006: 220).

65. 'rain'.

**Proto-Yeniseian**. **xur* (S. Starostin 1995: 297). Preserved in all daughter languages; however, there is a problematic relationship between the listed forms and the original Proto-Yeniseian word for 'water'. In Ket-Yugh, 'rain' is easily analyzable as a compound form: **ʔur* 'water' + **ʔes* 'sky'. However, Kott, Arin, and Pumpokol consistently feature *different* resonants in the root morphemes for 'water' and 'rain', e.g., Arin *kur* 'rain' vs. *kul* 'water', Pumpokol *ur-ait* (where *-ait* < **ʔes*) 'rain' vs. *ul* 'water'. This is accounted for in S. Starostin's reconstruction, where original **xur* 'rain' is opposed to **xurɹ* 'water'. It is not excluded that the two roots are, in the end, related (through some non-trivial morphophonological connection) on a higher level than Proto-Yeniseian, but for Proto-Yeniseian it is indeed preferable to separate them.

**Proto-Athabaskan**. **kʰʸa:n* is retained in all three branches.

**Eyak.** *k'u=leh*, literally 'something is happening', a transparent innovation.

**Proto-Athabaskan-Eyak**. **kʰa:n* (Athabaskan).

**Proto-Eskimo**. **cila-łuɣ* (Fortescue et al. 2010: 85), retained with the meaning 'rain' in Yupik and Inuit. Literally 'bad weather' with **cila* 'weather' and the suffix **-łuɣ* 'bad'.

**Proto-Aleut**. **kim-ðux* (Bergsland 1994: 239; Golovko 1994: 65, 203), attested in Atkan and Attuan; derived from **kim-s-* 'to descend, go down'. In Eastern and occasionally in Atkan, superseded with **čiχ-ta-l-* 'to rain', **čiχ-ta-χ* 'rain' (Bergsland 1994: 139), whose original Proto-Aleut meaning was 'to be wet'.

**Proto-Chukotian**. **muqä-* (Fortescue 2005: 179), retained as the basic root for '(to) rain' in Koryak and Kerek. In Chukchi, superseded with the root **ilə-* 'damp' (Fortescue 2005: 97).

**Proto-Itelmen**. **čuɣʷ* (Volodin, Khaloimova 2001: 158; Fortescue 2005: 383).

**Proto-Nivkh**. **ləɣ* ~ **nəɣ* (Fortescue 2016: 99; Savelyeva, Taksami 1965: 130; Savelyeva, Taksami 1970: 171).

**Proto-Samoyed**. **sɒr-ö* (Janhunen 1977 135–136), retained in Northern Samoyed and Mator, is derived from the verb **sɒrɒ-* 'to rain' (< Proto-Uralic **sʸaða-* 'to rain'). Kamass and Selkup have another derivative from the same verb.

**Proto-Yukaghir**. **tiwo* (Nikolaeva 2006: 440) is retained in both Yukaghir languages. Cf. also the root **liŋkə* (Nikolaeva 2006: 255) meaning 'rain' and 'to drink' in Chuvan and 'rain, water; to drink water' in Omok, but absent from modern Yukaghir languages.

66. 'red'.

**Proto-Yeniseian**. **sur-* (S. Starostin 1995: 278). Preserved everywhere, with the likely exception of Arin: Arin *tʸu:ra* cannot be regarded as a regular reflexation of Proto-Yeniseian **sur-* (the regular reflexation is found in Arin *sur* 'blood' q.v.).

**Proto-Athabaskan**. **=čʰi:xʸ* (Pacific Coast, Apachean) is the main candidate.

**Eyak.** *=č'e:ʔ.*

**Proto-Athabaskan-Eyak**. **=čʰi:x* (Athabaskan), **=č'e:ʔ* (Eyak).

**Proto-Eskimo**. **kavi-ʁ-* (Fortescue et al. 2010: 177), retained in all branches. In Inuit, tends to be superseded with the new formation **aðuɣ-valuɣ*, lit. 'blood-like' (Fortescue et al. 2010: 6).

**Proto-Aleut**. **ulu:-ða-* (Bergsland 1994: 436; Golovko 1994: 134, 220), attested in all branches. Derived from **ulu-* 'meat' (q.v.).

**Proto-Chukotian**. **yərrə-* (Fortescue 2005: 123), retained in all languages except for Chukchi. Can be cognate to the Itelmen term.

**Proto-Itelmen**. **č'a-* (Volodin, Khaloimova 2001: 164; Volodin 1976: 320; Fortescue 2005: 123). Can be cognate to the Chukotian term.

**Proto-Nivkh**. **baɣ-la-* (Fortescue 2016: 19; Savelyeva, Taksami 1965: 190; Savelyeva, Taksami 1970: 250).

**Proto-Samoyed**. **nʸar-* (Janhunen 1977: 107–108) is retained in Northern Samoyed, Mator and Selkup.

**Proto-Yukaghir**. Kolyma **keylə-nʸ-*, apparently the same root as 'dry' (Nikolaeva 2006: 204) vs. Tundra **nʸamučə-nʸ-* (Nikolaeva 2006: 287).

67. 'road'.

**Proto-Yeniseian**. It seems that Ket-Yugh preserved the original lexical distinction between *qɨq* 'summer road' (S. Starostin 1995: 301) and **qoʔt* 'winter road' (S. Starostin 1995: 261), whereas Kott, Arin, and Pumpokol generalized one word of the two. We have to include both terms into comparison as synonyms.

**Proto-Athabaskan**. **tʰeɲ(e)* is retained in all three branches.

**Eyak.** *tʰa:*, cognate to Athabaskan with **ɲ > 0* or simply with the loss of nasalization (vowel correspondence between Eyak *tʰa:* and **tʰeɲ(e)* is unclear).

**Proto-Athabaskan-Eyak**. **tʰeɲ* (Athabaskan, Eyak).

**Proto-Eskimo**. **apʁ-un* (Fortescue et al. 2010: 41), retained in Yupik and Inuit. The final element is the instrumental suffix **-un*. In some lects, superseded with **tumə* 'track, footprint' (Fortescue et al. 2010: 381).

**Proto-Aleut**. **aka-lu-χ* (Bergsland 1994: 43; Golovko 1994: 204), attested in all branches.

**Proto-Chukotian**. **rəʁet* (Fortescue 2005: 258), retained in all languages except for Alutor. Tends to be superseded with **winvə* 'track' (Fortescue 2005: 329).

**Proto-Itelmen**. **ktχʷazʸ* ~ **ktχʷaɬ* (Volodin, Khaloimova 2001: 150; Fortescue 2005: 258).

**Proto-Nivkh**. **ɖiv* (Fortescue 2016: 53; Savelyeva, Taksami 1965: 132; Savelyeva, Taksami 1970: 354).

**Proto-Samoyed**. **ätɒ* (Janhunen 1977: 24) is retained in Mator, Kamass and Selkup. The root also has derivatives in Nenets.

**Proto-Yukaghir**. **yaw-ul* (Nikolaeva 2006: 186), attested in Tundra Yukaghir. Kolyma has the root **čuɣö* (Nikolaeva 2006: 144), apparently borrowed from Tungusic **ǯuku-* 'road'.

68. 'root'.

**Proto-Yeniseian**. **ci:ǯ* (S. Starostin 1995: 217). Best preserved in Ket-Yugh and in Pumpokol. In Kott-Arin, the original simple form was replaced with **tem-bul*, where **-bul* may be the same root as 'foot', q.v. and the first part is technically etymologizable as an assimilated form of the original **či:ǯ* (cf. the dialectal Arin form *tɨy-bul* ~ *tuy-bul*).

**Proto-Athabaskan**. The main candidates are **χat* (Pacific Coast) and **=qe:c'* (Northern). The latter one, **=qe:c'*, is supported by the Eyak cognate *qe:c'* 'root'. As for **χat*, if such forms as Ahtena *=ʁahteʔ* 'main root of tree', Tanaina *=ʁeχta* 'curved root or branch used for ribs on boat', Tututni *=ɣahteʔ* 'root of tree' are related, they imply an original complex morphological structure contracted into *=ʁat* is some Pacific Coast languages. Can Tlingit *χa:t* 'root' be an old loan from Pacific Coast?

**Eyak.** *qe:c'*.

**Proto-Athabaskan-Eyak**. **=qe:c'* (Athabaskan, Eyak).

**Proto-Eskimo**. **aku-ʁ* (Fortescue et al. 2010: 15), retained in all branches.

**Proto-Aleut**. **halɣi-χ* (Bergsland 1994: 52; Golovko 1994: 219), attested in Atkan and Attuan. Superseded with *quŋ-lux 'root' (Bergsland 1994: 338) in Eastern.

**Proto-Chukotian**. There are two main candidates. First, **tätqu* (Fortescue 2005: 282), which is the basic term in Koryak (Zhukova 1990: 77) and Alutor (Kibrik et al. 2004: 533), meaning 'cambium' in Chukchi. Second, **kinmä* (Fortescue 2005: 138), the basic term in Chukchi, which is also attested in other languages with a 'root' semantics. We treat both as synonyms. Cf. also **nən(n)əl*, which denotes 'root (generic)' in a dialect of Chukchi (Moll, Inenlikei 1957: 88) and 'lobe of root' in Koryak and Alutor (Fortescue 2005: 191).

**Proto-Itelmen**. **pŋil* (Volodin, Khaloimova 2001: 163; Fortescue 2005: 385).

**Proto-Nivkh**. There are two candidates, both are attested in all languages with the gloss 'root'. First, **mir-ləɣ* ~ **vir-ləɣ* (Fortescue 2016: 105; Savelyeva, Taksami 1965: 187; Savelyeva, Taksami 1970: 54). Second, **oz* (Fortescue 2016: 131; Savelyeva, Taksami 1970: 247). It is unclear how the semantic difference between the two is to be reconstructed ('root of plant' vs. 'root of tree' or 'root' vs. 'base of trunk'?), we treat them as synonyms.

**Proto-Samoyed**. **wɒncə̹* (Janhunen 1977: 171), retained in all Samoyed languages, goes back to Proto-Uralic **wancə̹w* 'root'.

**Proto-Yukaghir**. Kolyma **larq-ul* (Nikolaeva 2006: 235). Tundra Yukaghir has *warulu:* 'root'. Nikolaeva compares this word with Kolyma *ožu:* 'thin root used as a thread for fastening boats' and reconstructs Proto-Yukaghir **wonč-*. The Proto-Yukaghir form is usually compared with Proto-Uralic **wancə̹w* 'root'. While the comparison of Tundra *warulu:* with Kolyma *ožu:* is acceptable phonologically, it faces a morphological problem, since there is no denominal suffix *-lu:* in Yukaghir. We suggest an alternative etymology: Tundra *warulu:* is derived with the deverbal noun suffix *-u:* from the verbal stem *warul-*, attested in the Tundra derivatives *warul-mu-* 'to become strong (of rope, thread)' and *warul-we-* 'id.'. The stem *warul-* itself is derived from the verb *war-* 'to be strong', related to Kolyma *ad-* 'firm, strong' < Proto-Yukaghir **wað-* (Nikolaeva 2006: 449-450). This derivation makes sense, because willow roots were used by the Tundra Yukaghir to tie together posts for Yukaghir traditional tents. It seems that the word *warulu:* originally referred to a specific kind of root used for tying, and only later became the basic word for any root.

69. 'round'.

**Proto-Yeniseian**. Poorly documented item, sometimes superseded with loans. Formally the best candidate is virtual **pVd-* reconstructed on the basis of the Kott form.

**Proto-Athabaskan**. Is it very likely that the actual Proto-Athabaskan system was binary with the opposition 'round 3D' / 'round 2D'. The unquestionable candidate is **=wa:nc'* / **=wa:ns*, retained in all three branches (the retained nasal component should point to a relatively recent PA suffixation **=wa:n-c'* / **=wa:n-s*). The second candidate, also attested in all three branches, is **če=ʁul* (in Krauss & Leer 1981: 108, 204, treated as a bisyllabic root **=čəʁʷəƛ'* / **=čəʁʷəl*). The exact meaning of initial **če-* is unclear, but this element is detachable and functions as a verbal prefix in the Pacific Coast lects (Hupa). In the Northern and Apachean groups, the sequence **če=ʁul* was fossilized and contracted.

**Eyak.** *qəmək'*, morphologically unclear, somewhat resembles PA **=wã:c'*, but details remain vague.

**Proto-Athabaskan-Eyak**. **=wVn-*, **=χVl* (Athabaskan).

**Proto-Eskimo**. Not reconstructible. In Yupik, 'round' is usually expressed with various suffixed derivatives from **akða-ɣ-* 'to roll' (Fortescue et al. 2010: 11). In Inuit, 'round' is usually expressed with various suffixed derivatives from **aŋva-luʁ-* (Fortescue et al. 2010: 39), whose literal proto-meaning is expected to be 'place of opening', i.e. 'round hole'(?).

**Proto-Aleut**. Not reconstructible with certainty. Cf. Atkan *imu-ðiɣa-* 'round' (Bergsland 1994: 198; Golovko 1994: 220) < **imu-χ* 'circle; area around' and Eastern *qim̥a-ðɣu-l-* 'round' (Bergsland 1994: 324; Bergsland & Dirks 1978: 153) < **qim̥a-* '?' + *-ðɣu-* 'to become, to get to'. Both looks like new formations.

**Proto-Chukotian**. Basic expressions for 'round' are based on the verbal root **kəvlə-* 'to roll' (Fortescue 2005: 156) at least in Chukchi and Alutor, but it may be a parallel development.

**Proto-Itelmen**. Not documented reliably or superseded with a Russian loan.

**Proto-Nivkh**. **bulk-bulk-* (Fortescue 2016: 27; Savelyeva, Taksami 1965: 192; Savelyeva, Taksami 1970: 275). The substantive **bulk* means 'small ball, skein', synchronic expressions for ‘(to be) round’ are derived from it via reduplication (or with a causative suffix as in Amur).

**Proto-Samoyed**. Not reconstructible for Proto-Samoyed.

**Proto-Yukaghir**. **pöm-nə-* (Nikolaeva 2006: 347–348), retained in both Yukaghir languages. Comparison with isolated Even forms meaning ‘to wind, be twisted’ and ‘loop on a thread, rope’, proposed in (Nikolaeva 2006: 348), is not convincing.

**Proto-Burushaski**. *pinḏoro ~ biḏiro* is borrowed from Dardic (cf. Khowar *pinḏoru*, Shina *biḏiru* ‘round’).

70. ‘sand’.

**Proto-Yeniseian**. **pən-əŋ* (S. Starostin 1995: 248). Preserved everywhere except in Kott (where the meaning shifted to ‘ashes’).

**Proto-Athabaskan**. **saːx* is retained in all three branches.

**Eyak.** *čʰiːš-k*.

**Proto-Athabaskan-Eyak**. **saːx* (Athabaskan), **čʰiːš* (Eyak).

**Proto-Eskimo**. **qavə-yaʁ* (Fortescue et al. 2010: 318), retained in all branches. The starting root is unclear, the suffix **-yaʁ* means ‘place where action takes place’. In some Inuit lects, superseded with the complex stem **ciɣu-ʁ-aʁ ~ *ciʁu-ʁ-aʁ* (Fortescue et al. 2010: 94), which resembles Aleut **čuɣu-χ* ‘sand’.

**Proto-Aleut**. **čuɣu-χ* (Bergsland 1994: 151; Golovko 1994: 243), attested in all branches.

**Proto-Chukotian**. **čəɣäy* (Fortescue 2005: 50), retained in all languages, for Chukchi see Moll, Inenlikei 1957: 145.

**Proto-Itelmen**. **tosʸx* (Volodin, Khaloimova 2001: 186; Fortescue 2005: 387).

**Proto-Nivkh**. **maʁ ~ *maɣ* (Fortescue 2016: 102; Savelyeva, Taksami 1965: 292; Savelyeva, Taksami 1970: 178), attested in Amur and North Sakhalin. In East and South Sakhalin, superseded with **ɢom-r* (Fortescue 2016: 141; Savelyeva, Taksami 1970: 178), which is probably derived from the verb **ɢom-* ‘to dwell near sea shore (e.g. in summer settlements)’ (Savelyeva, Taksami 1970: 144) with the common deverbal suffix **-r*.

**Proto-Samoyed**. **yɒə̆* (Janhunen 1977: 36–37), the root for ‘earth (soil)’, also means ‘sand’ in Enets and Nganasan; its derivative **yɒə̆-rɒ* ‘sandy / sandy bank’ is reflected in all Samoyed languages. Another word for ‘sand’, **puə̆rɒ* (Helimski 1997: 251), attested only in Mator and Kamass, is more likely an areal isogloss than retention from Proto-Samoyed.

**Proto-Yukaghir**. **noŋqə* (Nikolaeva 2006: 309) has the meaning ‘sand’ in Kolyma Yukaghir, its Tundra cognate means ‘ashes’, q.v. The Tundra word for ‘sand’, going back to **öninčʸə* (Nikolaeva 2006: 331), apparently is a loan from Tungusic.

71. ‘to say’.

**Proto-Yeniseian**. **saga-* (S. Starostin 1995: 269). Preserved in all the languages where it is attested, but the original semantics raises doubts: it is possible that the actual meaning of Proto-Yeniseian **saga-* was closer to ‘speak, talk’ than to ‘say’, considering that in Ket-Yugh at least, the highly irregular verb **=ma* ‘to say’ looks more archaic than **saga-*; formally, however, it is difficult to project **=ma* onto the Proto-Yeniseian level due to its conspicuous absence from old records of Kott material.

**Proto-Athabaskan**. **=niː* is retained in all three branches.

**Eyak.** *=le*, cognate to Athabaskan.

**Proto-Athabaskan-Eyak**. **=niː*.

**Proto-Eskimo**. **pi-* (Fortescue et al. 2010: 282), retained in all branches with polysemy: ‘to do, act / to say’. Cognate to the Aleut stem.

**Proto-Aleut**. **hi-l- ~ *hi-χta-l-* (Bergsland 1994: 167, 168; Golovko 1994: 165, 166, 266), attested in all braches. Cognate to the Eskimo stem.

**Proto-Chukotian**. **iv-* (Fortescue 2005: 105), retained in all languages. Distinct from **təv-* ‘to tell’ (Fortescue 2005: 304).

**Proto-Itelmen**. **χine-* (Volodin, Khaloimova 2001: 205; Volodin 1976: 33, 149, 245; Fortescue 2005: 262), a Western verb. Distinct from **ɬo-* 'to tell, speak' (Volodin, Khaloimova 2001: 52; Fortescue 2005: 304).

**Proto-Nivkh**. **it-* (Fortescue 2016: 80; Savelyeva, Taksami 1965: 387; Savelyeva, Taksami 1970: 99). Distinct from **pur-* 'to tell' (Fortescue 2016: 138; Savelyeva, Taksami 1970: 406).

**Proto-Samoyed**. **mɒ- ~ *mɒn-* (Janhunen 1977: 88), retained in Northern Samoyed and Kamass, goes back to Proto-Uralic **moni-* 'to say' (reflected in Mari and Hungarian).

**Proto-Yukaghir**. **mon-* (Nikolaeva 2006: 274), retained in both Yukaghir languages, is cognate to, or borrowed from, Proto-Samoyed **mɒn-* 'to say'.

72. 'to see'.

**Proto-Yeniseian**. **t=...=ɔŋ* (S. Starostin 1995: 290). Preserved in Ket-Yugh and Kott, but apparently lost in Arin and Pumpokol. Ket-Yugh and Kott agree on the basic structure of the verb, consisting of the directional prefix **t=* and the root **=ɔŋ*, separated by grammatical morphemes such as the tense and conjugation markers.

**Proto-Athabaskan**. **=ʔeːn* is retained in all three branches.

**Eyak.** *=ʔe ~ =ʔã*, cognate to Athabaskan.

**Proto-Athabaskan-Eyak**. **=ʔVːn* (Athabaskan, Eyak).

**Proto-Eskimo**. An unstable item which cannot be reconstructed with certainty.

**Proto-Aleut**. **uku-l- ~ *uku-χta-l-* (Bergsland 1994: 429, 430; Golovko 1994: 133, 190), attested in all branches, polysemy: 'to see / to look'.

**Proto-Chukotian**. **ləʁu-* (Fortescue 2005: 167), retained in all languages. Can be cognate to the Itelmen term.

**Proto-Itelmen**. **əɬčku-* (Volodin, Khaloimova 2001: 140; Fortescue 2005: 376). Can be cognate to the Chukotian term.

**Proto-Nivkh**. **nətə* (Fortescue 2016: 112; Savelyeva, Taksami 1965: 84; Savelyeva, Taksami 1970: 94). Distinct from **ama-* 'to look, watch' (Fortescue 2016: 12; Savelyeva, Taksami 1965: 395; Savelyeva, Taksami 1970: 498).

**Proto-Samoyed**. **mə̆ncV̦-* (Janhunen 1977: 86–87) is retained in Nenets, Enets, Kamass and Selkup.

**Proto-Yukaghir**. **yöː-* (Nikolaeva 2006: 191) is retained in both Yukaghir languages, although the Tundra reflex of this root is not the main word for 'to see' in that language. The main Tundra word for 'to see', *ičuo-* (Nikolaeva 2006: 460), is borrowed from Tungusic **iče-* 'to see'.

73. 'seed'.

**Proto-Yeniseian**. Not reconstructible due to lack of data.

**Proto-Athabaskan**. Not reconstructible reliably.

**Eyak.** Not documented.

**Proto-Athabaskan-Eyak**. Not reconstructible.

**Proto-Eskimo**. Not reconstructible.

**Proto-Aleut**. Not reconstructible.

**Proto-Chukotian**. Not reconstructible.

**Proto-Itelmen**. Not reconstructible.

**Proto-Nivkh**. Not reconstructible.

**Proto-Samoyed**. Not reconstructible reliably. Nenets and Enets use reflexes of **sə̆ymä* 'eye' (Janhunen 1977: 132); in most other languages the word for 'seed' is not attested.

**Proto-Yukaghir**. Not reconstructible.

74. 'to sit'.

**Proto-Yeniseian**. **xu-* (S. Starostin 1995: 297). Preserved in Kott-Arin and still seen in the Ket-Yugh infinitive form *u-ŋ* 'to sit'; replaced in Ket-Yugh with **ses-* otherwise.

**Proto-Athabaskan**. *=*ta:* is retained in all three branches.

**Eyak.** =*ta*, cognate to Athabaskan.

**Proto-Athabaskan-Eyak**. *=*ta:* (Athabaskan, Eyak).

**Proto-Eskimo**. **aqu-mə-ɣa-* ~ **aqu-vət-* (Fortescue et al. 2010: 44), retained in Yupik (**aqu-mə-ɣa-*) and Inuit (**aqu-vət-*).

**Proto-Aleut**. **uŋu-či-l-* (Bergsland 1994: 448; Golovko 1994: 266), attested in all branches.

**Proto-Chukotian**. There are two candidates. First, **vakəʁo-* and the resultative **vakəʁo-tva-* (Fortescue 2005: 312), which mean 'to sit down' and 'to sit' respectively in Chukchi and Kerek; in Alutor, the deverbative 'seat' is attested. The stem **vakəʁo-* is morphologically unclear. Second, **təva-ɣal-* (Fortescue 2005: 304), meaning 'to sit down' in Koryak and Alutor, from which 'to sit' is derived in Koryak (resultative *va-ɣalə-tva-*), whereas in Alutor *tva-lʸʔat-* 'to sit; to be located' the habitual suffix is used. The verb **təva-* itself means 'to be (somewhere), live, exist' everywhere in Chukotian (Fortescue 2005: 304). The problem is that **təva-* and its derivative **təva-ɣal-* 'to sit down' are likely to be cognate to Itelmen **ɬa-* 'to sit / to live', **ɬa-wul-* 'to sit down' (q.v.). Various scenarios of semantic development can be proposed. We prefer to posit both **vakəʁo-tva-* and **təva-* as technical synonyms for 'to sit'.

**Proto-Itelmen**. **ɬa-* (Volodin, Khaloimova 2001: 205; Fortescue 2005: 304). The same root in **ɬa-wul-* 'to sit down' (Volodin, Khaloimova 2001: 202; Fortescue 2005: 304).

**Proto-Nivkh**. **tiv-* (Fortescue 2016: 148; Savelyeva, Taksami 1965: 386; Savelyeva, Taksami 1970: 98).

**Proto-Samoyed**. **ɒmtǝ̆-* (Janhunen 1977: 17–18) is retained in all Samoyed languages.

**Proto-Yukaghir**. Kolyma **montə-* (Nikolaeva 2006: 276) vs. Tundra **saɣanə-* ~ **saŋqənə-* (Nikolaeva 2006: 393).

75. 'skin (human)'.

**Proto-Yeniseian**. Not reconstructible due to lack of data. The Ket-Yugh word (Ket *î:*, Yugh *iɔl* ~ *iɣɔl* ~ *igɔl*) is comparable with Kott *e:k* 'hair', meaning that the original meaning of the etymon was probably closer to 'body hair; animal hair, fur' than to 'skin'.

**Proto-Athabaskan**. Not reconstructible reliably.

**Eyak.** =*tʰah.*

**Proto-Athabaskan-Eyak**. *=*tʰa(:)n* (Eyak). We follow Leer 2008a and reconstruct a medial nasal on account of vowel nasalization and aspiration in Eyak.

**Proto-Eskimo**. **uvinə-ɣ* (Fortescue et al. 2010: 419), meaning '(human) skin' in Inuit and '(human) skin' or '(human) body' in Yupik. Distinct from **ami-ʁ* (Fortescue et al. 2010: 25), which means 'hide, animal skin' in both Yupik and Inuit.

**Proto-Aleut**. **qačχi-χ* (Bergsland 1994: 292; Golovko 1994: 78, 218), attested in all branches.

**Proto-Chukotian**. **kəlɣə* (Fortescue 2005: 145), retained in all languages. Cognate to the Itelmen term.

**Proto-Itelmen**. **kilɣi-* (Volodin, Khaloimova 2001: 162; Fortescue 2005: 145), meaning 'skin (of human) / body'. Cognate to the Chukotian term.

**Proto-Nivkh**. **hal* (Fortescue 2016: 70; Savelyeva, Taksami 1965: 182; Savelyeva, Taksami 1970: 422), polysemy 'skin (of human) / body'. Distinct from **ŋaɣ-r* 'skin (of animal)' (Fortescue 2016: 118; Savelyeva, Taksami 1965: 182; Savelyeva, Taksami 1970: 233).

**Proto-Samoyed**. **kopɒ* (Janhunen 1977: 73), retained in all Samoyed languages, goes back to Proto-Uralic **kopa* 'skin'.

**Proto-Yukaghir**. Kolyma **qa:r* (Nikolaeva 2006: 379) vs. Tundra **sawa* (Nikolaeva 2006: 399). Nikolaeva compares the Kolyma word to Tundra *qayr* 'skin from the head of an animal', but the comparison is phonologically irregular. In both Yukaghir languages the word for 'skin' is used also for 'bark' ("skin of tree") and 'cloud' ("skin of sky").

76. ‘to sleep’.

**Proto-Yeniseian**. **=qɔt*, see notes on ‘to lie’; in Proto-Yeniseian, the meanings ‘lie’ and ‘sleep’ were most likely expressed by the same root.

**Proto-Athabaskan**. An unstable item. A possible candidate is **=laːɬ* attested in Pacific Coast and as a relic in Northern.

**Eyak.** *=cʰuʔt.*

**Proto-Athabaskan-Eyak**. **=ɬaːɬ* (Athabaskan), **=cʰVt* (Eyak). Leer (2010: 179) compares Eyak *=cʰuʔt* with Tlingit *=kʰiːt* ‘to snore’, if so the PAET form should be **=kʸʰVt*.

**Proto-Eskimo**. **qava-ʁ-* (Fortescue et al. 2010: 317) means ‘to sleep’ in Yupik. In Inuit it is either retained with the semantics of sleeping as shaman’s words or narrowed into the meaning ‘sleep on back in water (of seal)’. The Proto-Inuit term is **činə-ɣ-* ‘to sleep’ (Fortescue et al. 2010: 87) without reliable Yupik cognates.

**Proto-Aleut**. **saʁa-l-* (Bergsland 1994: 345; Golovko 1994: 271), attested in all branches.

**Proto-Chukotian**. **yəlq-ät-* (Fortescue 2005: 120), retained in all languages except for Kerek. If the common verbalizer **-ät-* is to be singled out, the root **yəlq-* is expected to be nominal or adjectival.

**Proto-Itelmen**. **ŋʷikɬ- ~ *ŋʷiksi-* (Volodin, Khaloimova 2001: 208; Fortescue 2005: 390).

**Proto-Nivkh**. **qo-* (Fortescue 2016: 141; Savelyeva, Taksami 1965: 403; Savelyeva, Taksami 1970: 150).

**Proto-Samoyed**. **kont-ö-* (Janhunen 1977: 73) is retained in all daughter languages. Similarity to Proto-Yukaghir **qont-oː-* ‘to lie’ may be due to borrowing in either direction.

**Proto-Yukaghir**. **yoŋčə-* (Nikolaeva 2006: 194–195) is retained in both Yukaghir languages, although the Tundra reflex of this root is not the main word for ‘to sleep’ in that language. The root of the main Tundra word for ‘to sleep’, *aː-we-* (Nikolaeva 2006: 115), is borrowed from North Tungusic **aː-* ‘to sleep’.

77. ‘small’.

**Proto-Yeniseian**. **pəɲ-* (S. Starostin 1995: 248). Preserved only in Ket-Yugh. Kott, Arin, and Pumpokol parallels are complex forms of either clearly secondary or etymologically obscure origins.

**Proto-Athabaskan**. An unstable item, not reconstructible reliably.

**Eyak.** *kʰuč’-k.*

**Proto-Athabaskan-Eyak**. **kʰVč’* (Eyak).

**Proto-Eskimo**. **mikə-* (Fortescue et al. 2010: 219), retained in all branches.

**Proto-Aleut**. **čuqu-ða-l-* (Bergsland 1994: 156; Golovko 1994: 151, 224), attested as generic ‘(to be) small’ in Atkan and Attuan, but shifted into the specific meaning ‘to be extremely small’ in Eastern, having been superseded with the new formation *aŋuna-laka-n ~ aŋunaː-ð(a)-laka-n*, lit. ‘not big’ (Bergsland 1994: 91; Bergsland & Dirks 1978: 90).

**Proto-Chukotian**. **əppəlu* (Fortescue 2005: 347), retained in all languages except for Alutor (Alutor ‘small’ < **məq-*, Fortescue 2005: 184). Morphologically unclear, if a compound, the second part can be cognate to the Alutor term. Distinct from the specific term **məl-* ‘small (мелкий), minute, fine’ (Fortescue 2005: 181).

**Proto-Itelmen**. **ulɣu-* (Volodin, Khaloimova 2001: 169; Fortescue 2005: 376).

**Proto-Nivkh**. **mat-k-* (Fortescue 2016: 100; Savelyeva, Taksami 1965: 204; Savelyeva, Taksami 1970: 177). The original root **mat-* ‘small’ is retained in derivatives.

**Proto-Samoyed**. **üćä̮* (Janhunen 1977: 31) is retained in Northern Samoyed and Selkup.

**Proto-Yukaghir**. **lʸuk-* (Nikolaeva 2006: 252–253) is retained in both Yukaghir languages.

78. ‘smoke’.

**Proto-Yeniseian**. **duʔ* (S. Starostin 1995: 224). Preserved in all daughter languages. Evidence for a final back consonant is weak and inconclusive.

**Proto-Athabaskan**. **ɬet* is retained in all three branches.

**Eyak.** *ɬãht*, cognate to Athabaskan.

**Proto-Athabaskan-Eyak**. **lVnt* (Athabaskan, Eyak), we follow Leer 2008a and reconstruct a medial nasal on account of vowel nasalization and aspiration in Eyak.

**Proto-Eskimo**. **puyu-ʁ* (Fortescue et al. 2010: 296), retained in all branches.

**Proto-Aleut**. **ʍaχ* (Bergsland 1994: 460; Golovko 1994: 170, 205), attested in all branches as the neutral term for 'smoke'. Distinct from **huyu-χ* 'white smoke; steam, vapor' (Bergsland 1994: 457; Golovko 1994: 168), which is cognate to Eskimo **puyu-ʁ* 'smoke'.

**Proto-Chukotian**. The lexical opposition 'visible smoke' / 'invisible smoke, smoke in house, fumes' is characteristic of the Chukotian-Itelmen area. The situation is rather tangled with three candidates competing with each other. One of the possible scenarios is to reconstruct the reduplicated stem **ipi-ʔipi* (Fortescue 2005: 103) with the meaning 'visible smoke', this is the Alutor term for 'visible smoke' (glossed simply 'smoke' in Kibrik et al. 2004: 398) and one of the two Koryak words for 'smoke' — *ipi-ip* apparently with the specific meaning 'visible smoke' (glossed as 'smoke; vapor' in Zhukova 1990: 35). The second stem is reduplicated **ŋəl-ŋəl* (Fortescue 2005: 201) 'invisible smoke', which is attested as the only term for 'smoke' in Chukchi (Moll, Inenlikei 1957: 93) and apparently Kerek (the exact Kerek semantics is unknown); Koryak *ŋəl-ŋəl* is glossed as 'smoke (in house)' in Zhukova 1967: 116 and simply as 'smoke' in Zhukova 1990: 67. The third term is **təqi-* (Fortescue 2005: 300), attested only is Alutor apparently with the meaning 'invisible smoke' (Kibrik et al. 2004: 557).

**Proto-Itelmen**. **t'i-* (Volodin, Khaloimova 2001: 92, 151; Fortescue 2005: 301), meaning 'visible smoke / vapor / to smoke'. Can be cognate to Chukotian **təqi-* 'a k. of smoke'. Distinct from **ŋačəz* 'invisible smoke' (Volodin, Khaloimova 2001: 151; Fortescue 2005: 201), which may ultimately be cognate to Chukotian **ŋəl-* 'invisible smoke'.

**Proto-Nivkh**. There are two candidates for 'smoke' intertwined between the lects. First, **taw-laŋ* (Fortescue 2016: 147), derived from the verb **taw-* 'to smoke' (Savelyeva, Taksami 1970: 380). Second, **tuv* (Fortescue 2016: 152–153; Savelyeva, Taksami 1965: 136; Savelyeva, Taksami 1970: 387), derived from **tuv-* 'to burn (tr.)' q.v. We treat them as synonyms.

**Proto-Samoyed**. **küntə̆* (Janhunen 1977: 79), the main word for 'smoke' in Enets, Nganasan and Mator, goes back to Proto-Uralic **künti*, whose Finno-Ugric reflexes mean 'fog'. Cf. **kə̆čku* (Janhunen 1977: 40) with reflexes meaning 'fog' (Enets, Nganasan) and 'smoke' (Selkup).

**Proto-Yukaghir**. Kolyma **lʸuː-l* (Nikolaeva 2006: 251) vs. Tundra **köyrinčʸə* (Nikolaeva 2006: 216).

79. 'to stand'.

**Proto-Yeniseian**. **=diŋ* ~ **=dik* (S. Starostin 1995: 221). Preserved everywhere except for Ket-Yugh; the basic equivalent for 'stand' in Ket-Yugh (**ʔipɨn*) finds no parallels in Kott, Arin, and Pumpokol, and so, technically, counts as a replacement, although from an unknown source.

**Proto-Athabaskan**. **=heːn* is retained in all three branches.

**Eyak.** *=ãːʔ*, cognate to Athabaskan.

**Proto-Athabaskan-Eyak**. **=hVːn* (Athabaskan, Eyak).

**Proto-Eskimo**. An unstable item, the best candidate is **naŋə-ʁ-* (Fortescue et al. 2010: 235) which is fragmentarily retained in both branches meaning 'to stand / to stand up' in Yupik and stative 'to stand' in Inuit. The obvious candidate for the Proto-Eskimo meaning 'to stand up' is **nʸəkəvə-* (Fortescue et al. 2010: 246) which retains the change-of-state semantics in the majority of the Yupik and Inuit lects. The derivative Yupik-Inuit stem **nʸəkəv-ʁ-a-* (Fortescue et al. 2010: 246) means generally 'to stand' in some Yupik lects and specifically 'to stand upright, be upright' in Inuit (apparently innovations according to the productive morphological model).

**Proto-Aleut**. There are two verbs with the meaning ‘to stand, be in upright position (of human)’, both are widely attested and coexist within dialects. The first one is **anqa-χta-l-* (Bergsland 1994: 78; Golovko 1994: 31, 272) from **anqa-l-* ‘to stand up’. The second one is **haχ-ta-l-* (Bergsland 1994: 33; Golovko 1994: 162) from the bound root **haʁ-* ‘to rise from lying position *vel sim.*’. The semantic and pragmatic difference is unclear.

**Proto-Chukotian**. **təvella-* (Fortescue 2005: 315), morphologically unclear, retained in all languages, usually means ‘to stand up’, although the meaning ‘to stand’ is also attested; the derived stem with the resultative suffix **-tva-* means ‘to stand’.

**Proto-Itelmen**. **tχ-zo- ~ *t-zo-* (Volodin, Khaloimova 2001: 210; Volodin 1976: 210, 258; Ono 2003: 30; Fortescue 2005: 242). Fluctuation *tχ ~ t* is unclear, either simplification of heavy clusters or two different root, which synchronously contaminate. Final **-zo-* is a continuative suffix (Fortescue 2005: 422).

**Proto-Nivkh**. **gəpr-* (Fortescue 2016: 64; Savelyeva, Taksami 1965: 411; Savelyeva, Taksami 1970: 127). Morphologically unclear.

**Proto-Samoyed**. **nu- ~ *nɨ-* (Janhunen 1977: 104) is retained in Nenets, Enets, Kamass and Selkup.

**Proto-Yukaghir**. **oŋq-oː-* (Nikolaeva 2006: 331–332), retained in both Yukaghir languages, is derived from the root **öŋkə-*.

80. ‘star’.

**Proto-Yeniseian**. **qɔːqa* (S. Starostin 1995: 265). Preserved in all daughter languages. In Kott and Arin, the word shows fusion with the same obscure prefix as in the word for ‘dog’ q.v. (**al=qɔːqa ~ *il=qɔːqa*).

**Proto-Athabaskan**. **seŋʔ ~ *cʰeŋʔ* is retained in all three branches, although phonetic reflexes are very complex (Krauss & Leer 1981: 65-68). It is unclear whether we should introduce a PA phoneme **m* for this case (**semʔ*) or simply attribute the observed variety of reflexes to the marginal phoneme **ŋ*2 postulated by Krauss and Leer (1981: 14-15) for cases when nasalization is accompanied with occasional and irregular labialization. The solution with **ŋ*2 (which we interpret as proper *ŋ*) seems more parsimonious.

**Eyak.** *laʔχc’-ɫ*, looks like a deverbative formation.

**Proto-Athabaskan-Eyak**. **sVŋ ~ *cʰVŋ* (Athabaskan).

**Proto-Eskimo**. Despites its scanty attestations, the best candidate is **aɣ-yaʁ* (Fortescue et al. 2010: 10), meaning ‘star’ in some Yupik lects, final **-yaʁ* is the non-productive suffix ‘place or thing where action takes place’. In other Yupik lects, it was superseded with a new formation derived from **iʁa-liʁ-* ‘moon shining (*vel sim.*)’ (Fortescue et al. 2010: 157). In Inuit, it was superseded with the new formation from **umɫuʁ* ‘day’ (Fortescue et al. 2010: 404).

**Proto-Aleut**. **sða-χ* (Bergsland 1994: 355; Golovko 1994: 105, 212), attested in Eastern and Atkan. In Attuan superseded with the deverbative from **siðʁi-sax-* ‘to shine’ (Bergsland 1994: 357).

**Proto-Chukotian**. **äŋär* (Fortescue 2005: 35), retained in all languages except for Kerek. Cognate to the Itelmen term.

**Proto-Itelmen**. **əŋezʸ* (Volodin, Khaloimova 2001: 156; Fortescue 2005: 36). Cognate to the Chukotian term.

**Proto-Nivkh**. **uɲ-ɣr* (Fortescue 2016: 156; Savelyeva, Taksami 1965: 161; Savelyeva, Taksami 1970: 394).

**Proto-Samoyed**. **kɨnsV-kɒyə̆* (Helimski 1997: 278), retained in Mator, Kamass and Selkup, goes back to Proto-Uralic **kunʸsʸV* ‘star’.

**Proto-Yukaghir**. Not reconstructible: words for ‘star’ in Yukaghir languages are derivatives or compounds with the root ‘hole’ (in fact, different roots for ‘hole’ in the two Yukaghir languages), reflecting the regional folklore motif “star as holes in the sky”.

81. ‘stone’.

**Proto-Yeniseian**. **ciʔ-s* (S. Starostin 1995: 217). Preserved in all daughter languages. The original paradigm is reconstructible as sg. **ciʔs*, pl. **ciʔŋ*; this means that **-s* is most likely a fossilized singulative suffix (cf. a similar case with the word for ‘eye’ q.v.).

**Proto-Athabaskan**. **c^he:* is retained in all three branches.

**Eyak.** *c^ha:*, cognate to Athabaskan.

**Proto-Athabaskan-Eyak**. **c^hV:* (Athabaskan, Eyak).

**Proto-Eskimo**. **uyaʁa-ɣ* (Fortescue et al. 2010: 420), retained in Yupik and Inuit.

**Proto-Aleut**. There are two almost equal candidates with generic semantics ‘stone’: **quɣa-na-χ* (Bergsland 1994: 332; Golovko 1994: 83, 216), attested in all branches; **nuχ* (Bergsland 1994: 284), attested in Eastern and Attuan. Difference is unclear, we treat them as synonyms.

**Proto-Chukotian**. **ɣəv-ɣəv* (Fortescue 2005: 93), reduplicated stem, retained in all languages. Cognate to the Itelmen term.

**Proto-Itelmen**. **wa-* (Volodin, Khaloimova 2001: 19, 160; Volodin 1976: 28, 129; Fortescue 2005: 93). Cognate to the Chukotian term. The Western Itelmen forms *kox* ‘stone’ (Ono 2003: 98), *kov-* ‘stone’ (Stebnitsky 1934: 91) can reflect the same proto-root.

**Proto-Nivkh**. **baʁ* (Fortescue 2016: 21; Savelyeva, Taksami 1965: 175; Savelyeva, Taksami 1970: 255).

**Proto-Samoyed**. **pɒy* (Janhunen 1977: 112) is retained in Northern Samoyed, Kamass and Selkup.

**Proto-Yukaghir**. Kolyma **söy-l* ‘small stone’ (Nikolaeva 2006: 410) vs. Tundra **qay-l* ‘stone’ (Nikolaeva 2006: 390). Cf. also Kolyma **pe:* ‘mountain / big stone’ (Nikolaeva 2006: 344–345), possibly connected with the Proto-Samoyed word for ‘stone’.

82. ‘sun’.

**Proto-Yeniseian**. **xiɢ-a* (S. Starostin 1995: 296). Preserved in all daughter languages. Proto-Yeniseian **xiɢ-a* ‘sun’ formally looks like an old suffixal derivative from **xiʔɢ* ‘day’ (S. Starostin 1995: 296) > Ket *iʔ*, etc.

**Proto-Athabaskan**. **ʂa̹:* ‘sun / moon’ is retained in all three branches. The retroflex shibilant points to PAE **x^w*.

**Eyak.** *qətə=kəɫ*, a descriptive formation ‘place of shriveling’ *vel sim*.

**Proto-Athabaskan-Eyak**. **x^wa:* (Athabaskan).

**Proto-Eskimo**. There are two candidates which are attested in both Yupik and Inuit in the criss-crossed configuration: **maca-ʁ* (Fortescue et al. 2010: 201) and **ciqi-nəʁ* (Fortescue et al. 2010: 91). We treat them as synonyms.

**Proto-Aleut**. **aʁa-ðaχ* (Bergsland 1994: 36; Golovko 1994: 270), attested in all branches. Derived from **aʁa-l-* ‘to become open; to open, clear up (of sky); to become visible’.

**Proto-Chukotian**. **tiðkə-tið* (Fortescue 2005: 285), reduplicated stem, retained in all languages except for Kerek.

**Proto-Itelmen**. **kulač* (Volodin, Khaloimova 2001: 208; Fortescue 2005: 393). Apparently with polysemy ‘sun / moon’ (see notes on ‘moon’).

**Proto-Nivkh**. **keŋ* (Fortescue 2016: 84; Savelyeva, Taksami 1965: 399; Savelyeva, Taksami 1970: 130).

**Proto-Samoyed**. **kɒyɒ* (Janhunen 1977: 58), retained as the word for ‘sun’ in all daughter languages save Selkup, where its meaning shifted to ‘heat’. Related to Finno-Ugric verb **kaya-* ‘appear, become visible’.

**Proto-Yukaghir**. Kolyma **puɣö* (Nikolaeva 2006: 366) vs. Tundra **yerpəɣə* (Nikolaeva 2006: 189). Modern Kolyma word for ‘sun’ reflects **yel^y-o:-nč^yə*, an active participle of an unattested passive verb **yel^y-o:-* from the root **ye:l^yə-* ‘to boil’ (Nikolaeva 2006: 187). This word replaced earlier **puɣö* (the same root as ‘warm’, q.v.), whose reflex in modern Kolyma Yukaghir means ‘summer’. Old Kolyma *pugu* ‘sun’ is attested in several wordlists; Jochelson notes that it is “an ancient word” (Jochelson 1926: 141) as opposed to modern *yelʸo:dʸə*. The meaning ‘sun’ is also retained in such modern Kolyma compounds as *puge-d-andʸə* ‘tsar [lit. sun chief]’ and *pugu-d-onora:* ‘rainbow [lit. sun tongue]’.

83. ‘to swim’.

**Proto-Yeniseian**. **su:y* (S. Starostin 1995: 279). Preserved in all daughter languages where it is attested, but not found in Arin or Pumpokol. In Kott, the verb only exists in conjunction with *ul* ‘water’; this may be a hint at some earlier meaning, but it might just as well be a Kott innovation, carried out in order to reduce homonymy with multiple other words that have the same phonetic shape (*šuy* ‘moon’, *šuy* ‘midge’, etc.).

**Proto-Athabaskan**. The best candidate is **=we:* retained in Pacific Coast and Northern.

**Eyak.** *=we*, cognate to Athabaskan.

**Proto-Athabaskan-Eyak**. **=we:* (Athabaskan, Eyak).

**Proto-Eskimo**. **puɣə-mə-* (Fortescue et al. 2010: 291), retained in all branches.

**Proto-Aleut**. **ɣučix-s* ~ **učix-s* (Bergsland 1994: 165; Golovko 1994: 245), attested in Eastern and Atkan, fluctuation of *ɣ-* is unlear. Cf. Attuan *taʁuχ-s-* ‘to swim’ (Bergsland 1994: 384).

**Proto-Chukotian**. **alʁeq-at-* (Fortescue 2005: 20), retained in all languages.

**Proto-Itelmen**. **ʍi-* (Volodin, Khaloimova 2001: 96, 186; Volodin 1976: 341; Fortescue 2005: 328). The specific meaning is ‘to swim downstream’, but this verb also functions as a generic term for ‘to swim’.

**Proto-Nivkh**. There are two candidates, both are attested with the meaning ‘to swim (of human) / to bath’ in Amur and East Sakhalin. First, **mrə-* (Fortescue 2016: 108; Savelyeva, Taksami 1965: 296; Savelyeva, Taksami 1970: 194). Second, **təm-* (Fortescue 2016: 149; Savelyeva, Taksami 1970: 325; Taksami 1996: 72, 160). We are forced to treat them as synonyms.

**Proto-Samoyed**. **u-* (Janhunen 1977: 29), retained in Nenets, Mator and Selkup, goes back to Proto-Uralic **uyi-* ‘to swim’.

**Proto-Yukaghir**. Not reconstructible due to insufficient attestation.

**Proto-Burushaski**. Yasin *minyˈa-* ‘to swim’ is the primary verbal stem which should represent the Proto-Bur. term. The Hunza compound expression *tam del-* ‘to wash, bathe, swim’, literally ‘to hit *tam*’ looks like a recent introduction. The bound noun *tam* ‘?’ also appears in the Shina complex verbs for ‘to swim’ and ‘to wash’, but the direction of borrowing was apparently from Burushaski to Shina, since, first of all, Shina *tam* lacks Indo-European etymology, and second, *tam* is only attested in the Gilgiti and Astori dialects of Shina which are the ones most influenced by Burushaski.

84. ‘tail’.

**Proto-Yeniseian**. **puɢ-aʒ* (S. Starostin 1995: 253). Preserved in all daughter languages, but not attested in Pumpokol. The stem ends in the same morpheme that is also found in Ket *ulʸ-et* = Kott *ul-ay* ‘rib’ and several other words denoting body parts; this seems to be an old fossilized suffix.

**Proto-Athabaskan**. **=kʰʸe-ʔ* is retained in all three branches, cognate to Eyak *=kə=kʰaʔ* ‘tail (of a bird)’.

**Eyak.** *=k=ƛ’ah*, a descriptive formation based on *=ƛ’ah* ‘rear, back end’.

**Proto-Athabaskan-Eyak**. **kʰV* (Athabaskan).

**Proto-Eskimo**. **pamyu-ʁ* (Fortescue et al. 2010: 272), retained in Yupik and Inuit.

**Proto-Aleut**. **hit-xi-χ* (Bergsland 1994: 217; Golovko 1994: 284), attested in Eastern and Atkan. Derived from **hit-* ‘to go out, come out; to come out, grow (of plant)’. Cf. Attuan *kimaʁu-χ* ‘tail (generic)’ (Bergsland 1994: 239).

**Proto-Chukotian**. **ŋoyŋ-ən* (Fortescue 2005: 198), retained in all languages. Cognate to the Itelmen term.

**Proto-Itelmen**. **ŋosx* (Volodin, Khaloimova 2001: 219; Fortescue 2005: 198). Cognate to the Chukotian term.

**Proto-Nivkh**. **ŋaki* ~ **ŋa=ki* (Fortescue 2016: 119; Savelyeva, Taksami 1965: 446; Savelyeva, Taksami 1970: 235). Can be derived from **ŋak* 'cartilage' (Fortescue 2016: 119; Savelyeva, Taksami 1970: 235). Alternatively a prefixal element **ŋa-*, common for body part names, can be singled out.

**Proto-Samoyed**. **tə̆ywɒ* (Janhunen 1977: 150) is retained in all daughter languages save Enets.

**Proto-Yukaghir**. **laq-il* (Nikolaeva 2006: 234–235) is retained in both Yukaghir languages.

**Proto-Burushaski**. Yasin *=ʃˈilan* 'tail' (no Hunza cognates) looks more archaic, since Hunza *=sˈum-al* 'tail' is derived from Common Burushaski **sum* 'sprout, shoot'.

85. 'that'.

**Proto-Yeniseian**. We tentatively reconstruct the tripartite system **ʔi* 'this' / **ʔu* 'that (medial)' / **ʔa* 'that (distal)', for Proto-Yeniseian, although it has been subjected to various modifications in modern languages.

**Proto-Athabaskan**. **yV:* is retained at least in Pacific Coast and Northern.

**Eyak.** *ʔəw* ~ *ʔu:*.

**Proto-Athabaskan-Eyak**. **yV* (Athabaskan), **ʔVw* (Eyak).

**Proto-Eskimo**. The synchronic Eskimo systems of the demonstrative pronouns are very complex, the same complexity is to be reconstructed for Proto-Eskimo (Fortescue et al. 2010: 497). For our purposes, we choose pronouns referring to visible non-moving compact objects on the horizontal axe: **uv-* 'this' / **iŋ-* 'that (medial)' / **ik-* 'that (distal)'.

**Proto-Aleut**. The Aleut deictic system (Bergsland 1997: 72; Golovko et al. 2009: 148– 149) is even more complex than the Eskimo ones. For our purposes, we choose pronouns referring to visible non-moving compact objects on the horizontal axe in front of the speaker: **u-ka* 'this' (Bergsland 1994: 426) / **a-ka* 'that (situated longitudinally)' (Bergsland 1994: 41) ~ **i-ka* 'that (situated transversally)' (Bergsland 1994: 187).

**Proto-Chukotian**. The basic system of demonstrative pronouns in the Chukotian languages in ternary: 'this' / 'that (medial, or close to the listener)' / 'that (distal, or far from the speaker and listener)'. On the basis of the Chukchi (Skorik 1961: 138), Koryak (Zhukova 1972: 191) and Alutor (Nagayama 2003: 304) data, the proto-system can be safely reconstructed as **ŋut-* ~ **ɣut-* 'this' (Fortescue 2005: 199) / **ən-* 'that (medial), 3rd p. pronoun' (Fortescue 2005: 342) / **ŋan-* 'that (distant)' (Fortescue 2005: 193). The only discrepancy between lects concerns the fluctuation of the initial consonant in the proximal pronoun: **ŋ-* in Chukchi and **ɣ-* in Koryak-Alutor.

**Proto-Itelmen**. According to Volodin 1976: 170, the system is binary: **tiʔ-* 'this' (Volodin, Khaloimova 2001: 223; Fortescue 2005: 200) / **nu-* 'that' (Volodin, Khaloimova 2001: 214; Fortescue 2005: 342).

**Proto-Nivkh**. Despite the fact that the full system of demonstrative pronouns is described as a complex opposition with several degrees of remoteness (Gruzdeva 1998: 26; Panfilov 1962: 241), it is likely that the basic system is to be reconstructed as a ternary one: **du-* 'this' (Fortescue 2016: 46) / **hu-* 'that (medial)' (Fortescue 2016: 77) / **a-* 'that (distal)' (Fortescue 2016: 7).

**Proto-Samoyed**. The reconstruction of Proto-Samoyed demonstratives is rather complicated. It involves at least three stems: **ta-* ~ **tä-* (Janhunen 1977: 150), **tə̆-* (Janhunen 1977: 144) and **ti-* (Janhunen 1977: 160–161). We tentatively reconstruct the basic opposition as that between distal **ta-* ~ **tä-* and proximal **tə̆-*. Such an opposition is directly preserved in Kamass and Selkup, while Northern Samoyed languages suffered various restructurings of the system. It is possible, but not certain, that **ti-* functioned as medial demonstrative.

**Proto-Yukaghir**. The Proto-Yukaghir system of demonstratives can be reconstructed as ternary: **ta-ŋ* (Nikolaeva 2006: 423–424) 'that (distal)', **an=ti-ŋ* ~ **an=tu-ŋ* (Nikolaeva 2006: 104) 'that (medial)', **ti-ŋ* (Nikolaeva 2006: 429–430) ~ **tu-ŋ* (Nikolaeva 2006: 437) 'this (proximal)'. Since medial demonstratives are derived from proximal ones by addition of a proclitic **an=*, we list only distal and proximal demonstratives.

86. 'this'.

**Proto-Yeniseian**. **ʔi*.

**Proto-Athabaskan**. **ti:* is retained in all three branches.

**Eyak.** *ʔəl ~ ʔã:*.

**Proto-Athabaskan-Eyak**. **ti:* (Athabaskan), **ʔVn* (Eyak).

**Proto-Eskimo**. **uv-*.

**Proto-Aleut**. **u-ka*.

**Proto-Chukotian**. **ŋut-* ~ **ɣut-*.

**Proto-Itelmen**. **tiʔ-*.

**Proto-Nivkh**. **du-* (Fortescue 2016: 46), see notes on 'that'.

**Proto-Samoyed**. **tə̆-* (Janhunen 1977: 144), see notes on 'that'.

**Proto-Yukaghir**. Although the basic stem for proximal demonstrative is **ti-ŋ* (Nikolaeva 2006: 429–430) in Kolyma and **tu-ŋ* (Nikolaeva 2006: 437) in Tundra, both stems are actually attested in both Yukaghir languages, and the semantic difference between them in Proto-Yukaghir is not clear.

87. 'thou'.

**Proto-Yeniseian**. **ʔaw* (S. Starostin 1995: 185). Preserved in all daughter languages. The "diphthongal" structure of this pronoun is rather unique for Proto-Yeniseian, so the regularity of the correspondences cannot be ascertained, but no better reconstruction can probably explain the discrepancy between Ket-Yugh **ʔu* and Kott-Arin **au*. The form **ʔaw* represents the direct stem of the 2nd p. sg. pronoun. The etymologically different oblique stem, lost in Kott-Arin, is still preserved in Ket-Yugh as **ʔuk* (possessive pronoun: 'your') or **ku* (verbal prefix of subject or object). These forms may have been influenced by Ket-Yugh **ʔu* 'you', but their velar constituent is completely autonomous, and there is no direct or indirect evidence that it was, at any time, present in the direct stem as well.

**Proto-Athabaskan**. **ɲe-n* is retained in all three branches.

**Eyak.** *ʔi:*, cognate to Athabaskan with **ɲ > 0*.

**Proto-Athabaskan-Eyak**. **ɲV* (Athabaskan, Eyak).

**Proto-Eskimo**. **əɬ=və-* ~ **əɬ=pə-* (Fortescue et al. 2010: 116), retained in all branches. The 2nd p. pl. pronoun is **əɬ=və-ci* 'you'. Origin and function of initial **əɬ=* are unclear, but the main meaningful morpheme is **=və-* ~ ** =pə-*, cf. the 2nd p. subj. exponent **-və-C-* (sg.), **-v-ci-* (pl.) in some verbal forms (Fortescue et al. 2010: 491, 494).

**Proto-Aleut**. **t(x)i=n* (Bergsland 1994: 397; Bergsland 1997: 57), attested in all branches. Meaning 'you (2 sg.) / (s)he self (3 sg. reflexive)'. The meaningful element is *-n*, see notes on 'I'.

**Proto-Chukotian**. **ɣəð* [direct] / **ɣən-* [obl.] (Fortescue 2005: 142), retained in all languages. Cognate to the Itelmen paradigm with the same suppletion.

**Proto-Itelmen**. **kəzʸa* [direct] / **kni-* [obl.] (Volodin, Khaloimova 2001: 236; Fortescue 2005: 142).

**Proto-Nivkh**. **ti* (Fortescue 2016: 32; Gruzdeva 1998: 25; Panfilov 1962: 231). Cf. **ti-n* 'you (pl.)' (Fortescue 2016: 33).

**Proto-Samoyed**. **tə̆-n* (Janhunen 1977: 147) is retained almost everywhere with the notable exception of (Tundra and Forest) Nenets, where 'thou' is etymologically 'thine body' and Forest Enets, where 'thou' is borrowed from Ket. Goes back to Proto-Uralic **ti-n* 'thou'.

**Proto-Yukaghir**. **tə-t* (Nikolaeva 2006: 429) is retained in both Yukaghir languages.

88. ‘tongue’.

**Proto-Yeniseian**. **ʔey* (S. Starostin 1995: 187). Preserved in Ket-Yugh and Pumpokol. In Proto-Kott-Arin, **ʔey* was replaced by **alup* (vocalism provisionally follows the Kott form rather than the controversial Arin variants), of unclear origin. Proto-Yeniseian **ʔey* ‘tongue’ is still preserved in Kott *ey*, pl. *e:y-aŋ*, but only in the meaning ‘voice; sound’; since the semantic shift ‘tongue’ > ‘voice’ (the actual meaning in Castrén’s vocabulary may have been ‘speech, language’) is more probable than the opposite, this increases the chances of **ʔey* as the original Proto-Yeniseian equivalent for ‘tongue’.

**Proto-Athabaskan**. **=cʰu-ʔ* is retained in all three branches, normally as the compound **=cʰu:-la:* with **=la-ʔ ~ *=la:* ‘tip’ (in some lects, the compound was reanalyzed as a root **=cʰu:l*).

**Eyak.** *=laʔt’*, cognate to Tlingit *ł’ú:t’* ‘tongue’, thus represent the Proto-Athabaskan-Eyak term (the marginal Eyak variant *=naʔt’* is unclear).

**Proto-Athabaskan-Eyak**. **łVt’* (Eyak).

**Proto-Eskimo**. The main candidates are Yupik-Sirenik **ulu* (Fortescue et al. 2010: 400) and Inuit **uqa-ʁ* (Fortescue et al. 2010: 411). We treat them as synonyms.

**Proto-Eskimo**. The main candidate is **ulu* which means ‘tongue’ in Yupik-Sirenik and ‘woman’s semi-lunar knife’ in Inuit (Fortescue et al. 2010: 400). In Yupik, the meaning ‘woman’s semi-lunar knife’ is expressed by the suffixed stem **ulu-ʁ-aʁ*, lit. ‘tongue-like’. In Inuit, **ulu* ‘tongue’ has been superseded with the stem **uqaʁ* ‘tongue / to speak’ (Fortescue et al. 2010: 411) without Yupik-Sirenik cognates.

**Proto-Aleut**. **umsu-χ* (Bergsland 1994: 442; Golovko 1994: 290), attested in Atkan and Attuan as ‘tongue’, having shifted into the meanings ‘flukes, whale’s tail; lap, blade as support’ in Eastern. Superseded with **aɣna-χ* (Bergsland 1994: 27) in Eastern, without cognates in other dialects.

**Proto-Chukotian**. **yilə-yil* (Fortescue 2005: 115), reduplicated stem, retained in all languages. Cognate to the Itelmen stem.

**Proto-Itelmen**. **il-čil* (Volodin, Khaloimova 2001: 224; Fortescue 2005: 115), reduplicated stem. Cognate to the Chukchi stem.

**Proto-Nivkh**. **hilɣ* (Fortescue 2016: 74; Savelyeva, Taksami 1965: 467; Savelyeva, Taksami 1970: 428). Perhaps derived from (or contaminated with) the verb **helel-* ‘to lick’ (Fortescue 2016: 72), if one analyzes the noun as **hil-ɣ* with the rare deverbal suffix **-x* (Panfilov 1962: 61) and the verb as a partly reduplicated stem **hel-el-*.

**Proto-Samoyed**. **käə̆ ~ *käə̆y* (Janhunen 1977: 66), retained in all daughter languages except Tundra Nenets, goes back to Proto-Uralic **käli* ‘tongue’.

**Proto-Yukaghir**. **wonor* (Nikolaeva 2006: 458) is retained in both Yukaghir languages.

89. ‘tooth’.

**Proto-Yeniseian**. **ʔi:ti* (S. Starostin 1995: 195). Preserved in all daughter languages, but not attested in Pumpokol.

**Proto-Athabaskan**. **=ʁu-ʔ* is retained in all three branches.

**Eyak.** *=χũ:-łə-yah*, the meaningful element is here the bound morpheme *χũ: ~χu:l* ‘tooth’ which is apparently cognate to Athabaskan **=ʁu-ʔ* ‘tooth’ and Tlingit *ʔu:χ* ‘tooth’, although the origin of the final nasal reflected in Eyak *χũ: ~χu:l* (*l* < **n*) is unclear.

**Proto-Athabaskan-Eyak**. **χu:* (Athabaskan, Eyak?). Apparently cognate to Tlingit *ʔu:χ* ‘tooth’ with a metathesis in either Proto-Tlingit or Proto-Athabaskan-Eyak.

**Proto-Eskimo**. **kəɣ-un* (Fortescue et al. 2010: 180), retained in all branches. Derived from **kəɣə-* ‘to bite’ (q.v.) with the instrumental suffix.

**Proto-Aleut**. **aɣa-lu-χ* (Bergsland 1994: 21; Golovko 1994: 15, 213), attested in all branches. In Eastern, tends to be superseded with the new instrumental formation *kiɣ-usi-χ* (Bergsland 1994: 238) from **kix-s-* ‘to bite’ (q.v.).

**Proto-Chukotian**. **wannə* (Fortescue 2005: 323), retained in all languages, although tends to be superseded with **rətnə* 'horn' (q.v.) in Chukchi.
**Proto-Itelmen**. **k'əp-* (Volodin, Khaloimova 2001: 157; Fortescue 2005: 396).
**Proto-Nivkh**. **ŋa=ɣzər* (Fortescue 2016: 118; Savelyeva, Taksami 1965: 163; Savelyeva, Taksami 1970: 233). A prefixal element **ŋa-*, common for body part names, can be singled out.
**Proto-Samoyed**. **timä* (Janhunen 1977: 163) is retained in all daughter languages.
**Proto-Yukaghir**. Kolyma **toð-iː*, derived from **toð-* 'to bite' (Nikolaeva 2006: 432) vs. Tundra **salʸqəriː* (Nikolaeva 2006: 394).

90. 'tree'.
**Proto-Yeniseian**. **ʔɔksi* (S. Starostin 1995: 198). Preserved everywhere except in Pumpokol, where the suppletive plural has replaced the old singular form. The word 'tree' was suppletive on the Proto-Yeniseian level; the plural form is reconstructed as **xaʔq* > Ket-Yugh **ʔaʔq*, Kott *ak* ~ *ax*, Pumpokol *hox-* in *hox-on*.
**Proto-Athabaskan**. **kʰʸen* is retained in all three branches.
**Eyak.** *lis*, can be a Russian loanword.
**Proto-Athabaskan-Eyak**. **kʰen* (Athabaskan), this root is retained in Eyak as *t=kʰĩh* 'stick; wood (timber)'; because the semantic shift 'tree' > 'timber' is much more frequent than *vice versa*, **kʰen* is to be posit as the Proto-Athabaskan-Eyak term for 'tree'.
**Proto-Eskimo**. **napa-ʁ-aq-tuʁ* (Fortescue et al. 2010: 236), a Yupik-Inuit derivative from **napa-* 'to be standing (upright)'.
**Proto-Aleut**. Not reconstructible.
**Proto-Chukotian**. **uttə* (Fortescue 2005: 310), retained in all languages.
**Proto-Itelmen**. **uʔ* (Volodin, Khaloimova 2001: 149; Fortescue 2005: 310).
**Proto-Nivkh**. **d̦iɣar* (Fortescue 2016: 52; Savelyeva, Taksami 1965: 126; Savelyeva, Taksami 1970: 353). Polysemy 'tree / firewood'. Morphologically unclear.
**Proto-Samoyed**. **pa* (Janhunen 1977: 117), retained in all daughter languages, goes back to Proto-Uralic **pawi* 'tree'.
**Proto-Yukaghir**. **saː-l* (Nikolaeva 2006: 392) is retained in both Yukaghir languages.
**Proto-Burushaski**. Borrowed from Indo-Aryan sources.

91. 'two'.
**Proto-Yeniseian**. **xɨn-a* (S. Starostin 1995: 296). Preserved in all daughter languages. Initial **x-* is reconstructed based on the presence of back consonants in Arin and Pumpokol. The suffix *-a* is a common element in the formation of Yeniseian numerals; the original root is simply **xɨn-*.
**Proto-Athabaskan**. **naː-* is retained in all three branches, regularly accompanied with various suffixes.
**Eyak.** *laʔt-*, comparison with Athabaskan **nV-* suggests that historically *-t-* is a fossilized suffix.
**Proto-Athabaskan-Eyak**. **naː-* (Athabaskan, Eyak).
**Proto-Eskimo**. **malʁu-ɣ* (Fortescue et al. 2010: 205), retained in all branches.
**Proto-Aleut**. **aːlax* ~ **ulax* (Bergsland 1994: 49, 570; Golovko 1994: 200), attested in all branches. Can be derivative from **a-/u-* 'to be'.
**Proto-Chukotian**. **ŋiðä-q* (Fortescue 2005: 197), retained in all languages.
**Proto-Itelmen**. **kasχ* (Volodin, Khaloimova 2001: 148; Fortescue 2005: 397).
**Proto-Nivkh**. **me* (Fortescue 2016: 103, 178; Panfilov 1962: 181, 214–215; Gruzdeva 1998: 24).
**Proto-Samoyed**. **kitä* (Janhunen 1977 71–72), retained in all daughter languages, goes back to Proto-Uralic numeral 'two'.
**Proto-Yukaghir**. **ki-* (Nikolaeva 2006: 209) is retained in Tundra, Omok, and Chuvan. Kolyma has **antəq-* ~ **wantəq-* (Nikolaeva 2006: 110–111) of unclear origin. The reflex of Proto-Yukaghir **ki-* 'two' is preserved in the word for 'seven': Kolyma *pur-ki-*,

Tundra *pus-ki-*, originally ‘two on [five]’ (Nikolaeva 2006: 365). Because of this, **antəq- ~ *wantəq-* must be an innovation.

92. ‘to go’.

**Proto-Yeniseian**. Proto-Yeniseian **hey-* ‘to go’ (S. Starostin 1995: 231) is reconstructible based on the Ket-Yugh infinitive (verbal noun) form **ʔey-iŋ* ‘to go’ and the exactly corresponding Kott infinitive *hey-aŋ*. The second, more hypothetical, reconstruction **=ʒe- ~ *=ʒen* reconciles two of the most basic Ket-Yugh and Kott equivalents for the meaning ‘to go’, namely, the Kott root **=in-* and Ket-Yugh **=de(n)*; in Kott, according to S. Starostin’s correspondences, **=ʒen* should have yielded **=yen*, with subsequent contractions (**i=yen-aŋ* ‘I go’ > *iːnaŋ*, etc.).

**Proto-Athabaskan**. **=haː* ‘to go / to come’ is retained in all three branches.

**Eyak.** *=a*, cognate to Athabaskan, see note on ‘to come’.

**Proto-Athabaskan-Eyak**. **=haː* (Athabaskan, Eyak).

**Proto-Eskimo**. Cannot be reconstructed with certainty. Cf. the stable verb **pi-yuɣ-* ‘to walk’ (Fortescue et al. 2010: 289).

**Proto-Aleut**. **huya-l-* (Bergsland 1994: 455; Golovko 1994: 213), attested in all branches.

**Proto-Chukotian**. **təlä* (Fortescue 2005: 295), retained as a basic term at least in Chukchi and Koryak.

**Proto-Itelmen**. **il-* (Volodin, Khaloimova 2001: 119, 158; Fortescue 2005: 247), meaning ‘to go / to go away’. Distinct from **łale-* ‘to walk’ (Volodin, Khaloimova 2001: 51; Fortescue 2005: 295).

**Proto-Nivkh**. **wi-* (Fortescue 2016: 163; Savelyeva, Taksami 1965: 164; Savelyeva, Taksami 1970: 54).

**Proto-Samoyed**. **men-* (Janhunen 1977: 94), retained in Nenets, Nganasan and Kamass, goes back to Proto-Uralic **meni-* ‘to go’.

**Proto-Yukaghir**. **qon-* (Nikolaeva 2006: 385–386), retained in Kolyma Yukaghir. Tundra Yukaghir preserves two derivatives from this root: *qanaː-* ‘to roam away (of nomads)’ and *qande-* ‘to accompany’. The main Tundra verb for ‘to go’, *uː-*, goes back to polysemous verb **uy-* (Nikolaeva 2006: 441–442), whose Kolyma reflex means ‘to work, to do’. It is unlikely that **uy-* was originally a verb of movement. **qon-* is related to, or borrowed from, Proto-Samoyed **kɒn-* ‘to go (away)’ (Janhunen 1977: 59–60).

**Proto-Burushaski**. Out of three roots involved in the Yasin and Hunza suppletive paradigms, two are present in both dialects: **ne-* & **gal-*. We reconstruct these two for the Proto-Burushaski paradigm.

93. ‘warm’.

**Proto-Yeniseian**. **xus-* (S. Starostin 1995: 299). Preserved in all daughter languages, with the probable exception of Kott, where there may have been a merger of the lexically distinct Proto-Yeniseian meanings ‘warm’ and ‘hot’ (Kott *fal ~ pʰal* is compared to such forms as Yugh *aːp*, Ket *aː* ‘heat’, etc. < Proto-Yeniseian **ʔap-* ‘hot’). The semantic opposition **xus-* ‘warm’ / **ʔap-* ‘hot’, best attested in Ket-Yugh, is probably of Proto-Yeniseian origin.

**Proto-Athabaskan**. The best candidate is **=zeɬ* retained in Pacific Coast and Northern.

**Eyak.** *=tə=χã*, based on the verb ‘to melt, thaw’, thus looks like a new formation.

**Proto-Athabaskan-Eyak**. **=seɬ* (Athabaskan).

**Proto-Eskimo**. **maqa-ʁ-* (Fortescue et al. 2010: 210), a Yupik-Sirenik term. In Inuit lects, ‘(to be) warm’ is usually derived from **uɣu-naʁ-* ‘to be burning hot’ (Fortescue et al. 2010: 395). The meaning ‘(to be) hot’ is usually expressed with the help of various derivatives from **uɣu-* ‘to be heated up, cooked’ (Fortescue et al. 2010: 394).

**Proto-Aleut**. **huʁ-naː-za-* (Bergsland 1994: 425; Golovko 1994: 167), attested in all branches. Derived from **huʁ-naː* ‘lee side of house’.

**Proto-Chukotian**. _*om-_ (Fortescue 2005: 205), meaning 'warm (of object / of weather)' in Koryak and Alutor, but only 'warm (of weather)' in Chukchi. Cognate to the Itelmen term. Distinct from _*təɣəl-_ 'hot' (Fortescue 2005: 293).

**Proto-Itelmen**. _*om-_ (Volodin, Khaloimova 2001: 213; Fortescue 2005: 205), meaning 'warm (of object / of weather)'. Cognate to the Chukchi term. Distinct from _*xka-_ 'hot' (Volodin, Khaloimova 2001: 146; Fortescue 2005: 339).

**Proto-Nivkh**. _*dak-_ (Fortescue 2016: 38; Savelyeva, Taksami 1965: 419; Savelyeva, Taksami 1970: 367), meaning 'to be warm'. Distinct from _*qav-_ 'to be hot' (Fortescue 2016: 140; Savelyeva, Taksami 1965: 119; Savelyeva, Taksami 1970: 146).

**Proto-Samoyed**. There are two candidates. The first, _*yu-pɒ_ (Janhunen 1977: 47–48), is derived from the verb _*yu-_ 'to be warm / to melt'. This is the main word for 'warm' in Nenets, Enets and Mator. The second candidate, _*päywä_ (Janhunen 1977: 120), means 'heat, warmth' in Nganasan and 'warm' in Selkup. Its Finnic and Saami cognates (< Proto-Uralic _*päywä_) mean 'sun, day'.

**Proto-Yukaghir**. _*puɣö_ 'warm, hot' (Nikolaeva 2006: 366) is retained in both Yukaghir languages.

94. 'water'.

**Proto-Yeniseian**. _*xurɪ_ (S. Starostin 1995: 298). Preserved in all daughter languages. Initial _*x-_ is reconstructed based on the velar reflexation _k-_ in Arin. For the difference between 'water' and 'rain', see notes on 'rain'.

**Proto-Athabaskan**. *$t^hu$: is retained in all three branches.

**Eyak.** _kiyah_, morphologically unclear; the old root is retained as the preverb $t^ha$ʔ 'into water'.

**Proto-Athabaskan-Eyak**. *$t^hV$: (Athabaskan). Leer (2008a) compares Proto-Athabaskan *$t^hu$: 'water' with Eyak $t^h$ãh 'wave' and reconstruct Proto-Athabaskan-Eyak *$t^ham$. Such a solution seems dubious since, firstly, it is based on a one-example rule _*Vm_ > Proto-Athabaskan _-u_, secondly, it implies that Proto-Athabaskan *$t^hu$: is unrelated to Proto-Athabaskan *$t^ha$:- 'water, into water (first element in compounds). The Proto-Athabaskan vowel alternation between *$t^hu$: and *$t^ha$:- is indeed unique, but Krauss and Leer (Krauss & Leer 1981: 87; Leer 1996) might be right analyzing Proto-Athabaskan *$t^hu$: 'water' as an old compound *$t^ha$:-_wV_ (the second element is unclear).

**Proto-Eskimo**. _*əmə-ʁ_ (Fortescue et al. 2010: 120), retained in all branches, the same stem as 'to drink' (q.v.).

**Proto-Aleut**. _*ta:ŋa-χ_ (Bergsland 1994: 392; Golovko 1994: 191), attested in all branches, the same stem as 'to drink' (q.v.).

**Proto-Chukotian**. _*miməl_ ~ _*mi=məl_ ~ _*iməl_ (Fortescue 2005: 99), retained in all languages.

**Proto-Itelmen**. _*iʔ_ (Volodin, Khaloimova 2001: 141; Fortescue 2005: 398).

**Proto-Nivkh**. _*ƫaʁ_ (Fortescue 2016: 30; Savelyeva, Taksami 1965: 88; Savelyeva, Taksami 1970: 444). Panfilov (1962: 61) proposes the analysis _*ƫa-ʁ_, but it is not certain.

**Proto-Samoyed**. _*wet_ (Janhunen 1977: 176), retained in all Samoyed languages, goes back to Proto-Uralic _*weti_ 'water'.

**Proto-Yukaghir**. Kolyma _*onč-i:_ (Nikolaeva 2006: 330–331) vs. Tundra _*law-yə_ (Nikolaeva 2006: 236). Both Kolyma and Tundra words for 'water' are derived from the respective verbs meaning 'to drink', q.v. Cf. also the root _*liŋkə_ (Nikolaeva 2006: 255), meaning 'rain' and 'to drink' in Chuvan and 'rain, water; to drink water' in Omok, but absent from modern Yukaghir languages.

95. 'we'.

**Proto-Yeniseian**. _*ʔaʒ-əŋ_ (S. Starostin 1995: 185). The Proto-Yeniseian equivalent for 'we' was clearly the regular plural form of 'I'; hence, see notes on 'I' for reconstruction peculiarities.

**Proto-Athabaskan**. _*nVh-_ is retained in all three branches.

**Eyak.** _ta:_ [direct stem] / $q^ha$: [oblique stem].

**Proto-Athabaskan-Eyak**. *nVh-* (Athabaskan), **tV* ~ *$q^hV$ (Eyak).

**Proto-Eskimo**. **uva=ku-t* (Fortescue et al. 2010: 418), retained in all branches. In some Yupik lects, phonetically contaminated with **uva=ŋa* 'I'. For desemanticized **uva-* see notes on 'I'; final *-t* is a regular plural suffix (cf. the same pattern in Aleut); the same *-*ku-t* is used as the 1st p. pl. subject exponent in verbal forms (Fortescue et al. 2010: 489).

**Proto-Aleut**. **t(x)i=ma-* (Bergsland 1994: 397; Bergsland 1997: 57), attested in all branches. Modified with the plural suffixes -(*i*)*s* or -(*i*)*n*. The meaningful element is *-ma-*, whereas *ti-* ~ *txi-* is a proclitic attached to the 1st, 2nd and reflexive 3rd person pronouns (Bergsland 1997: 56). Sometimes the forms *ti=ŋ-is* ~ *ti=ŋ-in* 'we' are used instead, literally 'I-PL' from *ti=ŋ* 'I'.

**Proto-Chukotian**. **mur-* (Fortescue 2005: 179), retained in all languages. Cognate to the Itelmen term.

**Proto-Itelmen**. *$muz^ya$ (Volodin, Khaloimova 2001: 236; Fortescue 2005: 179). Cognate to the Chukotian term.

**Proto-Nivkh**. All Nivkh lects possess the category of clusivity. The pronoun **ɲi-n* (Fortescue 2016: 114; Gruzdeva 1998: 25; Panfilov 1962: 231) means 'we (exclusive)'. In Amur, it was additionally modified with the standard plural suffix *-*kun*. The suffix *-n* can be singled out on the basis of comparison with **ɲi* 'I' q.v., **mer-n* 'we (inclusive)' and the mirroring pair **ti* 'thou' / **ti-n* 'you (pl.)'. See Panfilov 1962: 50–51 for rare *-*n* which expresses something related to humans or animate creatures in general. Additionally one can suspect that the plural suffix *-*kun* is to be historically analyzed as *-*ku-n* with the same *-n* as in the plural personal pronouns (thus Panfilov 1962: 95). The exclusive pronoun is opposed to **mer-n* 'we (inclusive)' (Fortescue 2016: 105; Gruzdeva 1998: 25; Panfilov 1962: 231). Panfilov (1962: 59) proposes to analyze it as **me-r-n* with the archaic suffix *-*r*, cf. the pronoun 'we (dual.)' which is likely based on the morpheme **me-* (Fortescue 2016: 103), but it requires additional investigation.

**Proto-Samoyed**. **me-* (Janhunen 1977: 91), retained in Nganasan, Mator, Kamass and Selkup, goes back to Proto-Uralic **me-* 'we'. Nenets and Enets replaced the original pronoun by dual/plural forms of 'I'.

**Proto-Yukaghir**. **mi-t* (Nikolaeva 2006: 269–270) is retained in both Yukaghir languages.

96. 'what?'.

**Proto-Yeniseian**. **si* ~ **ʔa=si* (S. Starostin 1995: 183). Preserved in all daughter languages where attested, but not found in Arin or Pumpokol. The Proto-Yeniseian morpheme clearly had the fricative *-*s-* as its main distinctive component, but the vocalic "framing" differs in between Ket-Yugh and Kott and is hard to reconstruct convincingly.

**Proto-Athabaskan**. The main candidates are **te* and **ye* which are intertwined between lects or used as single compounds **te-ye* or **ye-te* (**te* is also the main candidate for 'who' q.v.).

**Eyak.** *te:*, cognate to Athabaskan **te* and Tlingit *ta:-t* 'what'.

**Proto-Athabaskan-Eyak**. **te* (Athabaskan, Eyak).

**Proto-Eskimo**. **cu-* (Fortescue et al. 2010: 97), retained in all branches. Modified with various suffixes, most frequently **cu-na*.

**Proto-Aleut**. **alqu-* (Bergsland 1994: 55; Bergsland 1997: 80; Golovko 1994: 287), attested in all branches.

**Proto-Chukotian**. **ðäq* (Fortescue 2005: 56), retained in all languages.

**Proto-Itelmen**. **aŋqa* (Volodin, Khaloimova 2001: 222; Fortescue 2005: 399).

**Proto-Nivkh**. **tu-nt* (Fortescue 2016: 152; Gruzdeva 1998: 28; Panfilov 1962: 253).

**Proto-Samoyed**. **mə* (Janhunen 1977: 91), retained as an interrogative pronoun 'who' in Enets and Nganasan, goes back to Proto-Uralic **mV* 'what?'.

**Proto-Yukaghir**. **leme* (Nikolaeva 2006: 239) is retained in both Yukaghir languages alongside of the variant **neme*. Since both assimilation and dissimilation are typologically normal in such sequences, we take **leme* ~ **neme* as synonyms.

97. ‘white’.

**Proto-Yeniseian**. **tak-am* (S. Starostin 1995: 282). Preserved in all daughter languages, but morphologically restructured in Yugh. Probably derived from Proto-Yeniseian **tik* ‘snow’, but discrepancies in vocalism remain unexplainable.

**Proto-Athabaskan**. **=qay* is retained in all three branches.

**Eyak.** *xiƛ’-kaʔ*, literally ‘snow-like’, a transparent new formation.

**Proto-Athabaskan-Eyak**. **=qay* (Athabaskan).

**Proto-Eskimo**. **qatə-ʁ-* (Fortescue et al. 2010: 316), retained in Yupik and Inuit. In Yupik some lects, superseded with **qak-cuʁ-* (Fortescue et al. 2010: 303), derived from **qakə-* ‘be bleached’

**Proto-Aleut**. **qum̥a-l-* (Bergsland 1994: 335; Golovko 1994: 184), attested in all branches.

**Proto-Chukotian**. **ilɣə* (Fortescue 2005: 96), meaning ‘white / clean’, retained in all languages.

**Proto-Itelmen**. **atx-* (Volodin, Khaloimova 2001: 16, 135; Volodin 1976: 320; Fortescue 2005: 77), meaning ‘white / light, bright’.

**Proto-Nivkh**. **ɢon-u-* (Fortescue 2016: 67; Savelyeva, Taksami 1965: 65; Savelyeva, Taksami 1970: 140, 144).

**Proto-Samoyed**. **sər* (Janhunen 1977: 138) is retained in all daughter languages save Mator. Possibly connected to Proto-Samoyed **sər* ‘ice’.

**Proto-Yukaghir**. **poy-inə* (Nikolaeva 2006: 355) is retained in Kolyma as the main word for ‘white’ and in Tundra as a secondary synonym. Another, less likely, candidate is the main root for ‘white’ in Tundra, **nʸaːwə* (Nikolaeva 2006: 291), that has no cognates in Kolyma.

98. ‘who?’.

**Proto-Yeniseian**. **ʔan-* (S. Starostin 1995: 181). Preserved only in Ket-Yugh; the Kott system of interrogative pronouns seems to have been restructured.

**Proto-Athabaskan**. The main candidate is **te* which is retained in Pacific Coast and Northern having been accompanied with various extensions. PA **te* is also a good candidate for PA ‘what’ q.v.

**Eyak.** *tuː*, cognate to Tlingit *ʔaː-tuː* ‘who’ (relationship with Athabaskan **te* is unclear).

**Proto-Athabaskan-Eyak**. **tuː* (Eyak).

**Proto-Eskimo**. **ki-na* (Fortescue et al. 2010: 190), retained in all branches. The final element -*na* is detachable. Cognate to the Aleut term.

**Proto-Aleut**. **kiːn* (Bergsland 1994: 239; Bergsland 1997: 81; Golovko 1994: 221), attested in all branches. Comparison with Eskimo **ki-n* ‘who?’ suggests that the final *-n* is a fossilized suffix.

**Proto-Chukotian**. **mikä* (Fortescue 2005: 175), retained in all languages. Can be cognate to the Itelmen term.

**Proto-Itelmen**. **k’e-* (Volodin, Khaloimova 2001: 165; Fortescue 2005: 175). Can be cognate to the Chukotian term.

**Proto-Nivkh**. **nar* (Fortescue 2016: 111; Gruzdeva 1998: 28; Panfilov 1962: 253).

**Proto-Samoyed**. **ke-* (Janhunen 1977: 69), retained in Northern Samoyed, Mator and Kamass, goes back to Proto-Uralic **ku-* ‘who?’.

**Proto-Yukaghir**. **kin* (Nikolaeva 2006: 211–212) is retained in both Yukaghir languages.

99. ‘woman’.

**Proto-Yeniseian**. **qem* (S. Starostin 1995: 266). Preserved in Ket-Yugh and, most likely, in Arin; possibly also in Pumpokol, if the attested word for ‘wife’ in that language had the same root as ‘woman’. In Kott-Arin, there is another stem for the meanings ‘woman’ and ‘wife’, functioning on its own in Kott (*alit*) and as part of a compound with the older word for ‘woman’ in Arin (**qem-alit*, with various assimilations and reductions in the actual attested dialectal forms). There are no parallels for this **ʔalit* in Ket-Yugh, and it is not clear why Arin turned the old word into a compound, and Kott retained only the newer part of this compound, but from the point of view of cognate distribution, this is the most economic scenario.

**Proto-Athabaskan**. Besides various descriptive new formations, the main candidate is the widely attested compound **č̨’e:ʔ-qe:* ‘woman’; the first element is a bound root **č̨’eʔ* ‘female’ normally used as the second element of compounds, the second element is not attested elsewhere, but the comparison with Eyak proves its antiquity.

**Eyak.** *qʰeʔ-ɬ*, comparison with the Athabaskan data suggests that final *-ɬ* should be the common nominal suffix *-ɬ*.

**Proto-Athabaskan-Eyak**. **qʰe:* (Athabaskan, Eyak).

**Proto-Eskimo**. **aʁna-ʁ* (Fortescue et al. 2010: 47), retained in Yupik and Inuit.

**Proto-Aleut**. **ayaɣa-χ* (Bergsland 1994: 115; Golovko 1994: 207), attested in all branches.

**Proto-Chukotian**. **ŋäv-* (Fortescue 2005: 195) means ‘female’, expressions for ‘woman’ are based on it in all languages. Cognate to the Itelmen term.

**Proto-Itelmen**. **ŋim-sx* (Volodin, Khaloimova 2001: 153; Fortescue 2005: 195). Cognate to the Chukotian term.

**Proto-Nivkh**. **taŋq* (Fortescue 2016: 146; Savelyeva, Taksami 1970: 393; Peiros, Starostin 1986: 146). In Amur, superseded with unclear *umgu* ‘woman’ (Fortescue 2016: 158; Savelyeva, Taksami 1965: 140; Savelyeva, Taksami 1970: 393).

**Proto-Samoyed**. **ne* (Janhunen 1977: 100), retained in all Samoyed languages, goes back to Proto-Uralic **näɣi* ‘woman’.

**Proto-Yukaghir**. **pay* (Nikolaeva 2006: 340) is retained in both Yukaghir languages.

100. ‘yellow’.

**Proto-Yeniseian**. Not reconstructible: the Ket word *qɜlʸ-ay-sʸ* is transparently derived from ‘gall’, the Kott word *šuy* is the same as ‘moon’, the Pumpokol word *tul-si* is the same as ‘red’.

**Proto-Athabaskan**. **=cʰuχ* ‘green / yellow’ is retained in all three branches.

**Eyak.** *χəwa:-cʰeʔq’-kaʔ*, literally ‘dog urine-like’, a transparent new formation.

**Proto-Athabaskan-Eyak**. **=cʰuχ* (Athabaskan).

**Proto-Eskimo**. Cannot be reconstructed with certainty. Cf. Inuit **quq-yuɣ* ~ **quq-cuʁ* ‘yellow(ish)’, lit. ‘urine-like’ (Fortescue et al. 2010: 348).

**Proto-Aleut**. **čum-nux* (Bergsland 1994: 153; Golovko 1994: 152), attested in all branches, polysemy: ‘yellow / brown / gray’. Morphologically unclear.

**Proto-Chukotian**. Not reconstructible with certainty.

**Proto-Itelmen**. Not reconstructible with certainty. It is possible that **ʍɬ-* ‘green/blue’ (Fortescue 2005: 337) actually should be reconstructed with polysemy ‘green/blue/yellow’, cf. comm. on ‘green’.

**Proto-Nivkh**. **evrq-wala-* (Fortescue 2016: 57; Savelyeva, Taksami 1965: 140; Savelyeva, Taksami 1970: 476). Literally ‘tinder-colored’ from **evrq* ‘tinder, amadou’.

**Proto-Samoyed**. **tʊsV-* ~ **cpsV*. The word for ‘yellow’ can only be reconstructed for Northern Samoyed.

**Proto-Yukaghir**. **nʸor-inə-* (Nikolaeva 2006: 311), retained in Tundra. This is the only possible candidate, since Kolyma Yukaghir word for ‘yellow’ is derived from the noun ‘fox’. The root **nʸor-* may be compared to Proto-Samoyed **nʸar-* ‘red’ (Janhunen 1977: 107–108).

101. ‘far’.

**Proto-Yeniseian**. **bi:rɪ* (S. Starostin 1995: 211). Preserved in Ket-Yugh and Kott.

**Proto-Athabaskan**. **=za:t* is retained in all three branches.

**Eyak.** *qə=ƛa:=ʔa:w*, based on the verb *=ʔa:w* ‘long’, thus apparently a new formation.

**Proto-Athabaskan-Eyak**. **=sa:t* (Athabaskan).

**Proto-Eskimo**. **uŋa-ðiɣ-* (Fortescue et al. 2010: 408), an Inuit term, derived from the bound root **uŋa-* 'area beyond (partition)' plus the suffix **-ðiɣ* 'being far in a direction'. An unstable item in Yupik.

**Proto-Aleut**. **ama:-txa-l-* (Bergsland 1994: 60; Golovko 1994: 27), attested in Eastern and Atkan. Derived from the locative word **ama-* 'away, out of sight'.

**Proto-Chukotian**. **əɣava* (Fortescue 2005: 339), retained in all languages.

**Proto-Itelmen**. **meč'a-* (Volodin, Khaloimova 2001: 60; Volodin 1976: 340, 341, 342; Stebnitsky 1934: 102; Fortescue 2005: 257), meaning 'far' in Western, the Southern derivative verb means 'to stay with smbd., at smbd.'s place'. According to Volodin 1976: 340, 341, **meč'a-* 'far' forms a pair with **t'mal-* 'near' in Western. Distinct from **t'aɬ-* whose meaning is rather 'distant' (Volodin, Khaloimova 2001: 92; Volodin 1976: 339; Fortescue 2005: 283) and which seems a more marginal word in Western according to Volodin's data.

**Proto-Nivkh**. **tə-l-v-* (Fortescue 2016: 154; Savelyeva, Taksami 1965: 123; Savelyeva, Taksami 1970: 388). Derived from the verb **tə-* 'to be far' (Savelyeva, Taksami 1970: 387).

**Proto-Samoyed**. **kuntə̑-kɒ* (Janhunen 1977: 78), retained in all languages save Nenets, is derived from **kuntə̑* 'long, length'.

**Proto-Yukaghir**. **yu:-kə* (Nikolaeva 2006: 199) is retained in both Yukaghir languages.

102. 'heavy'.

**Proto-Yeniseian**. **səɢ-* (S. Starostin 1995: 273). Preserved in all daughter languages, but not attested in Pumpokol.

**Proto-Athabaskan**. **=ta:ʔs* is retained in all three branches.

**Eyak.** *=ta:s*, cognate to Athabaskan.

**Proto-Athabaskan-Eyak**. **=ta:s* (Athabaskan, Eyak).

**Proto-Eskimo**. **uqi-ma-ŋit-* (Fortescue et al. 2010: 414), retained in Yupik and Inuit. Eventually from **uqiɣ* 'heaviness', but morphological details are not entirely cleat (**-ŋit* is a negative suffix).

**Proto-Aleut**. **kayaɣ-na-l-* (Bergsland 1994: 234; Golovko 1994: 60), attested in Eastern and Atkan. According to the examples in Bergsland & Dirks 1978, this is the basic term for 'heavy' in Eastern. Derived from the bound root **kayaɣ-* 'heavy' (Bergsland 1994: 234). In Atkan, almost superseded with the new formation **iɣna-tu-l-* (Bergsland 1994: 179; Golovko 1994: 47), attested however in Eastern as well, which is derived from **iɣna-χ* 'weight'.

**Proto-Chukotian**. **itčə-* ~ **iččə-* (Fortescue 2005: 94), retained in all languages.

**Proto-Itelmen**. **kaz-* (Volodin, Khaloimova 2001: 215; Fortescue 2005: 369).

**Proto-Nivkh**. **ber-* (Fortescue 2016: 22; Savelyeva, Taksami 1965: 428; Savelyeva, Taksami 1970: 257).

**Proto-Samoyed**. **säc̮* (Janhunen 1977: 139) is retained in Northern Samoyed, Mator, Kamass and Selkup.

**Proto-Yukaghir**. Kolyma **niɣey-* ~ **niŋkəy-* (Nikolaeva 2006: 299) vs. Tundra **iralʸə-* (Nikolaeva 2006: 462).

103. 'near'.

**Proto-Yeniseian**. **ʔuti* ~ **xuti* (S. Starostin 1995: 201). Preserved only in Ket-Yugh. In Kott, PY **ʔuti* 'near' is preserved in the adverbial form *uti-ga* 'here'. Lack of parallels in Arin means that the Proto-Yeniseian equivalent of the Ket-Yugh forms could have been **ʔuti* or **xuti*.

**Proto-Athabaskan**. An unstable item. The possible candidate is the verb **=ten* 'to be near' (Pacific Coast), but the PA meaning and functions of this root were much wider, the postposition/enclitic **=ten* expressed a general locative semanics 'where' and 'when', thus the verbal use in the specific meaning 'to be near' is rather a Pacific Coast innovation. A more appropriate candidate is the postposition **=ʁa:* ‘near’ as attested in Northern and Apachean.

**Eyak.** *=ta:-* (postoposition). Also cf. the postposition *=χ* ‘in contact with, glancing, brushing, moving against point on, along part of, extending to point on, adhering to’ which is cognate to the PA postposition **=ʁa:* ‘near’.

**Proto-Athabaskan-Eyak**. **=χV* (Athabaskan), **=tV* (Eyak).

**Proto-Eskimo**. **qanə-t-* (Fortescue et al. 2010: 309), retained in Yupik and Inuit.

**Proto-Aleut**. **am̥i-ʁ-* (Bergsland 1994: 65; Bergsland & Dirks 1978: 86), probably the basic expression for ‘near’ in Eastern. In Atkan, the negated new formation *ama:txa-laka-n* ‘near’ is used (Bergsland 1994: 60; Golovko 1994: 27), lit. ‘not far’.

**Proto-Chukotian**. There are two similar candidates. First, **čəmčä-* (Fortescue 2005: 52), a basic root for ‘near’ in Chukchi (Inenlikei 2005), meaning ‘neighboring’ in Koryak. Second, **äymə- ~ *čäymə-* (Fortescue 2005: 28), a basic root for ‘near’ in Koryak (Zhukova 1990: 109) and Alutor (Kibrik et al. 2004: 518), meaning ‘to approach’ in Chukchi and Kerek. Fortescue is apparently correct that the original shape of the latter root is **äymə-* and the initial *č-* in is the result of influence on the part of **čəmčä-*. We treat both roots as synonyms.

**Proto-Itelmen**. **t’mal-* (Volodin, Khaloimova 2001: 136; Volodin 1976: 340, 341; Fortescue 2005: 298, 379), attested in Western, Sothern and Eastern.

**Proto-Nivkh**. **ma-la-* (Fortescue 2016: 100; Savelyeva, Taksami 1965: 69; Savelyeva, Taksami 1970: 173). Derived from **ma-* ‘to be near’ (Savelyeva, Taksami 1970: 172).

**Proto-Samoyed**. **wan-i-* (Helimski 1997: 301) is retained in Enets, Mator and Kamass.

**Proto-Yukaghir**. **me:-kə* (Nikolaeva 2006: 262–263) is retained in Kolyma; its Tundra cognate means ‘till, up to’. The Tundra word *eyuoke* ‘near’ means literally ‘not far’.

104. ‘salt’.

**Proto-Yeniseian**. **čəʔ* (S. Starostin 1995: 216. Preserved in Ket-Yugh and in Kott (as part of a compound). Arin and Pumpokol *tus* ‘salt’ are borrowed from Turkic.

**Proto-Athabaskan**. An unstable item which cannot be reconstructed with certainty.

**Eyak.** *ti:yaʔ*, morphologically unclear, perhaps a new formation.

**Proto-Athabaskan-Eyak**. Not reconstructible.

**Proto-Eskimo**. **taʁəyu-ʁ* (Fortescue et al. 2010: 364), retained in all branches. Cognate to the Aleut term, if not a loan in any direction.

**Proto-Aleut**. **taʁayu-χ* (Bergsland 1994: 384; Bergsland & Dirks 1978: 171; Golovko 1994: 116), attested in Eastern and Atkan; suspiciously close to Proto-Eskimo **taʁəyu-ʁ* ‘salt’, so can be an Eskimo loan. A more frequent Eastern term for ‘salt’ is **alaʁu-χ*, whose Common Aleut meaning is ‘sea, ocean’ (Bergsland 1994: 50; Bergsland & Dirks 1978: 82). It is possible that the concept ‘salt’ should not be reconstructed for Proto-Aleut at all.

**Proto-Chukotian**. Superseded with Russian or Eskimo loanwords.

**Proto-Itelmen**. Cannot be reconstructed with certainty (cf. Fortescue 2005: 387).

**Proto-Nivkh**. **davɬ(-iŋ)* (Fortescue 2016: 41), borrowed from Tungusic lects: Orok *dawsũ*, Evenki *dawasun*, Nanai *daosõ*, all ‘salt’. In their turn, the Tungusic forms represent Mongolic loanwords.

**Proto-Samoyed**. **sər* (Janhunen 1977: 138), the Proto-Samoyed word for ‘ice’, in Northern Samoyed languages also means ‘salt’. Mator and Kamass words for ‘salt’ are borrowed from Turkic, Selkup word for ‘salt’ is apparently an Iranian loan.

**Proto-Yukaghir**. **loɣo-* (Nikolaeva 2006: 246) is attested in several old Yukaghir wordlists. Modern Tundra and Kolyma words for ‘salt’ are Russian borrowings.

**Proto-Burushaski**. **bayu* was borrowed in Balti, Domaaki (as *payu*) and the Shina dialects neighboring Burushaski (as *pažu*, probably due to contamination with *paž-* ‘to cook’), apparently not *vice versa* since *payu* lacks Indo-Aryan etymology and there is an inherited term for ‘salt’ in other Dardic lects (Anton Kogan, p.c.).

105. ‘short’.

**Proto-Yeniseian**. A single candidate is not selectable: Ket-Yugh **pɔʔl* ‘short’ and Kott *tʰuːki* (< **tuk-*?) have more or less equal chances at representing the Proto-Yeniseian item.

**Proto-Athabaskan**. An unstable item. A possible candidate is **=quč’* which is likely to be reconstructed at least for the Proto-Northern level.

**Eyak.** *=tik’*.

**Proto-Athabaskan-Eyak**. **=quč’* (Athabaskan), **=tik’* (Eyak).

**Proto-Eskimo**. **nani-t-* (Fortescue et al. 2010: 233), retained in Yupik and Inuit.

**Proto-Aleut**. **aðu-laka-* (Bergsland 1994: 14; Golovko 1994: 219), attested in all branches. Literally ‘not long’ from **aðu-l-* ‘long’ (q.v.).

**Proto-Chukotian**. **ikmə-* (Fortescue 2005: 95), retained in all languages. Cognate to the Itelmen term.

**Proto-Itelmen**. **ikum-* (Volodin, Khaloimova 2001: 163; Fortescue 2005: 95). Cognate to the Chukotian term.

**Proto-Nivkh**. **bʁaq-* (Fortescue 2016: 25; Savelyeva, Taksami 1965: 188; Savelyeva, Taksami 1970: 298). Morphologically unclear.

**Proto-Samoyed**. **kǝ̆ym* (Janhunen 1977: 51) is retained in all Samoyed languages.

**Proto-Yukaghir**. **monmə-* (Nikolaeva 2006: 275) is attested in Tundra Yukaghir. The Kolyma word *čitnədin-yukoː-* ‘short’ means literally ‘small to long’ (Nikolaeva 2006: 134).

106. ‘snake’.

**Proto-Yeniseian**. Not reconstructible. The original meaning of Proto-Ket-Yugh **čiːk*, considering the external evidence and distribution of cognates, must have been ‘fish’ q.v. Kott-Arin **ʔaŋ-koy* is clearly a composite formation where the second component is **koy* ‘worm’ q.v., and the first one remains unclear.

**Proto-Athabaskan**. **ƛ’eʁeš* is retained in all three branches, morphologically unclear.

**Eyak.** *χuhχ-ʔa-ʔluw-yuː*, literally ‘big worm’, a new formation.

**Proto-Athabaskan-Eyak**. Not reconstructible.

**Proto-Eskimo**. Not reconstructible.

**Proto-Aleut**. Not reconstructible, superseded with loans.

**Proto-Chukotian**. Not reconstructible.

**Proto-Itelmen**. Not reconstructible.

**Proto-Nivkh**. **gəl-ə-ŋa* (Fortescue 2016: 64; Savelyeva, Taksami 1965: 162; Savelyeva, Taksami 1970: 126). Derived from **gəl-* ‘long’ q.v., final **-ŋa* may be **ŋa* ‘animal’ (Fortescue 2016: 117).

**Proto-Samoyed**. **ki-wä* (Janhunen 1977: 72), retained as a word for ‘snake’ only in Selkup, goes back to Proto-Uralic **küyi-wä* ‘snake’ (Aikio 2002: 43–44).

**Proto-Yukaghir**. Not reconstructible.

107. ‘thin’.

**Proto-Yeniseian**. **pakse-m* ‘thin 2D’ (S. Starostin 1995: 245). Preserved in all daughter languages where attested, but not found in Arin or Pumpokol. In Proto-Yeniseian, as in attested languages, the word must have been applied to flat objects. Morphological segmentation of the stem into **pak-si-m* is conditioned by external comparison; Yeniseian-internally, **-m* is indeed a derivational suffix, but **pakse-* (or **paksi-*) functioned as a monolithic stem already in Proto-Yeniseian. Distinct from **tɔq-* ‘thin 1D’ (S. Starostin 1995: 287). Preserved in all daughter languages where attested, but not found in Arin or Pumpokol. In Proto-Yeniseian, as in attested languages, the word must have been applied to 1D-objects.

**Proto-Athabaskan**. The main candidates are **=t’aːn* ‘thin 2D’ (all three branches, can be the same root as PAE **=t’aːn* ‘leaf’) and **=č’eːq’* ‘thin 1D’ (Pacific Coast, Northern).

**Eyak.** *=cʰic-k*.

**Proto-Athabaskan-Eyak**. **=t'aːn*, **=č'eːq'* (Athabaskan), **=cʰiːc* (Eyak). If Nikolaev is correct and PA **=č'eːq'* is to be compared with Eyak *=č'ãːq* 'to be weak' (the correspondence **q' / q* is irregular), the PAE should be reconstructed as **=č'Vnq'*. Leer (2010: 179) compares Eyak *=cʰic-k* with Tlingit *=kʰéxʷ-kʷ* 'light, fluffy', if so the PAET form should be **=kʸʰVkʸ*.

**Proto-Eskimo**. Only tentative reconstruction can be proposed due to inconsistence of the available lexicographic data. Provisionally we fill the slot with two terms. Firstly, **ami-t-* (Fortescue et al. 2010: 26) which means 'to be narrow / to be thin 1D' in Yupik and simply 'to be narrow' in Inuit. Secondly, **caɣə-t-* (Fortescue et al. 2010: 67), which means 'to be thin 2D / to be thin 1D' in Inuit, not attested in Yupik.

**Proto-Aleut**. Reconstruction is based on the Atkan data: **iʁiχ-s-* ~ **iʁiʁi-ða-l-* 'to be thin 1D' (Bergsland 1994: 185; Golovko 1994: 48), **ičaː-qi-ða-l-* 'to be thin 2D' (Bergsland 1994: 170; Golovko 1994: 56). In Eastern, both terms are superseded with negated new formations: *hanatu-laka-* and *aːntuːða-laka-*, both literally mean 'not thick' (Bergsland 1994: 70).

**Proto-Chukotian**. The opposition **ɣət-* 'thin 1D' (Fortescue 2005: 91) / **vəlɣə-* 'thin 2D' (Fortescue 2005: 319) can be safely reconstructed. This system is generally retained in Chukchi, Koryak, Alutor. Chukotian **vəlɣə-* 'thin 2D' is cognate to the Itelmen term with the same meaning.

**Proto-Itelmen**. **kčonʸ-* 'thin 1D' (Volodin, Khaloimova 2001: 36; Fortescue 2005: 395) / **olʸɣʷe-* 'thin 2D' (Volodin, Khaloimova 2001: 67; Fortescue 2005: 319). Itelmen **olʸɣʷe-* 'thin 2D' is cognate to the Chukotian term with the same meaning.

**Proto-Nivkh**. **nok-* 'thin 1D' (Fortescue 2016: 112; Savelyeva, Taksami 1965: 421; Savelyeva, Taksami 1970: 212) is opposed to **hizk-i-la-* 'thin 2D' (Fortescue 2016: 75; Savelyeva, Taksami 1970: 428). In East and South Sakhalin, **nok-* acquires the shape **nozk-* under the influence on the part of **hizk-*.

**Proto-Samoyed**. **yɒptɒ* (Janhunen 1977: 38) is retained in Northern Samoyed, Mator and Selkup.

**Proto-Yukaghir**. Tundra **čöŋkə-*, also attested in Kolyma, but not as the main word for 'thin' (Nikolaeva 2006: 140) vs. Kolyma **keywə-* (Nikolaeva 2006: 204).

108. 'wind'.

**Proto-Yeniseian**. **bey* (S. Starostin 1995: 208). Preserved in all daughter languages.

**Proto-Athabaskan**. **=č̨'ey*, literally 'it blows', is retained in all three branches.

**Eyak.** *k'uːy*, apparently cognate to Athabaskan.

**Proto-Athabaskan-Eyak**. **k'ʷey* (Athabaskan, Eyak).

**Proto-Eskimo**. **anuqə* (Fortescue et al. 2010: 33), retained in Yupik and Inuit.

**Proto-Aleut**. **sla-χ* (Bergsland 1994: 367; Golovko 1994: 189), attested in Eastern and Atkan.

**Proto-Chukotian**. **kətə=yɣ* (Fortescue 2005: 155), attested as a basic term in all languages. Historically 'strong wind' with **kət* 'hard' (Fortescue 2005: 152) and **yəɣə-* 'wind' (Fortescue 2005: 119), the latter root is scarcely retained with the meaning 'wind', although in the majority of cases it has shifted into the meaning 'cool, cold (of weather)'.

**Proto-Itelmen**. **sʸpal* (Volodin, Khaloimova 2001: 140; Fortescue 2005: 400).

**Proto-Nivkh**. **la* (Fortescue 2016: 92; Savelyeva, Taksami 1965: 82; Savelyeva, Taksami 1970: 152).

**Proto-Samoyed**. **märkä* (Janhunen 1977: 93) is retained in all daughter languages save Nganasan.

**Proto-Yukaghir**. **ilʸeyə* (Nikolaeva 2006: 172) is retained in both Yukaghir languages.

109. '(earth)worm'.

**Proto-Yeniseian**. **koy* (S. Starostin 1995: 242). Preserved only in Kott (although the Arin and Pumpokol equivalents are simply not attested). In Ket, replaced in the meaning 'worm' by *ut'iɣ*, a compound of 'snake' with an unclear first component (see notes on the Ket form); in Yugh, replaced by *ɔlli* ‘worm / small insect’, cognate with Ket *ɔləŋgəs* ‘spider’, indicating a more generic term than simply ‘worm’.

**Proto-Athabaskan**. **quːn* is retained in all three branches.

**Eyak.** *χuhχ*.

**Proto-Athabaskan-Eyak**. **quːn* (Athabaskan), **χVnχ* (Eyak; we follow Leer 2008a and reconstruct a medial nasal on account of vowel aspiration in Eyak).

**Proto-Eskimo**. Not reconstructible, ‘earthworm’ is an unstable and poorly documented concept. Cf. Proto-Eskimo **qupəl-ʁuʁ* ‘worm (e.g., in meat), maggot’ (Fortescue et al. 2010: 347) and **quma-ʁ* ‘intestinal worm’ (Fortescue et al. 2010: 344).

**Proto-Aleut**. **iχči-χ* ‘worm (in general)’ (Bergsland 1994: 185; Golovko 1994: 56, 286), attested in Eastern and Atkan.

**Proto-Chukotian**. There are two main candidates. First, **kəmʁə* (Fortescue 2005: 148), meaning ‘worm (generic)’ in Chukchi and apparently Kerek. Second, **ənɣäm* (Fortescue 2005: 343), meaning ‘worm (generic)’ in Koryak and Alutor. The second one seems to be cognate to the Itelmen term, so it has a better chance to represent a Proto-Chukotian term. Nevertheless we prefer to treat both as synonyms.

**Proto-Itelmen**. **xim* (Volodin 1976: 108; Ono 2003: 102; Fortescue 2005: 343), meaning ‘worm (generic)’. Can be cognate to the Chukotian term.

**Proto-Nivkh**. **perŋ* (Fortescue 2016: 133; Savelyeva, Taksami 1970: 284) is a generic term for ‘worm’ and ‘(crawling) insect’. At least in Amur, this seems to be a default expression used for ‘worn’.

**Proto-Samoyed**. **cu̧k ~ *cu̧kə* (Janhunen 1977: 34) is attested as the main word for ‘worm’ and ‘insect’ in Mator and Selkup. Its Northern Samoyed cognates mean rather ‘fly / larva of a fly’. The main word for ‘worm’ in Northern Samoyed, **kə̆lə̆-*, lacks cognates in Southern languages. We list both words as synonyms.

**Proto-Yukaghir**. **könčʸə* (Nikolaeva 2006: 218) is attested in both Yukaghir languages, although its Kolyma reflex is not the main word for ‘worm’ in that language. Another, less likely, candidate is **kelinčʸə* (Nikolaeva 2006: 205), attested only in Kolyma.

110. ‘year’.

**Proto-Yeniseian**. **sɨɢa* (S. Starostin 1995: 275). Preserved in all daughter languages. S. Starostin has proposed that **-ɢa* is an old suffixal element, present also in such words denoting time as **si-ɢ* ‘night’ q.v., **xiʔ-ɢ* ‘day’ (see ‘sun’), but this is questionable.

**Proto-Athabaskan**. **χay* ‘winter / year’ is retained in all three branches.

**Eyak.** *leh q-ʔya*, a descriptive formation based on the preverb *leh* ‘through complete seasonal cycle’ which is the main meaningful morpheme here.

**Proto-Athabaskan-Eyak**. **χay* (Athabaskan).

**Proto-Eskimo**. **ukyu-ʁ* (Fortescue et al. 2010: 397), retained in Yupik and Inuit, polysemy ‘year / winter’.

**Proto-Aleut**. **slu-χ* (Bergsland 1994: 368), attested in all branches, polysemy ‘summer / year’. Also the world **qanax ~ *qanɣi-χ* ‘winter’ can be used with polysemy ‘winter / year’ (Bergsland 1994: 308) in all branches.

**Proto-Chukotian**. **təɣivi* (Fortescue 2005: 292), retained in all languages, except for Kerek. Cognate to the Itelmen term.

**Proto-Itelmen**. **txa-z* (Fortescue 2005: 292). Cognate to the Chukotian term.

**Proto-Nivkh**. **aɲi* (Fortescue 2016: 14; Savelyeva, Taksami 1965: 117; Savelyeva, Taksami 1970: 33). Resembles Proto-Tungusic **aɲŋa-niː* ‘year’, but it can be an accidental similarity, since the stems for ‘year’ in the neighboring Tungusic languages are phonetically far from the Nivkh forms: Orok *anaɲı*, Evenki *anŋaniː*, Nanai *ayŋanı*.

**Proto-Samoyed**. **poə̑* (Janhunen 1977: 127) is retained in all daughter languages.

**Proto-Yukaghir**. Not reconstructible. The Kolyma word *nʸə=molɣil* ‘year’ consists of the reciprocal marker *nʸə-* and the word *molɣil* ‘joint’. Tundra Yukaghir has an extremely polysemous word *sukun* ‘clothes / stuff / ground / sky / weather / year / age / world / life / fact / event’ that can also form a compound with the Tundra word for ‘joint’: *sukun-mol*$^{y}$*ɣal* ‘year / age’.

Berger, Hermann. 1998. *Die Burushaski-Sprache von Hunza und Nager* (Neuindische Studien 13). 3 vols. Wiesbaden: Harrassowitz.

Bergsland, Knut. 1986. Comparative Eskimo-Aleut phonology and lexicon. *Journal de la Société Finno-Ougrienne* 80. 63–137.

Bergsland, Knut. 1989. Comparative aspects of Aleut syntax. *Journal de la Société Finno-Ougrienne* 82. 7–80.

Bergsland, Knut. 1994. *Aleut dictionary*. Fairbanks, Alaska: Alaska Native Language Center, University of Alaska Fairbanks.

Bergsland, Knut. 1997. *Aleut grammar* (Alaska Native Language Center Research Paper no. 10). Fairbanks, Alaska: Alaska Native Language Center, University of Alaska Fairbanks.

Bergsland, Knut & Moses Dirks. 1978. *Qawalangim tunugan kaduuĝingin / Eastern Aleut grammar and lexicon*. Anchorage, Alaska: University of Alaska.

Driem, George van. 2001. *Languages of the Himalayas: an ethnolinguistic handbook of the greater Himalayan region containing an introduction to the symbiotic theory of language* (Handbook of Oriental Studies, Section 2 South Asia 10). 2 vols. Leiden: Brill.

Enrico, John. 2005. *Haida dictionary: Skidegate, Masset, and Alaskan dialects*. 2 vols. Fairbanks, AK: Alaska Native Language Center.

Fortescue, Michael. 1984. *West Greenlandic*. London: Croom Helm.

Fortescue, Michael. 1998. *Language relations across Bering Strait: reappraising the archaeological and linguistic evidence* (Open Linguistics Series). London: Cassell.

Fortescue, Michael. 2003. Diachronic typology and the genealogical unity of Chukotko-Kamchatkan. *Linguistic Typology* 7. 51–88.

Fortescue, Michael. 2005. *Comparative Chukotko-Kamchatkan dictionary*. Berlin: De Gruyter.

Fortescue, Michael. 2016. *Comparative Nivkh dictionary* (Languages of the World. Dictionaries 62). München: LINCOM GmbH.

Fortescue, Michael, Steven Jacobson & Lawrence Kaplan. 2010. *Comparative Eskimo dictionary: with Aleut cognates* (Research Paper / Alaska Native Language Center no. 9). 2nd ed. Fairbanks, AK: Alaska Native Language Center, University of Alaska Fairbanks.

Georg, Stefan. 2008. Yeniseic languages and the Siberian linguistic area. In Alexander Lubotsky, Jos Schaeken, Jeroen Wiedenhof, Rick Derksen & Sjoerd Siebinga (eds.), *Evidence and Counter-Evidence: Essays in Honour of Frederik Kortlandt* (Studies in Slavic and General Linguistics 33), vol. 2, 151–168. Amsterdam: Rodopi.

Georg, Stefan & Aleksandr P. Volodin. 1999. *Die itelmenische Sprache: Grammatik und Texte* (Tunguso-Sibirica 5). Wiesbaden: Harrassowitz.

Golovko, Evgenii V. 1994. *Slovar' aleutsko-russkij i russko-aleutskij (beringovskij dialekt) [Aleut-Russian and Russian-Aleut dictionary (Bering Island dialect)]*. Saint Petersburg: Prosveshchenie.

Golovko, Evgenii V., Nikolai B. Vakhtin & A. S. Asinovskiy. 2009. *Jazyk komandorskikh aleutov. Dialekt ostrova Beringa [Language of the Commander Islands Aleuts. Bering Island dialect]*. Saint Petersburg: Nauka.

Gruzdeva, Ekaterina. 1998. *Nivkh* (Languages of the World 111). München: LINCOM EUROPA.

Helimski, Eugen. 1982. *Drevnejšie vengersko-samodijskie jazykovye paralleli [Ancient linguistic parallels between Hungarian and Samoyed]*. Moscow: Nauka.

Helimski, Eugen. 1997. *Die matorische Sprache : Wörterverzeichnis, Grundzüge der Grammatik, Sprachgeschichte* (Studia Uralo-Altaica 41). Szeged: Department of Altaic Studies, University of Szeged.

Helimski, Eugen. 2005. The 13th Proto-Samoyedic vowel. In Beáta Wagner-Nagy (ed.), *Mikola-konferencia 2004*, 27–39. Szeged: SzTE Department of Finnougristics.

Inenlikei, Petr I. 2005. *Slovar' čukotsko-russkij i russko-čukotskij [Chukchi-Russian and Russian-Chukchi dictionary]*. 3rd ed. Saint Petersburg: Prosveshchenie.

Janhunen, Juha. 1977. *Samojedischer Wortschatz: gemeinsamojedische Etymologien* (Castrenianumin Toimitteita 17). Helsinki: Helsingin yliopisto: Suomalais-ugrilainen seura.

Jochelson, Waldemar. 1926. *The Yukaghir and the Yukaghirized Tungus* (Memoir of the American Museum of Natural History 9). Leiden: Brill / Stechert.

Kari, James. 2010. The concept of geolinguistic conservatism in Na-Dene prehistory. In James Kari, Ben A. Potter & et al. (eds.), *The Dene-Yeniseian connection* (Anthropological Papers of the University of Alaska 5), vol. 1–2, 194–222. Fairbanks, Alaska: University of Alaska Fairbanks.

Kassian, Alexei S. 2011a. Annotated Swadesh wordlists for the Athapaskan group (Na-Dene family). Ongoing project. In George S. Starostin (ed.), *Global lexicostatistical database*. http://starling.rinet.ru/cgi-bin/main.cgi?root=new100.

Kassian, Alexei S. 2011b. Annotated Swadesh wordlists for the Eyak group (Na-Dene family). In George S. Starostin (ed.), *Global lexicostatistical database*. http://starling.rinet.ru/cgi-bin/main.cgi?root=new100.

Kassian, Alexei S. 2011c. Annotated Swadesh wordlists for the Tlingit group (Na-Dene family). In George S. Starostin (ed.), *Global lexicostatistical database*. http://starling.rinet.ru/cgi-bin/main.cgi?root=new100.

Kassian, Alexei S. 2011d. Annotated Swadesh wordlists for the Haida group (Haida family). In George S. Starostin (ed.), *Global lexicostatistical database*. http://starling.rinet.ru/cgi-bin/main.cgi?root=new100.

Kassian, Alexei S., George S. Starostin & Mikhail A. Zhivlov. 2015a. Proto-Indo-European-Uralic comparison from the probabilistic point of view. *Journal of Indo-European Studies* 43(3–4). 301–347.

Kassian, Alexei S., George S. Starostin & Mikhail A. Zhivlov. 2015b. Lexicostatistics, probability, and other matters. *Journal of Indo-European Studies* 43(3–4). 376–392.

Kibrik, Aleksandr E., Sandro V. Kodzasov & Irina A. Muravyeva. 2004. *Language and folklore of the Alutor people* (Endangered Languages of the Pacific Rim A2-042). Kyoto: Nakanishi.

Koryakov, Yuri B. 2018. Problema «jazyk ili dialekt» i samodijskie jazyki [Language vs. dialect question and the Samoyedic languages]. *Ural-Altaic Studies* 31. 156–217.

Krauss, Michael E. 1965. Eyak: a preliminary report. *Canadian Journal of Linguistics* 10. 167–187.

Krauss, Michael E. 1976. Na-Dene. In Thomas A. Sebeok (ed.), *Native Languages of the Americas*, vol. 1, 283–358. New York: Springer Science + Business Media.

Krauss, Michael E. 1977. Proto-Athabaskan-Eyak Fricatives and the First Person Singular. Alaska Native Language Archive, ms. http://www.uaf.edu/anla/item.xml?id=CA961K1977a (1 January, 2019).
Krauss, Michael E. & Jeff Leer. 1981. *Athabaskan, Eyak, And Tlingit sonorants* (Alaska Native Language Center Research Papers 5). Fairbanks, Alaska: University of Alaska Fairbanks.
Kurebito, Tokusu, Megumi Kurebito, Yukari Nagayama, Chikako Ono & Mitsuhiro Yazu. 2001. *Comparative basic vocabulary of the Chukchee-Kamchatkan language family* (Endangered Languages of the Pacific Rim A2-011). Vol. 1. Suita, Japan: Faculty of Informatics, Osaka Gakuin University.
Kurilov, Gavril N. 2001. *Jukagirsko-russkij slovar' [Yukaghir-Russian dictionary]*. Novosibirsk: Nauka.
Leer, Jeff. 1979. *A conversational dictionary of Kodiak Alutiiq*. Fairbanks, Alaska: University of Alaska Fairbanks.
Leer, Jeff. 1996. Comparative Athabaskan lexicon. Alaska Native Language Archive, ms. http://www.uaf.edu/anla/item.xml?id=CA965L1996a (1 January, 2019).
Leer, Jeff. 2008a. Proto-Athabascan nasal codas and Eyak aspirated vowels. In Tokusu Kurebito (ed.), *Linguistic typology of the North*, vol. 1, 1–16. Tokyo: Research Institute for Languages and Cultures of Asia and Africa, Tokyo University of Foreign Studies.
Leer, Jeff. 2008b. Recent advances in AET comparison. Handout presented at the Dene-Yeniseian Symposium, Fairbanks, AK. https://www.uaf.edu/files/anlc/leer-2008.pdf (28 November, 2018).
Leer, Jeff. 2010. The palatal series in Athabascan-Eyak-Tlingit, with an overview of the basic sound correspondences. In James Kari, Ben A. Potter & et al. (eds.), *The Dene-Yeniseian connection* (Anthropological Papers of the University of Alaska 5), vol. 1–2, 168–193. Fairbanks, Alaska: University of Alaska Fairbanks.
MacLean, Edna Ahgeak. 2014. *Iñupiaq to English dictionary*. Fairbanks, Alaska: University of Alaska Press.
Menovshchikov, Georgiy A. 1964. *Jazyk sirenikskix èskimosov (fonetika, očerk morfologii, teksty i slovar') [Language of the Sirenik Eskimos. Phonology, morphology sketch, texts, dictionary]*. Moscow: Nauka.
Menovshchikov, Georgiy A. 1988. *Slovar' èskimossko-russkij i russko-èskimosskij [Eskimo-Russian and Russian-Eskimo dictionary]*. Leningrad: Prosveshchenie.
Moll, Tatiana A. & Petr I. Inenlikei. 1957. *Čukotsko-russkij slovar' [Chukchi-Russian dictionary]*. Leningrad: Ministerstvo prosvescheniya.
Mudrak, Oleg A. 2000. *Ètimologičeskij slovar' čukotsko-kamčatskix jazykov [Etymological dictionary of the Chukotko-Kamchatkan languages]*. Moscow: Yazyki russkoy kultury.
Mudrak, Oleg A. 2005. Itelmen etymology. In Sergei A. Starostin (ed.), *The Tower of Babel. An etymological database project*. http://starling.rinet.ru/cgi-bin/main.cgi.
Mudrak, Oleg A. 2008. *Svod kamčadal'skoj leksiki po pamjatnikam XVIII veka [Kamchatkan lexicon according to the 18th century sources]*. Moscow: Vostochnaya Literatura.
Nagayama, Yukari. 2003. *Očerk grammatiki aljutorskogo jazyka [Grammatical outline of Alutor]* (Endangered Languages of the Pacific Rim A2-038). Kyoto: Nakanishi.
Nikolaev, Sergei L. 2014. Toward the reconstruction of Proto-Na-Dene. *Journal of Language Relationship* 11. 103–123.
Nikolaeva, Irina A. 2006. *A historical dictionary of Yukaghir* (Trends in Linguistics 25). Berlin: Mouton de Gruyter.
Ono, Chikako. 2003. *A lexicon of words and conversational phrases for the Itelmen Northern dialect* (Endangered Languages of the Pacific Rim A2-029). Kyoto: Nakanishi.
Oswalt, Robert. 1970. The detection of remote linguistic relationships. *Computer Studies in the Humanities and Verbal Behavior* 3. 117–129.
Panfilov, Valery Z. 1962. *Grammatika nivxskogo jazyka [Grammar of the Nivkh language]*. Vol. 1. 2 vols. Moscow: AN SSSR.

Panfilov, Valery Z. 1965. *Grammatika nivxskogo jazyka [Grammar of the Nivkh language]*. Vol. 2. 2 vols. Moscow: AN SSSR.
Peiros, Ilya I. & Sergei A. Starostin. 1986. O tonax nivxskogo jazyka [On Nivkh tones]. In Petr Ya. Skorik (ed.), *Paleoaziatskie jazyki [Paleoasiatic languages]*, 213–218. Leningrad: Nauka.
Pozdniakov, Konstantin. 2014. On the threshold of relationship and the "stability index" of basic lexicon in mass comparison: Atlantic languages. *Journal of Language Relationship* 11. 187–210.
Savelyeva, Valentina N. & Chuner M. Taksami. 1965. *Russko-nivxskij slovar' [Russian-Nivkh dictionary]*. Moscow: Sovetskaya entsiklopediya.
Savelyeva, Valentina N. & Chuner M. Taksami. 1970. *Nivxsko-russkij slovar' [Nivkh-Russian dictionary]*. Moscow: Sovetskaya entsiklopediya.
Skorik, Petr Ya. 1958. K voprosu o klassifikacii čukotsko-kamčatskix jazykov [On the classification of the Chukotko-Kamchatkan languages]. *Voprosy Jazykoznanija* 1. 21–35.
Skorik, Petr Ya. 1961. *Grammatika čukotskogo jazyka. Č. 1. Fonetika i morfologija imennyx častej reči [Chukchi grammar. Part 1. Phonology, nominal morphology]*. Moscow: AN SSSR.
Skorik, Petr Ya. 1977. *Grammatika čukotskogo jazyka. Č. 2. Glagol. Narečija. Služebnye slova [Chukchi grammar. Part 2. Verb, adverbs, auxiliary words]*. Leningrad: Nauka.
Starostin, George S. 2012. Dene-Yeniseian: a critical assessment. *Journal of Language Relationship* 8. 117–138.
Starostin, George S. 2013a. Annotated Swadesh wordlists for the Yeniseian group (Yeniseian family). In George S. Starostin (ed.), *Global lexicostatistical database*. http://starling.rinet.ru/cgi-bin/main.cgi?root=new100.
Starostin, George S. 2013b. Annotated Swadesh wordlists for the Burushaski group (Burushaski family). In George S. Starostin (ed.), *Global lexicostatistical database*. http://starling.rinet.ru/cgi-bin/main.cgi?root=new100.
Starostin, Sergei A. 1995. Sravnitel'nyj slovar' enisejskix jazykov [Comparative dictionary of the Yeniseian languages]. In Sergei A. Starostin (ed.), *Ketskij sbornik: lingvistika / Studia ketica: Linguistics*, 176–315. Moscow: Vostochnaya literatura.
Starostin, Sergei A. 2005a. Yenisseian etymology. In Sergei A. Starostin (ed.), *The Tower of Babel. An etymological database project*. http://starling.rinet.ru/cgi-bin/main.cgi.
Starostin, Sergei A. 2005b. Burushaski etymology. In Sergei A. Starostin (ed.), *The Tower of Babel. An etymological database project*. http://starling.rinet.ru/cgi-bin/main.cgi.
Starostin, Sergei A. 2007a. Opredelenie ustojčivosti bazisnoj leksiki [Defining the stability of basic lexicon]. *Trudy po jazykoznaniju [Works on linguistics]*, 827–839. Moscow: Jazyki slavjanskix kul'tur.
Starostin, Sergei A. 2007b. Praenisejskaja rekonstrukcija i vnešnie svjazi enisejskix jazykov [Proto-Yeniseian reconstruction and the external connections of the Yeniseian languages]. *Trudy po jazykoznaniju [Works on linguistics]*, 147–246. Moscow: Jazyki slavjanskix kul'tur.
Stebnitsky, S.N. 1934. Itel'menskij jazyk [Itelmen language]. In E.A. Kreinovich (ed.), *Jazyki i pis'mennost' narodov severa. Č. 3. Jazyki i pis'mennost' paleoaziatskix narodov [The language and literature of the people of the North. P. 3. The language and literature of the Paleoasiatic peoples]*, 85–104. Moscow: GUPI.
Taksami, Chuner M. 1996. *Slovar' nivxsko-russkij i russko-nivxskij [Nivkh-Russian and Russian-Nivkh dictionary]*. 2nd ed. Saint Petersburg: Prosveshchenie.
Thomason, Sarah Grey & Terrence Kaufman. 1988. *Language contact, creolization, and genetic linguistics*. Berkeley: University of California Press.
Vajda, Edward J. 2001. *Yeniseian peoples and languages: a history of Yeniseian studies with an annotated bibliography and a source guide*. Richmond: Curzon.
Volodin, Aleksandr P. 1976. *Itel'menskij jazyk [Itelmen language]*. Leningrad: Prosveshchenie.

Volodin, Aleksandr P. 1997. Čukotsko-kamčatskie jazyki [Chukotko-Kamchatkan languages]. In Aleksandr P. Volodin (ed.), *Jazyki mira: paleoaziatskie jazyki [Languages of the World: Paleoasiatic languages]*, 12–22. Moscow: Indrik.

Volodin, Aleksandr P. & Klavdia N. Khaloimova. 2001. *Slovar' itel'mensko-russkij i russko-itel'menskij [Itelmen-Russian and Russian-Itelmen dictionary]*. 2nd ed. Saint Petersburg: Prosveshchenie.

Werner, Heinrich. 2002. *Vergleichendes Wörterbuch der Jenissej-Sprachen*. 3 vols. Wiesbaden: Harrasowitz.

Worth, Dean S. 1962. La place du kamtchadal parmi les langues soi-disant paleosiberiennes. *Orbis* 11. 579–599.

Zhukova, Alevtina N. 1967. *Russko-korjakskij slovar' [Russian-Koryak dictionary]*. Moscow: Sovetskaya entsiklopediya.

Zhukova, Alevtina N. 1972. *Grammatika korjakskogo jazyka. Fonetika i morfologija [Koryak grammar: phonology, morphology]*. Leningrad: Nauka.

Zhukova, Alevtina N. 1990. *Slovar' korjaksko-russkij i russko-korjakskij [Russian-Koryak and Koryak-Russian dictionary]*. 2nd ed. Leningrad: Prosveschenie.
