## Supplementary figures and images for "Circumpolar peoples and their languages: lexical and genomic data suggest ancient Chukotko-Kamchatkan–Nivkh and Yukaghir-Samoyedic connections"

### Supplementary Figure 1

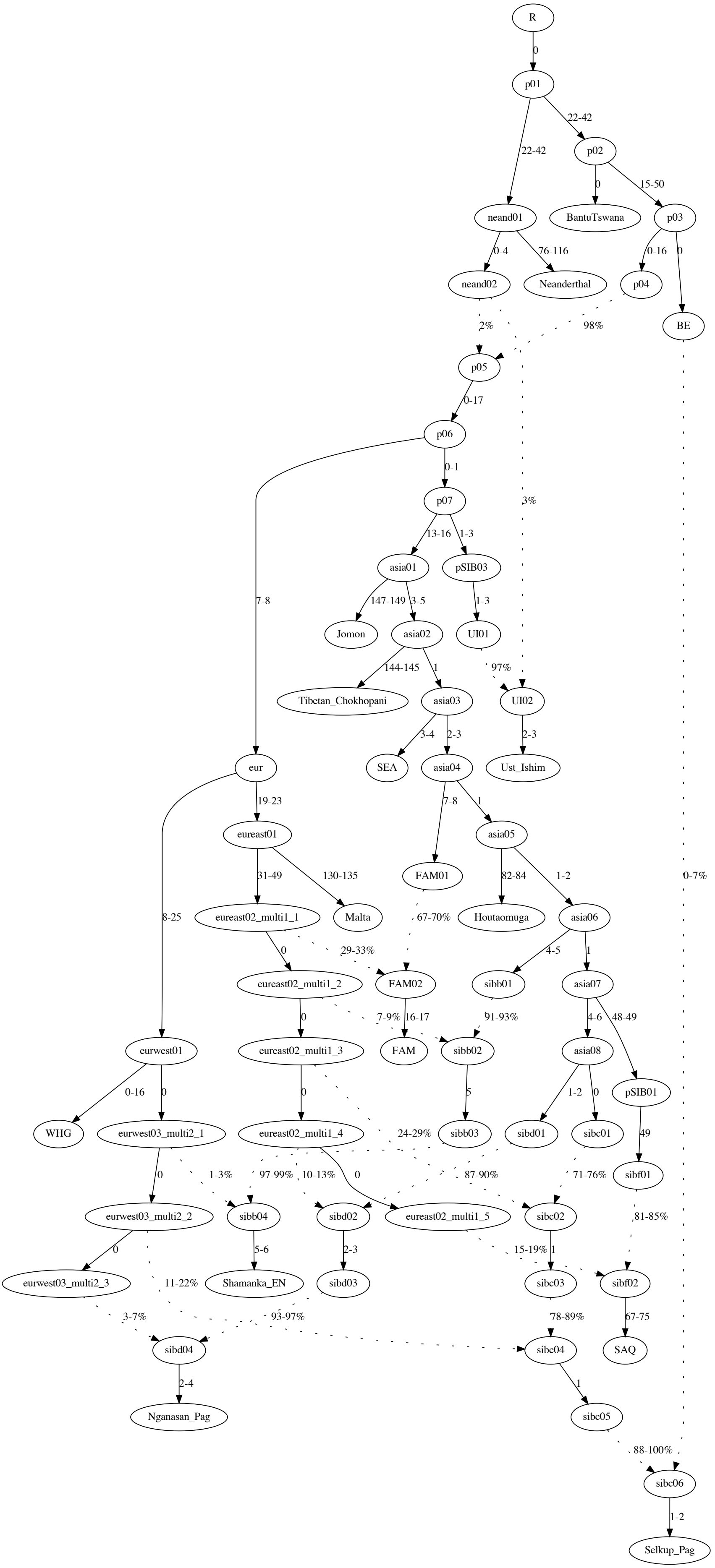

### Supplementary Figure 2

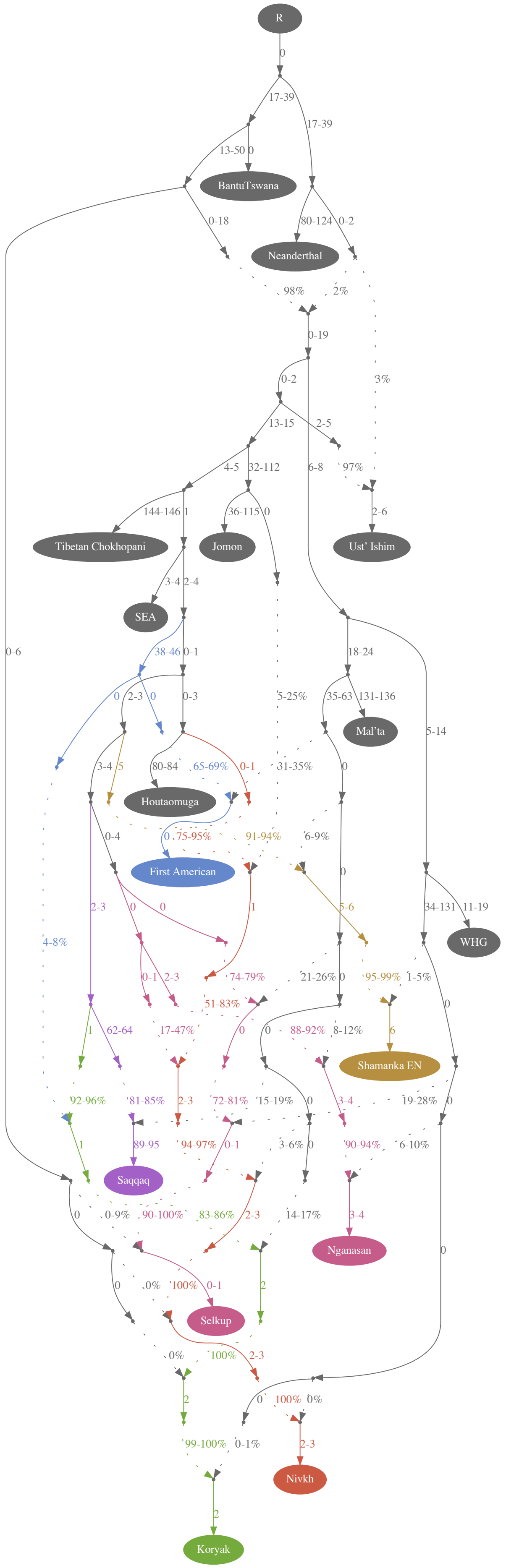
